# Multivariate pattern analysis identifies potential inter-trial resting-state EEG biomarkers in fibromyalgia

**DOI:** 10.1101/2025.05.30.657056

**Authors:** Dino Soldic, María Carmen Martín-Buro, David López-García, Ana Belén del Pino, Roberto Fernandes-Magalhaes, David Ferrera, Irene Peláez, Luis Carretié, Francisco Mercado

## Abstract

Fibromyalgia involves widespread musculoskeletal pain and hypersensitivity, often accompanied by neurological, cognitive, and affective disturbances. Resting-state electroencephalography studies have revealed abnormal brain activity in chronic pain conditions, with anxiety and symptom duration potentially exacerbating these alterations. This study applied multivariate pattern analysis to differentiate inter-trial resting-state electroencephalography signals between fibromyalgia patients and healthy controls across frequency bands associated with pain processing, incorporating state and trait anxiety scores. It also examined differences between patients with short and long duration symptoms and identified the most relevant scalp regions contributing to the models. Fifty-one female participants (25 fibromyalgia patients, 26 controls; aged 35-65) were included. Patients were classified into short-term (12) and long-term (13) groups. Normalized power spectral density values were extracted from electroencephalography data and used to train machine learning classifiers, with Haufe-transformed weights computed to determine key scalp contributions. The models distinguished patients from controls with area under the curve values exceeding 0.75 across all frequency bands, reaching 0.99 in beta and gamma bands when anxiety was included. Symptom duration was also a relevant factor, as the model differentiated short-from long-term fibromyalgia patients with area under the curve values up to 0.96 in beta and gamma bands. Alterations in theta power within frontal and parietal regions, along with frequency-specific contributions, highlight disrupted pain processing in fibromyalgia and suggest cumulative effects of prolonged symptom duration. Future resting-state studies leveraging multivariate pattern analysis may support the development of potential biomarkers to improve diagnosis and guide treatment strategies in clinical settings.

## 1 Introduction

Fibromyalgia (FM) affects 2-8% of the population, predominantly women (Clauw, 2014). The syndrome is characterized by widespread musculoskeletal pain and hypersensitivity, also affecting neurological, cognitive, and affective domains (Fallon et al., 2018; Ferrera et al., 2020, 2021; Giorgi et al., 2022; Pujol et al., 2014, 2022). As a nociplastic pain syndrome (Clauw, 2024), FM presents a controversial neurobiological basis and highly variable comorbid symptoms (e.g., fatigue and insomnia). Since current clinical diagnosis relies heavily on subjective self-reports and tender point assessments, FM is associated with high misdiagnosis rates and significant diagnostic delays, often taking several years for patients to receive an accurate diagnosis (Fitzcharles et al., 2021; Kumbhare et al., 2018; Wolfe et al., 2010, 2016). These challenges highlight the clinical urgency for objective electrophysiological biomarkers and have prompted researchers to record electromagnetic brain signals to better understand its underlying mechanisms. Electroencephalography (EEG) is especially useful, as it can detect abnormalities in basal brain function even during resting-state (RS) conditions. In this line, anomalous cortical activity in FM has been shown to be more concentrated in parietal regions within theta, alpha, and beta bands (Martín-Brufau et al., 2021). Additionally, Fallon et al. (2018) observed increased theta activity associated with heightened pain perception, localized via source reconstruction to the anterior cingulate, medial prefrontal, and dorsolateral prefrontal cortices. Other studies report diminished connectivity between default mode network regions and somatosensory cortices in the theta band (Choe et al., 2018; Hsiao et al., 2017), a pattern linked to hypersensitive neural processing in FM (Martín-Brufau et al., 2021). Complementing these findings, González-Villar et al. (2020) observed increased alpha and beta activity in frontal regions, possibly reflecting dysregulated top-down control contributing to FM patients’ attentional and cognitive deficits (Fernandes-Magalhaes et al., 2022, 2024). While most RS-EEG studies use continuous recordings, some recent work has leveraged short inter-trial epochs to probe residual task-related dynamics (D’Croz-Baron et al., 2021; Murphy et al., 2018). These intervals are not equivalent to conventional RS-EEG but can serve as a practical window into cortical activity following task engagement.

Among the factors that may influence these RS alterations, symptom duration is particularly relevant. Continuous nociceptive input, characteristic of chronic pain conditions, appears to alter processing in cortical regions (Apkarian et al., 2004; Bekkelund et al., 1995; Jin et al., 2013; Rodriguez-Raecke et al., 2013), reduce deactivations, and disrupt inhibitory interactions between the dorsolateral and medial prefrontal cortices (Baliki et al., 2008). Similar patterns have been observed in FM, with several studies linking symptom duration to increased theta power in frontal and parietal regions, suggesting that prolonged exposure to pain and leads to progressive alterations in FM (Villafaina, Collado-Mateo, Fuentes-García, et al., 2019). Furthermore, frontocentral activity changes associated with ongoing tonic pain and fatigue support the notion of a cumulative neural impact with prolonged FM exposure (Alves et al., 2023; Fallon et al., 2018). However, chronic pain is not only a defining feature of FM but may also be influenced by other comorbid symptoms, particularly affective factors (Henao-Pérez et al., 2022). Anxiety is among the most prevalent affective symptoms in FM (Galvez-Sánchez et al., 2020) and often leads to a marked reduction in quality of life (Catalá et al., 2023; Cetingok et al., 2022). Rather than a mere comorbidity, anxiety acts as a critical regulator of pain perception through maladaptive top-down modulation (Bushnell et al., 2013; Garcia-Larrea & Bastuji, 2018). Consequently, incorporating anxiety into neurophysiological models allows for a transition from identifying pure nociceptive signaling toward characterizing affectively modulated pain phenotypes (Ploghaus et al., 2001), reflecting the reality that the neural signature of FM is inextricably linked to the patient’s affective state (Kim et al., 2015). This link is further supported by findings associating trait anxiety with disrupted activity within the default mode network and state anxiety with alterations in the salience network (Saviola et al., 2020).

Given the syndrome’s variability and complexity, machine learning (ML) methods have gained popularity for their ability to improve statistical sensitivity and specificity by detecting subtle signal changes often overlooked by conventional analyses, thereby preserving valuable information (Maris & Oostenveld, 2007; Thanh Nhu et al., 2022; Zafar et al., 2018). In contrast to traditional univariate approaches that assess power changes in isolated EEG channels, multivariate pattern analysis (MVPA) considers the entire scalp topography simultaneously (López-García et al., 2022). This approach reflects network level disruptions rather than localized alterations, making it particularly suited for capturing the high dimensional nature of FM. For instance, López-Solà et al. (2017) identified new cortical regions involved in FM using MVPA of functional magnetic resonance imaging data. Furthermore, Hsiao et al. (2021) employed MVPA to predict individual differences in heat pain sensitivity, combining RS-EEG features such as power spectral density, functional connectivity, and psychometric metrics. Their study demonstrated that combining neurophysiological and neuropsychological variables enhances predictive performance, highlighting the utility of ML in uncovering underlying neural mechanisms based on RS-EEG features (Abdellatef et al., 2023).

Despite their potential, such approaches have yet to be systematically applied to FM populations, highlighting the need for methods that deepen our understanding of FM’s neurobiological foundations and reliably differentiate patients from HC, while accounting for clinical symptoms. Therefore, this study aims to: 1) differentiate inter-trial resting-state (itRS)-EEG signals of FM patients and HC across frequency bands leveraging state and trait anxiety scores; 2) distinguish patients based on symptom duration; and 3) identify neural activity in key scalp regions contributing to classification. While these signals were extracted from task-free intervals rather than conventional RS recordings, this itRS approach allows for the investigation of background neural activity within a controlled cognitive context.

## 2 Materials and methods

### 2.1 Paradigm and data acquisition

#### 2.1.1 Participants

Fifty-one women participated in the study, comprising 25 fibromyalgia (FM) patients (mean age ± SD = 52.15 ± 6.06) and 26 healthy controls (HC; mean age ± standard deviation [SD] = 48.73 ± 7.01), with ages ranging from 35 to 65 years. FM patients were recruited through the Fibromyalgia and Chronic Fatigue Syndrome Association of Móstoles (AFINSYFACRO) and the Madrid Fibromyalgia Association (AFIBROM). They were evaluated by rheumatologists from hospitals in the Community of Madrid and met the diagnostic criteria defined by the American College of Rheumatology (Wolfe et al., 1990, 2010). HC were recruited through posters displayed around the Faculty of Health Sciences at Rey Juan Carlos University, as well as through emails sent to the entire university community. Participants with neurological disorders, metabolic alterations, cognitive impairments, substance abuse or dependence, or psychiatric disorders were excluded from the study. FM patients had a mean diagnosis duration of 94 months (SD = 50.77). Prescribed medications and therapeutic interventions were continued due to medical recommendations and ethical guidelines. Group differences in age and anxiety scores between FM and HC participants were assessed using two-tailed independent-samples *t*-tests with a 95% confidence interval, while differences in educational level were evaluated using a chi-square (*χ²*) test.

#### 2.1.2 Procedure

The study adhered to the ethical standards of the World Medical Association’s Helsinki Declaration (1964) and its later amendments. All participants provided written informed consent, and the study was approved by the Rey Juan Carlos University Research Ethics Committee (ref: 201000100011550).

Upon arrival at the Laboratory of Cognitive Neuroscience (Faculty of Health Sciences, Rey Juan Carlos University), participants completed a sociodemographic and clinical questionnaire. Sociodemographic data included age, educational level, and regular medication use. Additionally, FM patients reported the duration of their diagnosis in months. FM and HC groups were matched for age [*t* (49) = 1.625, *p* = 0.111, *d* = 0.455] and educational level [*χ^2^*(4) = 8.106, *p* = 0.088, *V* = 0.399]. To assess both state (STAI-S) and trait (STAI-T) anxiety the State-Trait Anxiety Inventory (STAI) (Spielberger et al., 1982) was administered. Sociodemographic and clinical data are presented in Table 1.

**Table 1.** Sociodemographic and clinical data.

|  | **Fibromyalgia**<br>**(N = 25)** | **Healthy Control**<br>**(N = 26)** | **T-test/**<br>**chi square** | **Significance**<br>**level** | **Effect size** |
| --- | --- | --- | --- | --- | --- |
| **Age (years. M ± SD)** | 51.60 ± 5.47 | 48.73 ± 7.01 | $t = 1.625$ | $p = 0.111$ | $d = 0.455$ |
| **Educational level** | | | $\chi^2 = 8.106$ | $p = 0.088$ | $V = 0.399$ |
| None, N (%) | 3 (5.9%) | 0 (0%) | - | - | - |
| Primary school, N (%) | 11 (21.6%) | 8 (15.7%) | - | - | - |
| Secondary school, N (%) | 7 (13.7%) | 6 (11.8%) | - | - | - |
| High school, N (%) | 3 (5.9%) | 11 (21.6%) | - | - | - |
| Bachelor's degree, N (%) | 1 (2%) | 1 (2%) | - | - | - |
| **Diagnosis history (months.**<br>**M ± SD)** | 94 ± 50.77 | - | - | - | - |
| **Diagnosis history (months.**<br>**Median)** | 92 | - | - | - | - |
| **Medication** |  |  |  |  |  |
| Antidepressants, N (%) | 14 (56%) | 0 (0%) | - | - | - |
| Anxiolytics, N (%) | 6 (24%) | 0 (0%) | - | - | - |
| Analgesics, N (%) | 10 (40%) | 0 (0%) | - | - | - |
| Antiepileptics/Psycholeptics,<br>N (%) | 4 (16%) | 0 (0%) | - | - | - |
| Other, N (%) | 9 (36%) | 0 (0%) | - | - | - |
| **Trait anxiety (STAI-T**<br>**percentile. M ± SD)** | 81.94 ± 19.33 | 29.73 ± 27.96 | $t = 7.781$ | $p < 0.001$ | $d = 2.172$ |
| **State anxiety (STAI-S**<br>**percentile. M ± SD)** | 60.52 ± 26.48 | 15.77 ± 18.94 | $t = 6.918$ | $p < 0.001$ | $d = 1.944$ |

Abbreviations: N = Sample size; M = Mean; SD = Standard deviation.

#### 2.1.3 EEG data acquisition and preprocessing

EEG recording took place in a room fully equipped to minimize noise, electromagnetic interference, and environmental influences, providing an optimal setting for the procedure. Continuous EEG activity was recorded from 60 homogeneously distributed electrodes (10-10 system) mounted on a cap (ElectroCap International), referenced to both mastoids. Vertical and horizontal eye movements were monitored through electrooculographic (EOG) recordings. The vertical EOG electrodes were positioned both below and above the left eye orbit, while the horizontal EOG electrodes were situated on the left and right orbital rims. Electrode impedances were maintained below 5 kΩ, and a bandpass filter of 0.1 – 50 Hz (3 dB points for -6 dB/octave roll-off) was applied to the recording amplifiers. Data were continuously digitized at a sampling rate of 250 Hz. Offline preprocessing was performed using a custom MATLAB R2023b (The MathWorks, Inc., 2023) script containing EEGLAB v2023.1 (Delorme & Makeig, 2004) functions.

The EEG dataset used in this study was recorded as a part of a broader investigation involving event-related potentials (ERP), which falls beyond the scope of this article. Consequently, we utilized an itRS design, extracting data from the inter-trial intervals (ITI) to represent background neural activity. In the original experiment, participants performed a simple visual task that required categorizing a central digit as either above or below five, while peripheral stimuli were simultaneously presented (see, for example, Carretié (2014)). Visual stimuli were displayed for 50 ms, with an inter-stimulus interval (ITI) of 3000 ms. During the ITI, participants stared at a fixation cross without engaging in any other activity. Previous studies described that ERPs can last up to 900 ms post-stimulus onset gradually returning to baseline cortical activity (Iemi et al., 2019). Meanwhile, during attentional tasks, activity in the default mode network increases greatly around 600 ms upon stimulus onset and around 200 ms after the response (Walz et al., 2014), reflecting a transition to RS brain activity (Raichle & Snyder, 2007). Thus, to ensure EEG segments were free from task-related activity, the maximum response time across all participants (850 ms) was used as a baseline, with an additional 500 ms safety added window. Epochs were then defined from 1350 ms after stimulus onset to 300 ms before the next stimulus onset. The resulting 1350 ms itRS epoch was chosen to maximize the likelihood that the signal reflects a return to baseline while acknowledging that, unlike conventional RS-EEG, these segments may be influenced by the cognitive set of the preceding task. This resulted in a total of 4893 itRS epochs with eyes open, each lasting 1350 ms, as per previous studies (D’Croz-Baron et al., 2021; Fallon et al., 2018). Prior to epoching, ocular channels were removed from the data, and the entire EEG recordings were digitally filtered using a zero-phase finite impulse response (FIR) filter implemented in EEGLAB’s *pop_eegfiltnew* function (0.5-50 Hz), with default filter order and transition bandwidth parameters. Eye blink artifacts were removed from the epoched data using Independent Component Analysis through EEGLAB’s *runica* algorithm. Out of 60 total components, an average of 1 component (*SD* = 0.4) was removed for the HC group and 1.31 components (*SD* = 0.61) were removed for the FM group. Electrodes exhibiting persistent high frequency noise or poor contact were interpolated using spherical splines. To prune epochs containing artifacts, a semi-automatic trial rejection was performed based on voltage thresholds set at ± 100 µV. To further ensure there was no lingering task activity in the data, epochs that still contained a response event (i.e., stimulus trigger) were manually selected and removed. Additionally, all remaining epochs underwent a final visual inspection to identify and exclude any residual non-biological artifacts (e.g., electrode drifts or abrupt muscle bursts) not captured by the automated threshold. As a result, 2351 epochs (mean = 94.04; *SD* = 1.89) for the FM group and 2455 epochs (mean = 94.42; *SD* = 2.86) for the HC group were retained for subsequent analyses. After artifact removal the data were re-referenced to the common average reference. The MATLAB script used for preprocessing is available at: https://github.com/dinosoldic/EEG-Preproc-ERP/blob/main/eegPreproc.m. A summary of the EEG analysis pipeline is presented in Figure 1.

**Figure 1.**
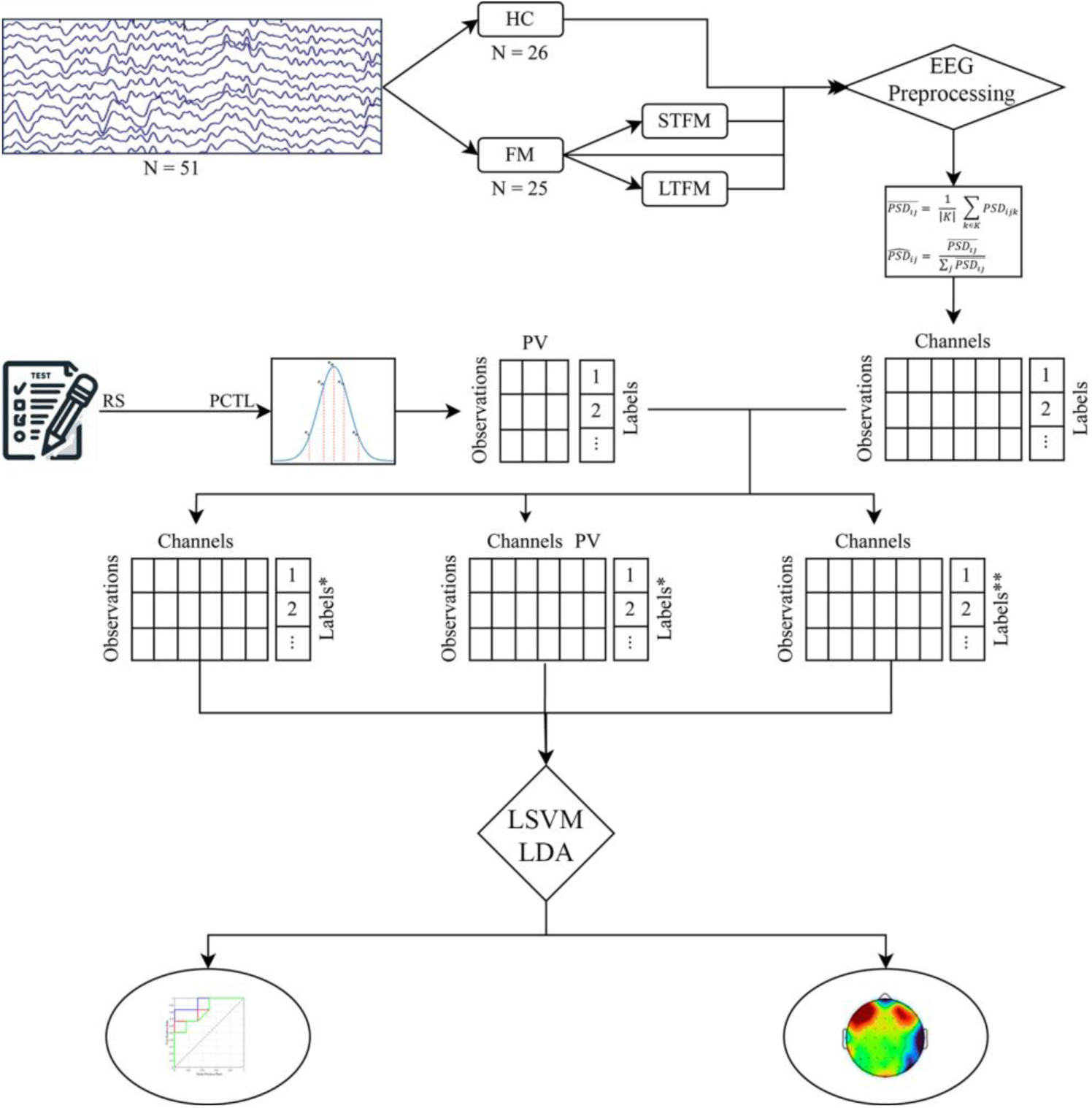
Pipeline of data processing and analyses. (*) indicates labels used for FM and HC comparison. (**) indicates labels used for STFM and LTFM comparison. Abbreviations: N = Sample size; HC = Healthy controls; FM = Fibromyalgia; STFM = Short-term fibromyalgia; LTFM = Long-term fibromyalgia; EEG = Electroencephalography; RS = Raw scores; PCTL = Percentile; PV = Psychometric values; LSVM = Linear support vector machine; LDA = Linear discriminant analysis.

### 2.2 Feature extraction

Building on previous studies examining RS data and patient diagnosis (Abdellatef et al., 2023; Hsiao et al., 2021; Ruiz De Miras et al., 2023), the EEG data was decomposed into frequency bands. The resulting bands were as follows: delta (2 - 3.5 Hz), theta (4 - 7.5 Hz), alpha (8 - 12 Hz), alpha-1 (8 - 10 Hz), alpha-2 (10 - 12 Hz), beta (13 - 30 Hz), beta-1 (13 - 18 Hz), beta-2 (18.5 - 21 Hz), beta-3 (21.5 - 30 Hz), and gamma (30.5 - 44 Hz) (Fallon et al., 2018; Villafaina, Collado-Mateo, & Fuentes-García, 2019). The inclusion of both broad bands and their respective sub-bands was intended as an exploratory multiscale spectral analysis. This approach allowed us to evaluate whether discriminative power was distributed across the entire frequency range or localized within specific spectral components, providing a more detailed characterization of the FM neural signature that might be obscured by traditional broad band definitions. These frequency bands were obtained using the *ft_freqanalysis* function in FieldTrip (Oostenveld et al., 2011), applying the multitaper method for fast Fourier transform (mtmfft) with discrete prolate spheroid sequences on frequencies of interest ranging from 0.5 to 50 Hz, spaced at 0.74 Hz.

Once extracted, power spectral density (PSD) values were averaged across bins within each frequency band (Equation 1), and normalized (Equation 2) by dividing the PSD value at each channel by the total power of its corresponding observation (Hsiao et al., 2021). This procedure was applied separately to each frequency band.

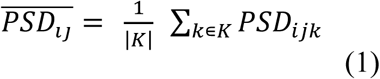

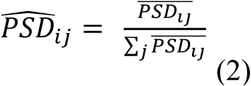

In Equation 1, 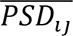 represents the averaged power for observation *i* at channel *j*. The term *K* denotes the set of frequency bins within a specific frequency band, *|K|* is the total number of bins in that set, and *PSD*_*ijk*_ is the power value at the *k*-th frequency bin. In Equation 2, 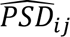 represents the normalized power for observation *i* at channel *j*, where the denominator serves as a scaling factor representing the total power across all *j* (where *j* = 1, … ,60) channels for that specific observation.

In addition to the frequency bands, psychometric values (PV) were extracted from self-reported anxiety scores obtained through the STAI questionnaire. These values consisted of the percentile scores for each participant on both the trait and state anxiety scales. These features were selected a priori due to the high prevalence of anxiety in FM populations and its known impact on neural processing (Galvez-Sánchez et al., 2020; Henao-Pérez et al., 2022). The inclusion of these psychometric measures was a secondary objective designed to complement the primary neurophysiological analysis. PV were added as features along with PSD to examine their potential contribution to classification performance and to assess whether anxiety-related measures provide additional discriminative power between groups. By modeling these features together, we aimed to determine the incremental validity of neurophysiological markers, specifically, whether EEG features capture unique variance beyond what is provided by subjective clinical scales. This integration allows for a characterization of the FM neuro-affective phenotype, testing the hypothesis that brain oscillations offer objective discriminative information that complements, rather than simply replicates, psychometric data. Since PSD and PV are measured in different units, they were normalized using the *zscore* function in MATLAB. Consequently, the training and testing datasets for comparing HC and FM were constructed using PSD, PV (STAI-S, STAI-T, and STAI-S + STAI-T), as well as combinations of PSD and PV (PSD + STAI-S, PSD + STAI-T, and PSD + STAI-S + STAI-T). As a result, the input data for the models contained as observations (rows) the full number of epochs (4,806) and a variable number of columns representing the features. Specifically, the PSD only models (one per frequency band) utilized 60 columns corresponding to the EEG channels, the PV only models used 1 or 2 columns for the anxiety scores, and the combination models utilized 61 or 62 columns by adding the psychometric scores as extra feature columns to the PSD set. In the combination models, the individual psychometric score for each participant was replicated across all of their corresponding EEG epochs.

Additionally, the FM group was divided into two subgroups based on diagnostic length: 12 patients were included in the short-term fibromyalgia (STFM) group and 13 in the long-term fibromyalgia (LTFM) group. Based on previous analyses in FM research (Amris et al., 2016; Villafaina, Collado-Mateo, Fuentes-García, et al., 2019), the median diagnostic length (92 months, *Q1* = 64, *Q3* = 107) was used as the cutoff, with participants at the median value being randomly assigned to either group to ensure balance. The STFM group had a mean diagnostic duration of 61 months (*SD* = 16.36), while the LTFM group had a mean of 124 months (*SD* = 52.75). Consequently, the input data for this specific comparison contained 2,351 observations corresponding to epochs as rows and 60 columns corresponding to the EEG channels as features, per frequency band. Although dichotomizing a continuous variable can reduce information in traditional statistical analyses (MacCallum et al., 2002), we used a median split here to define two groups for supervised classification. This choice was theoretically motivated by evidence suggesting that chronic pain leads to progressive, stage-like alterations in neural architecture (Alves et al., 2023; Villafaina, Collado-Mateo, Fuentes-García, et al., 2019). By employing a classification framework rather than a regression model, we were able to quantify the distinctness of these neurophysiological stages and identify the specific scalp topographies that drive group separation, which serves as a critical first step in defining stage-specific biomarkers. This approach allows the application of classifiers while maintaining balanced class sizes, with performance metrics providing objective validation of the models. To control for potential medication effects on brain activity within the FM group, a nonparametric permutation test was applied, given the small sample sizes and non-normal data distribution. Group-averaged PSD values were used for each channel and frequency band. To strictly control the family-wise error rate across all 60 channels and prevent type I errors, significance was assessed via 5000 permutations using the distribution of the maximum statistic (*t*-max) (Nichols & Holmes, 2002). This approach provides a single, stringent threshold for the entire scalp, ensuring that any reported differences are not due to multiple comparisons.

### 2.3 Classification models and performance evaluation

To distinguish between FM and HC, as well as between STFM and LTFM, linear discriminant analysis (LDA) and linear support vector machine (LSVM) models were trained. LSVM was defined using MATLAB’s *fitcsvm* function with a linear kernel, automatic kernel scaling, box constraint set to 1, and standardized predictors. LDA was built using MATLAB’s *fitcdiscr* with a linear discriminant type, no regularization (Gamma = 0), and fill coefficients enabled. Class names were explicitly specified in both models to ensure consistent label mapping. These models have previously been considered particularly well suited for EEG decoding in cognitive neuroscience due to their simplicity, high computational efficiency, and ability to handle high dimensional data (Cortes & Vapnik, 1995; Garrett et al., 2003; Sha’abani et al., 2020; Yu & Wang, 2022). Furthermore, they exhibit high robustness to noise, which is especially useful since EEG data, even after preprocessing, tend to contain leftover noise signals (Khanam et al., 2023). Given that both LDA and LSVM are well established frameworks for EEG classification, we performed an exploratory comparison of both models to ensure the robustness of our results across different algorithmic approaches, as it was not possible to determine a priori which would be most suitable for this specific itRS-EEG dataset.

Following PROBAST+AI guidelines (Moons et al., 2025), model performance was evaluated using accuracy (ACC), area under the receiver operating characteristic curve (AUC), recall, precision, F1-score, and calibration intercept and slope. Calibration was assessed following the weak calibration procedure described Van Calster et al. (2019). To mitigate potential overfitting, ten-fold cross-validation was employed. In this procedure, the dataset is split into ten subsets (folds) and in each iteration, one fold serves as the test set while the remaining nine are used for training. This process is repeated ten times so that each fold is used once for testing, and the performance metrics are then averaged across folds to obtain a robust estimate. Additionally, to account for data variability and assess the stability of the performance metrics, the cross-validation procedure was repeated 1000 times, and mean values were reported. As a baseline comparison, models were also trained on data with randomly permuted class labels. Given the relatively small sample size, no participants were reserved for external validation. However, in accordance with PROBAST+AI recommendations, this does not necessarily indicate a high risk of bias, provided that internal validation is rigorous and appropriately implemented. Overall judgment of prediction model was labeled as low across all four domains: development quality and applicability concern, and evaluation risk of bias and applicability concern. Full assessment is described in Supplementary Table 1. Finally, a two-tailed Wilcoxon signed-rank test was conducted to compare the classification performance of the LSVM and LDA models, in order to determine the most suitable approach.

### 2.4 Channel contribution analysis

Since our focus was on linear model variants, the weight assigned to each variable (i.e., EEG channel) after training can be extracted from the model parameters to represent its contribution to performance. However, these raw weights should not be treated as straightforward indicators of brain activity since a channel with a large weight does not necessarily indicate the presence of the signal of interest, and a channel with a small weight may still carry relevant information (Blankertz et al., 2011; Haufe et al., 2014). To improve neurophysiological interpretability, the weight vectors were transformed into activation patterns (corrected weights) by multiplying the weight vector by the data covariance matrix, following the method described by Haufe et al. (2014):

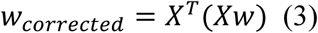

where *X* is the normalized data matrix and *w* is the weight vector obtained from the model. This transformation allows the resulting values to be more accurately interpreted as the spatial distribution of neural activity across scalp sensors (López-García et al., 2022). Furthermore, given that we considered the FM and STFM groups as true positive and the HC and LTFM groups as true negative in their respective analyses, positive values indicate a higher contribution of a particular channel to correctly classify an observation as FM or STFM, while negative values would reflect a greater contribution to classifying it as HC or LTFM.

## 3 Results

### 3.1 PSD spatial distribution differences

As described in the methods section, power spectral density (PSD) values from the itRS epochs were averaged within each group, and group-level difference (FM - HC and STFM - LTFM) maps were computed to explore spatial patterns prior to model input. The resulting spatial distributions are illustrated in Figure 2 and further detailed in Supplementary Tables 2 and 3. Relative to the HC group, the FM group showed a tendency toward concentrated activity in prefrontal regions across delta, theta, and alpha bands, as well as in prefrontal and right occipitotemporal areas for beta and gamma bands. In contrast, the HC group appeared to show concentrated activity in left frontotemporal and right frontal regions in beta and gamma bands. Furthermore, the STFM group’s activity appeared concentrated in prefrontal regions across all frequency bands, in the left frontotemporal region for delta, and in frontocentral and right frontal regions for beta and gamma bands. Meanwhile, the LTFM group’s activity was localized around occipitotemporal regions from delta to beta bands, and in left frontotemporal and right temporal regions for beta and gamma bands.

**Figure 2.**
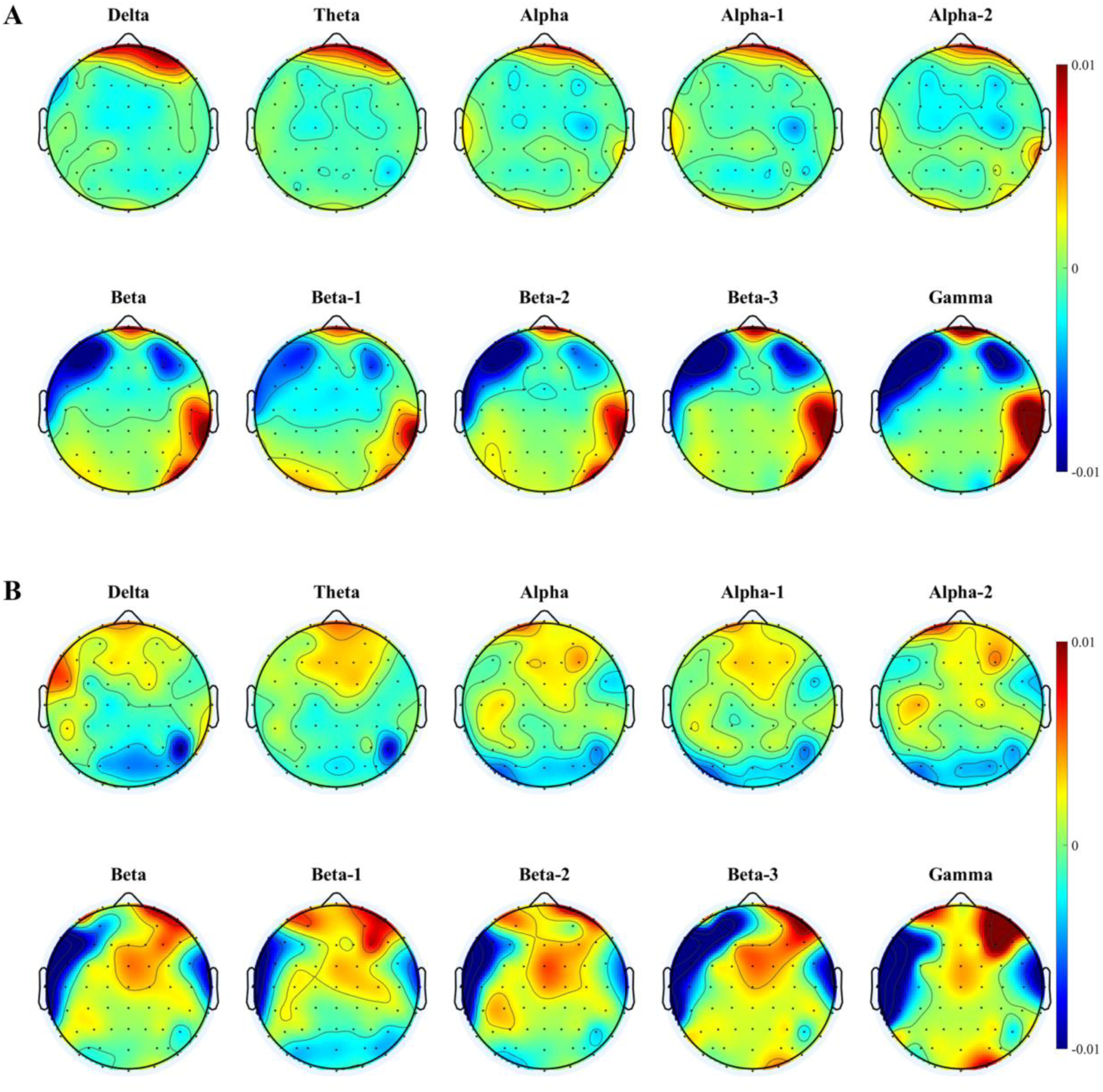
Group-level spatial distribution of normalized PSD across frequency bands. (**A**) Differences in normalized PSD between FM and HC (calculated as FM minus HC). (**B**) Differences in normalized PSD between STFM and LTFM (calculated as STFM minus LTFM). Warm colors indicate higher power in the first group (FM or STFM), while cool colors indicate higher power in the second group (HC or LTFM). The color scale represents the difference in relative power between groups. Note that because PSD was normalized by the total power of each observation, the values are unitless proportions.

### 3.2 Model performance

To identify the most suitable model for distinguishing patients from controls based on power spectral density (PSD) values, both linear discriminant analysis (LDA) and linear support vector machine (LSVM) models were trained across frequency bands. A two-tailed Wilcoxon signed-rank test revealed that LSVM significantly outperformed LDA (*p* < 0.001) in delta (*W* = 500500), theta (*W* = 500158), beta (*W* = 500499), beta-2 (*W* = 500500), beta-3 (*W* = 479925), and gamma (*W* = 500500) bands, whereas LDA showed superior performance in alpha (*W* = 0), alpha-1 (*W* = 6153), alpha-2 (*W* = 905), and beta-1 (*W* = 1602) bands. However, based on calibration scores, LDA demonstrated significantly greater overall stability, with slope values closer to 1 and intercept values closer to 0, except for the slope in the delta band and intercepts in beta-3 and gamma bands. Thus, LDA was selected for subsequent analyses. Performance metrics are illustrated in Figure 3 and further detailed in Supplementary Table 4.

**Figure 3.**
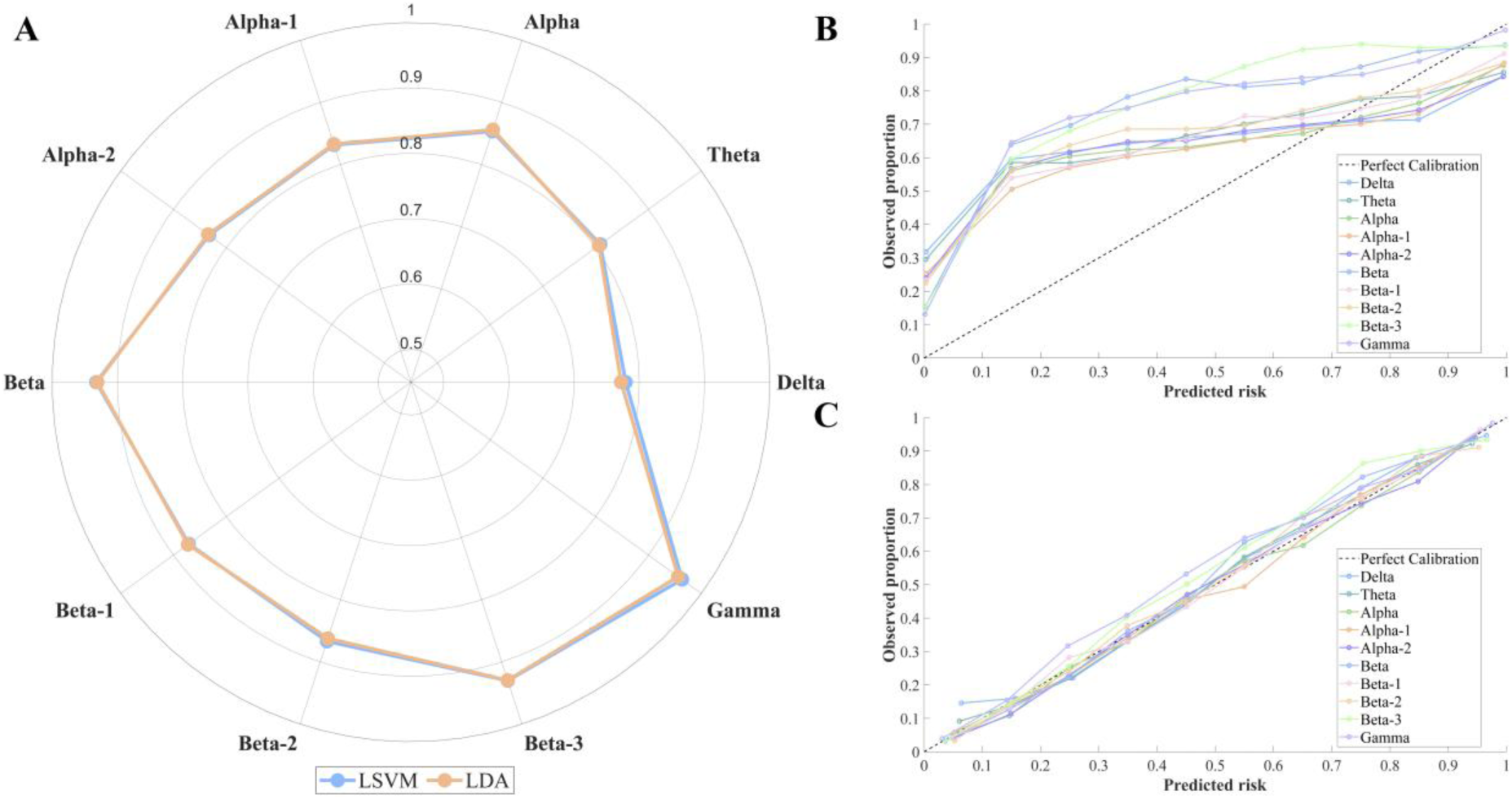
Comparative performance and reliability of classification models. (**A**) Classifier discrimination performance (AUC) across frequency bands using PSD values for LDA and LSVM models. Calibration curves per frequency band for (**B**) LSVM and (**C**) LDA showing observed outcome proportions (Y-axis) versus predicted probabilities (X-axis) ranging from 0 to 1. These curves evaluate how well the predicted risk matches actual outcomes. Points along the diagonal indicate perfect calibration, while deviations above or below the diagonal indicate under- or overestimation of risk. Calibration complements discrimination metrics (AUC) by assessing the reliability of predicted probabilities for each class across frequency bands.

### 3.3 Classification using discriminative features of EEG data

#### 3.3.1 EEG features and anxiety scores in FM classification

To distinguish HC from FM, power spectral density (PSD) from EEG data and psychometric values (PV) from the STAI questionnaire were used to train a linear discriminant analysis (LDA) model. Performance across feature combinations is shown in Figure 4 and detailed in Supplementary Table 5. To ensure the validity of our results, we performed a permutation test by randomly shuffling class labels. The resulting accuracy of approximately 0.5 (chance level) across all frequency bands indicates that the model is unable to extract discriminative patterns from the data in the absence of a true label-feature relationship, supporting that the observed performance reflects meaningful group-related structure rather than trivial dataset biases or label leakage. With PSD alone the model distinguished well between groups, particularly in beta-3 (*ACC =* 0.86, *AUC =* 0.93) and gamma (*ACC =* 0.89, *AUC =* 0.96) bands. When using anxiety scores alone, the model was also able to distinguish well between groups using STAI-S (*ACC =* 0.82, *AUC =* 0.91), STAI-T (*ACC =* 0.8, *AUC =* 0.92), and STAI-S + STAI-T (*ACC =* 0.86, *AUC =* 0.94). Combining either state (PSD + STAI-S) or trait (PSD + STAI-T) anxiety with PSD further improved performance across all bands, achieving AUC values above 0.99 for both beta-3 and gamma bands. The best distinction was obtained by combining all features, especially in beta-3 (*ACC =* 0.97, *AUC =* 1.00) and gamma (*ACC =* 0.98, *AUC =* 1.00). On the other hand, the calibration curves were relatively stable and close to the ideal diagonal line when using PSD alone, whereas the inclusion of PV led to greater deviations from perfect calibration, indicating over-confidence in the predicted probabilities for PSD + STAI-S, and under-confidence for PSD + STAI-T and PSD + STAI-S + STAI-T. These deviations indicate modest miscalibration in the combined-feature models.

**Figure 4.**
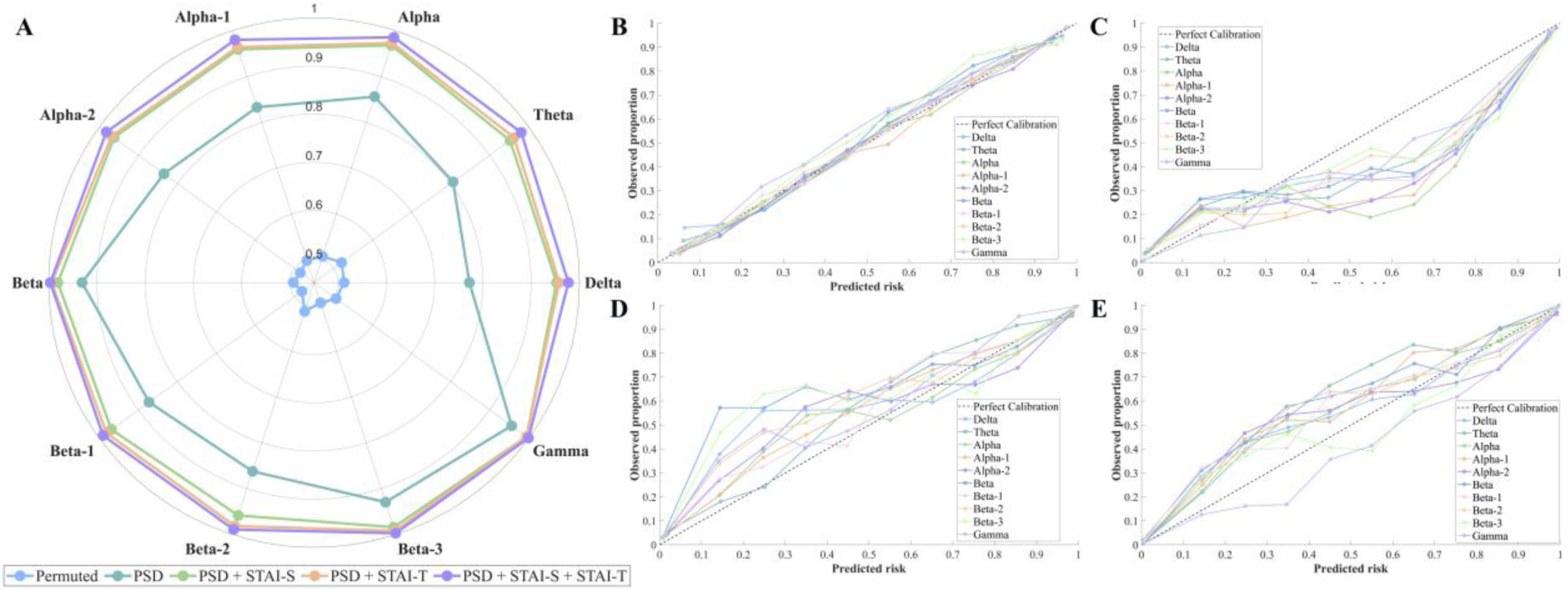
Integrative classification performance: EEG features and anxiety scores. (**A**) Classifier discrimination performance (AUC) across frequency bands for FM and HC. Calibration curves per frequency band for (**B**) PSD, (**C**) PSD + STAI-S, (**D**) PSD + STAI-T, and (**E**) PSD + STAI-S + STAI-T showing observed outcome proportions (Y-axis) versus predicted probabilities (X-axis) ranging from 0 to 1. These curves evaluate how well the predicted risk matches actual outcomes. Points along the diagonal indicate perfect calibration, while deviations above or below the diagonal indicate under- or overestimation of risk. Calibration complements discrimination metrics (AUC) by assessing the reliability of predicted probabilities for each class across frequency bands.

#### 3.3.2 Classifying STFM and LTFM using EEG features

To compare short-term fibromyalgia (STFM) and long-term fibromyalgia (LTFM) groups, power spectral density (PSD) from EEG data was used to train a linear discriminant analysis (LDA) model. Model performance is shown in Figure 5 and further detailed in Supplementary Table 6. The model successfully distinguished STFM from LTFM patients, particularly in beta-1 (*ACC* = 0.87, *AUC* = 0.94), beta-2 (*ACC* = 0.87, *AUC* = 0.94), beta-3 (*ACC* = 0.94, *AUC* = 0.98), and gamma (*ACC* = 0.96, *AUC* = 0.99) bands. Calibration curves were generally stable and close to the ideal diagonal, indicating good agreement between predicted probabilities and observed outcomes. However, beta-1, beta-3, and gamma bands showed slightly larger deviations from the diagonal, suggesting minor miscalibration in these frequency ranges.

**Figure 5.**
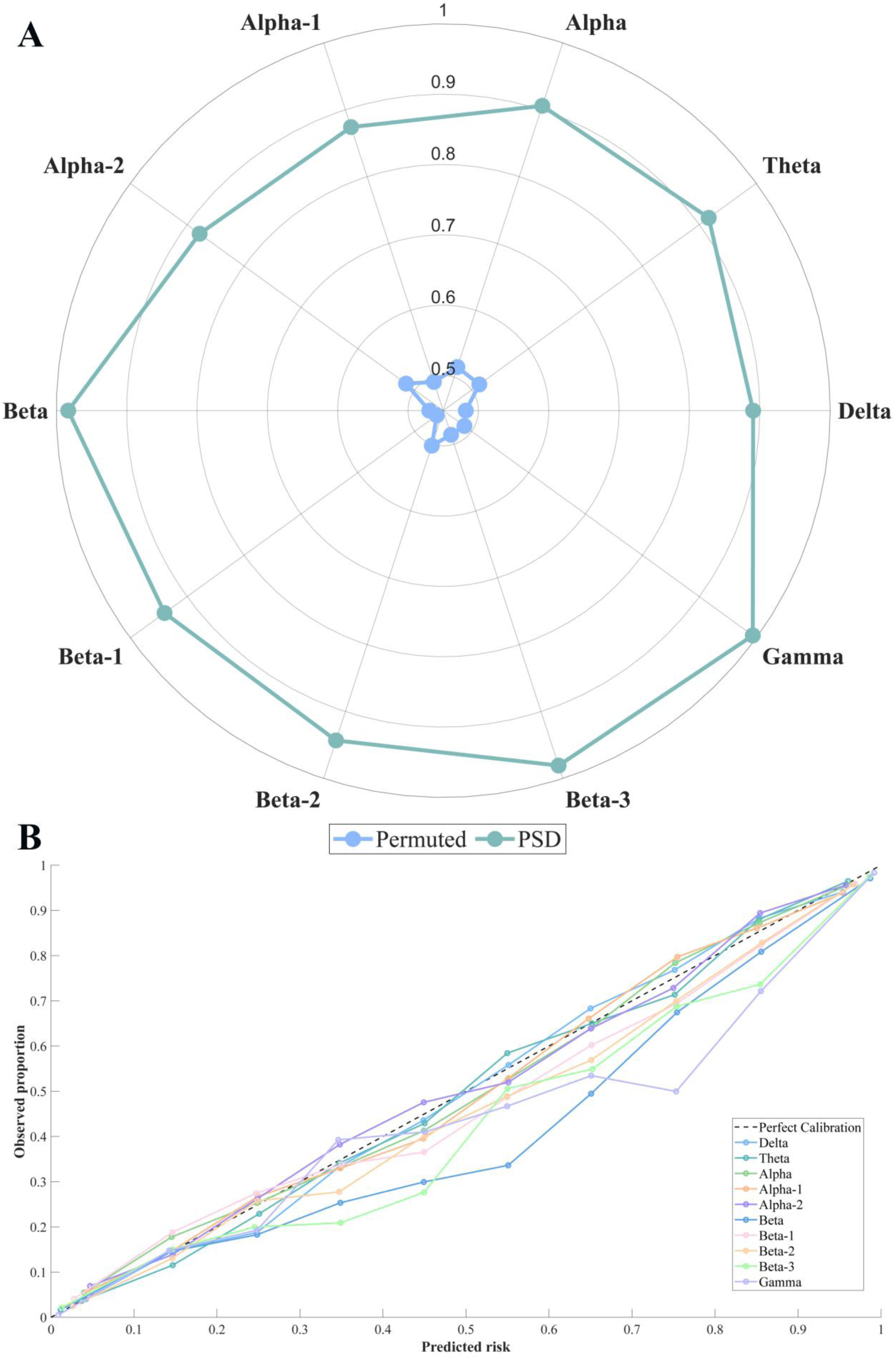
Performance metrics for symptom duration classification (STFM and LTFM). (**A**) Classifier discrimination performance (AUC) across frequency bands for STFM and LTFM. (**B**) Calibration curves per frequency band using PSD values showing observed outcome proportions (Y-axis) versus predicted probabilities (X-axis) ranging from 0 to 1. These curves evaluate how well the predicted risk matches actual outcomes. Points along the diagonal indicate perfect calibration, while deviations above or below the diagonal indicate under- or overestimation of risk. Calibration complements discrimination metrics (AUC) by assessing the reliability of predicted probabilities for each class across frequency bands.

#### 3.3.3 Channel contribution to group classification

To interpret the neural scalp sites contributing to classification, the correction method proposed by Haufe et al. (2014) was applied to the weight matrix obtained after classification. This transformation provides a more accurate spatial representation of relevant neural activity across frequency bands distinguishing FM from HC, as well as STFM from LTFM. The corrected weight maps are shown in Figure 6, and detailed in Supplementary Tables 7 and 8. In both comparisons, delta through alpha bands exhibited similar spatial patterns primarily in prefrontal regions. In the STFM and LTFM comparison, occipitotemporal regions also showed notable contributions. For beta through gamma bands, the FM and HC comparison revealed similar activity patterns in prefrontal, frontal, and temporoparietal regions. In contrast, the STFM and LTFM comparison highlighted significant contributions in temporal and frontal regions for beta and gamma bands, with additional involvement of parieto-occipital areas specifically for beta-3 and gamma.

**Figure 6.**
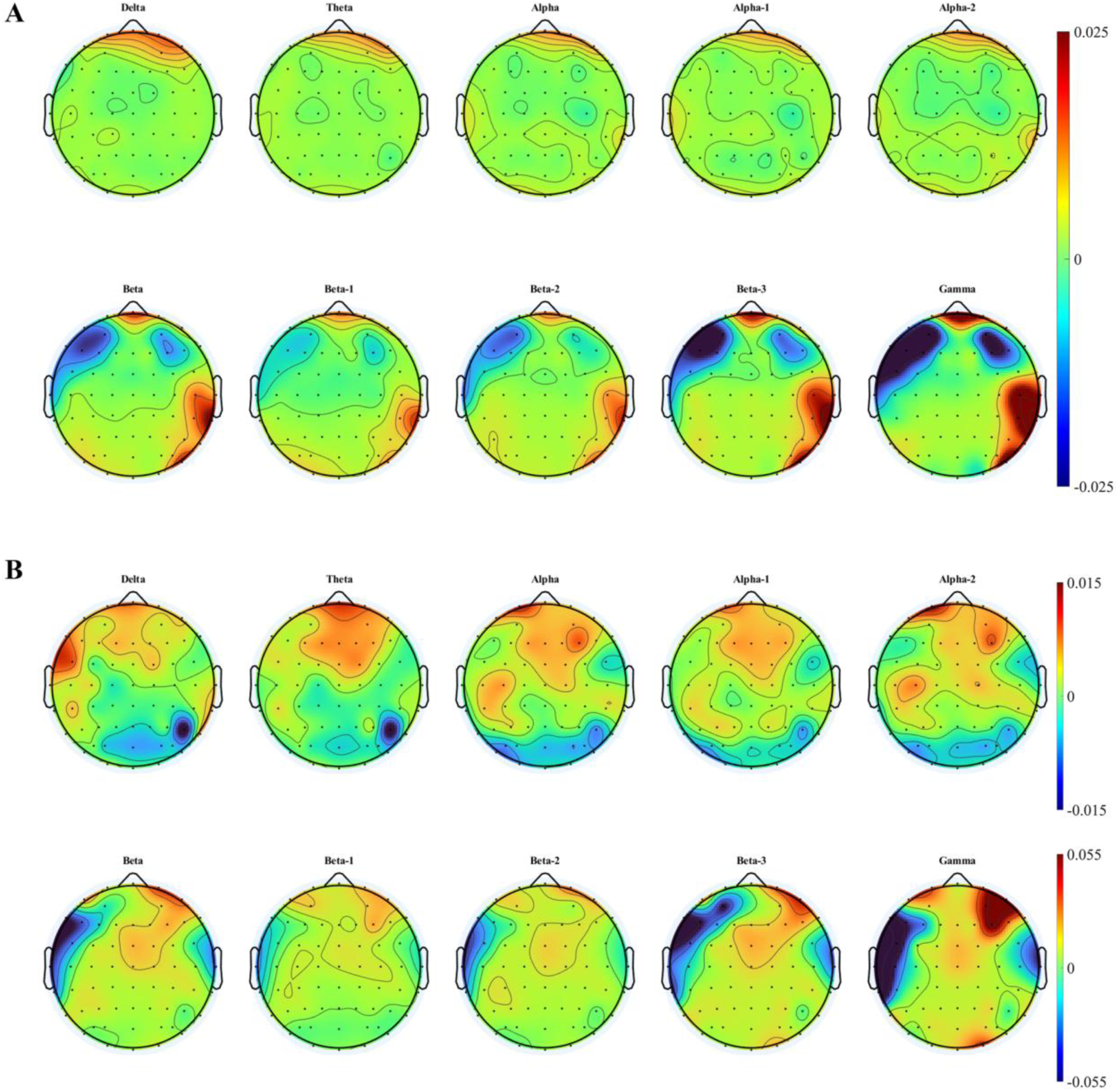
Topographical distribution of Haufe-transformed activation patterns across frequency bands. Scalp maps represent the activation patterns derived from Haufe-transformed model weights. (**A**) Scalp site contributions to the FM and HC classification; positive values (warm colors) indicate a higher contribution toward the FM group, while negative values (cool colors) indicate a higher contribution toward the HC group. (**B**) Scalp site contributions to the STFM and LTFM classification; positive values indicate a higher contribution toward the STFM group, while negative values indicate a higher contribution toward the LTFM group. The color scale represents model activation in arbitrary units; as these patterns are derived from normalized features, the values are dimensionless.

### 3.4 Medication effects on brain activity in the FM group

To assess potential medication effects on brain activity in FM patients, we conducted nonparametric permutation tests using the maximum statistic (*t*-max) correction method to compare mean PSD values between medicated and non-medicated patients. The results for each channel and frequency band are detailed in Supplementary Table 9. No significant differences were observed across the majority of comparisons (*p* > 0.05). While an isolated effect was observed at channel AF7 in the beta-2 band (*p* = 0.03) for the analgesics condition, the lack of spatial clustering or persistence across adjacent frequency bands suggests this was a spurious finding rather than a robust neurophysiological effect. Furthermore, sample sizes for anxiolytics and antiepileptics/psycholeptics (*N* = 6 and *N* = 4, respectively) were insufficient for meaningful statistical analysis.

## 4 Discussion

In the present study, we investigated abnormalities in EEG activity in FM patients during eyes open itRS. We applied LDA models in three novel analyses. First, we trained the models with PSD and anxiety scores to distinguish FM patients from HC. Second, we used diagnosis duration to distinguish between short-term FM (STFM) and long-term FM (LTFM) patients. Both analyses demonstrated high discriminative potential across frequency bands in this exploratory cohort. Third, we identified key scalp regions whose neural activity contributed most to the classification. This approach integrated EEG and psychometric data, linking brain activity to symptom duration and anxiety. By aligning our methodology with the study’s objectives, we were able to highlight the critical role these variables play in understanding the heterogeneity and complexity of FM.

Building on our integrated approach, we identified specific neural features enabling accurate classification of FM patients and HC. Notably, beta and gamma frequency bands emerged as the most discriminative. Within the beta band, particularly beta-3, classification performance was strongest, with scalp key contributions over bilateral frontal, right temporal, and right parieto-occipital regions. As the analysis was conducted in sensor space rather than source space, cortical interpretations should be done carefully. Nonetheless, brain areas situated under these scalp locations, such as the anterior insular, dorsolateral prefrontal, and secondary somatosensory cortices have been implicated in pain perception and its modulation, as well as the emotional and attentional processing frequently altered in FM (Alves et al., 2023; González-Villar et al., 2020; Villafaina, Collado-Mateo, & Fuentes-García, 2019). Moreover, beta-3 activity has been linked to interactions between sensory, affective, and attentional circuits involved in pain processing (Alves et al., 2023). Abnormal beta oscillations have been linked not only to pain processing but also to impairments in motor control, visual attention, and emotion regulation (Apkarian et al., 2004; Baliki et al., 2008; Engel & Fries, 2010; Hauck et al., 2015). While research on gamma activity in nociplastic pain is limited, increased gamma oscillations have been associated with heightened pain sensitivity and enhanced attentional focus on somatosensory stimuli (Alves et al., 2023; Hauck et al., 2007; Stefani et al., 2019). Furthermore, the high discriminative performance of these high-frequency bands in our model aligns with the thalamocortical dysrhythmia framework of chronic pain (Llinás et al., 1999). According to this model, persistent pain leads to a disruption of the resonant loops between the thalamus and the cortex, causing a shift in dominant rhythms and a concomitant edge effect increase in high frequency beta and gamma oscillations (Vanneste et al., 2018). In FM, this overexcitation likely reflects the neurophysiological noise of a centralized pain system (Cagnie et al., 2014; Fitzcharles et al., 2021) that is unable to return to a true state of rest.

Although beta oscillations relate to pain processing and broader cognitive functions, alpha and theta bands have been more directly linked to pain inhibition and cognitive control (Fallon et al., 2018; Villafaina, Collado-Mateo, & Fuentes-García, 2019). Specifically, RS studies report a negative correlation between pain levels and alpha power, indicating that chronic pain in FM patients may act as a persistent noxious stimulus that progressively diminishes alpha activity (Martín-Brufau et al., 2021; Villafaina, Collado-Mateo, & Fuentes-García, 2019). This aligns with our findings, where prefrontal regions exhibited strong positive weights contributing significantly to the classification of FM patients. Additionally, increased prefrontal theta PSD in FM might indicate diminished inhibitory control over pain processing. As previously suggested, this dysregulation could contribute to maladaptive feedback loops in which pain is amplified and poorly modulated, as brain areas near these scalp locations, such as the dorsolateral prefrontal and medial prefrontal cortices, are involved in these processes (Fallon et al., 2018; Henderson et al., 2013).

In addition to pain-related symptoms, emotional symptoms such as anxiety are common in FM patients and often more severe than in age-matched controls, contributing to reduced quality of life (Catalá et al., 2023; Cetingok et al., 2022; Henao-Pérez et al., 2022). Consistent with recent studies involving heat pain (Hsiao et al., 2021), our model’s performance improved greatly when combining either state or trait anxiety with PSD, achieving the highest accuracy when both anxiety scores were included. Notably, combining trait anxiety with PSD outperformed the combination of state anxiety with PSD. This agrees with previous research suggesting that although both state and trait anxiety can intensify pain perception in FM patients, trait anxiety shows a stronger association (Galvez-Sánchez et al., 2020). These findings highlight that integrating psychological and electrophysiological variables provides a more comprehensive characterization of FM than analyzing either dimension in isolation. Notably, EEG features alone achieved high discriminative performance (e.g., 0.96 AUC in the gamma band), suggesting that a robust neural signature of FM is present even in the absence of psychometric information. According to this, it appears that while psychometric scales capture subjective symptom severity, the EEG features provide an objective neurophysiological marker of the underlying state (Alves et al., 2023; Vanneste et al., 2017). Furthermore, the incremental improvement observed when combining both measures suggests that EEG may capture neurophysiological variance not fully reflected in self-report, potentially related to mechanisms of central sensitization and nociplastic pain (Clauw, 2024; Fitzcharles et al., 2021), that self-reported distress cannot fully reflect. Clinically, this neural fingerprint is vital for identifying patients who may present with atypical or low-anxiety profiles but still possess the same neurophysiological vulnerability. By incorporating objective EEG markers, we move toward a multimodal neuro-affective profile that ensures patients are not excluded from diagnosis or treatment based solely on subjective metrics.

Evidence from FM and other chronic pain conditions suggests that prolonged exposure to pain alters brain structures and disrupts functional connectivity (Apkarian et al., 2004; Baliki et al., 2008; Bekkelund et al., 1995; Jin et al., 2013; Kuchinad et al., 2007; McCrae et al., 2015; Rodriguez-Raecke et al., 2013). Although determining symptom onset in FM is challenging, as diagnosis often occurs years after initial symptoms, longer symptom duration appears to be associated with more pronounced neural alterations (Alves et al., 2023). This is consistent with our finding of high classification accuracy across all frequency bands when comparing STFM and LTFM groups. Additionally, altered theta power within frontal, frontocentral, and parietal regions has been associated with ongoing tonic pain and fatigue in FM (Fallon et al., 2018; Villafaina, Collado-Mateo, Fuentes-García, et al., 2019). In our study, corrected weight values revealed strong contributions from frontal and frontocentral regions to the classification of the STFM group, while parietal regions contributed more substantially to the LTFM group. Regarding other frequency bands, the observed patterns may indicate exacerbations due to prolonged exposure to FM symptoms in the same regions commonly affected in FM when compared to HC, particularly those involved in pain processing and cognitive control. However, given the relatively small subgroup sizes, these symptom-duration findings should be interpreted as exploratory and require confirmation in larger, fully independent cohorts.

A key consideration in the current study is the nature of the itRS data. Unlike conventional RS recordings where participants are instructed to remain still without external stimulation for several minutes, our data were extracted from the 3000 ms ITI of an active task. While task-related responses and visual triggers were carefully removed, these segments may still include residual neural processes. Notably, recent work suggests that short inter-trial or post-task intervals can meaningfully capture ongoing neural activity and task-related reactivation, providing insight into background cortical dynamics (D’Croz-Baron et al., 2021; Murphy et al., 2018; Walz et al., 2014). In FM, these intervals may additionally reflect heightened vigilance or anticipatory processes, which are characteristic of the condition (Cagnie et al., 2014). Therefore, while our results should not be interpreted as classical RS activity, they likely represent a semi-active state that captures a stable cognitive set of sustained vigilance and expectancy, which may be particularly relevant in FM populations (Fernandes-Magalhaes et al., 2022, 2024).

The current study has several limitations that necessitate a cautious interpretation of the results. First, while our repeated cross-validation provides a stable estimate of model performance, the dataset includes multiple epochs per participant, which could introduce subtle trial-level dependencies between training and test folds. Along with the relatively small sample size and lack of an independent validation dataset, this means that reported classification accuracies may be somewhat optimistic and should be considered exploratory. In line with this, calibration curves indicated minor deviations from ideal probability calibration, suggesting some degree of overconfidence and reinforcing the need for independent validation. Second, while the channels showing significant medication effects did not appear to drive the classification, the impact of long-term pharmacological treatment on cortical oscillations cannot be entirely dismissed. Finally, although task-related activity was carefully removed to create an itRS design, task anticipatory neural activity and carryover effects from preceding trials might still influence background activity, supporting the need for future studies to replicate these patterns using conventional RS recordings.

In sum, our findings reveal significant differences in itRS-EEG activity between FM patients and HC, as well as between STFM and LTFM. Incorporating anxiety scores significantly improved model performance, supporting a neuro-affective profile that captures the interplay between emotional and neural factors in FM. By viewing these markers through this integrative lens, we acknowledge that in FM, nociplastic pain and affective dysregulation are often clinically and physiologically inseparable. These results highlight the discriminative power of beta and gamma oscillations, particularly in regions associated with pain processing and cognitive control. Beyond their diagnostic potential, these high frequency signatures offer a translational tool for monitoring treatment response. For instance, normalizing these oscillations could serve as an objective measure of efficacy for interventions like pharmacotherapy or neuromodulation, such as repetitive transcranial magnetic stimulation or transcranial direct current stimulation. While these exploratory findings require independent validation, they represent a significant step toward objective, personalized management of FM.

## 5 Data availability statement

The data supporting the findings of this study are available in the OSF repository at https://osf.io/gyk89/

## 6 Acknowledgements

The authors would like to thank all participants for their involvement in the study. A preprint of this work has previously been published on bioRxiv (Soldic et al., 2025).

## 7 Conflict of interests

The authors declare that they have no known competing financial interests or personal relationships that could have appeared to influence the work reported in this paper.

## 8 CRediT authorship contribution statement

**Dino Soldic:** Conceptualization, Data curation, Formal analysis, Methodology, Software, Visualization, Writing – original draft, Writing – review & editing. **María Carmen Martín-Buro:** Conceptualization, Formal analysis, Software, Supervision, Validation, Writing – review & editing. **David López-García:** Methodology, Software, Formal analysis, Supervision, Validation, Writing – review & editing. **Ana Belén del Pino:** Data curation, Investigation, Methodology, Visualization, Writing – review & editing. **Roberto Fernandes-Magalhaes:** Conceptualization, Data curation, Investigation, Methodology, Supervision, Writing – review & editing. **David Ferrera:** Investigation, Methodology, Supervision, Writing – review & editing. **Irene Peláez:** Investigation, Methodology, Supervision, Writing – review & editing. **Luis Carretié:** Investigation, Resources, Supervision, Writing – review & editing. **Francisco Mercado:** Conceptualization, Funding Acquisition, Investigation, Methodology, Project administration, Resources, Supervision, Validation, Writing – review & editing.

## 9 Funding statement

This work was supported by grant PID2020-115463RB-I00 from the Ministerio de Ciencia e Innovación of Spain.

## 10 Declaration of generative AI use

No artificial intelligence assisted technologies were used in the conception of the study, generation of original ideas, or data analysis of this manuscript. Only standard editorial tools (e.g., spelling and grammar checks) were used.

## Notes

https://osf.io/gyk89/

### Competing Interest Statement

The authors have declared no competing interest.

### Summary of Updates

This version of the manuscript has been revised to update the information that will be reflected upon the article's publication.

## References

Abdellatef, E., Emara, H. M., Shoaib, M. R., Ibrahim, F. E., Elwekeil, M., El-Shafai, W., Taha, T. E., El-Fishawy, A. S., El-Rabaie, E.-S. M., Eldokany, I. M., & Abd El-Samie, F. E. (2023). Automated diagnosis of EEG abnormalities with different classification techniques. Medical & Biological Engineering & Computing, 61(12), 3363–3385. 10.1007/s11517-023-02843-w

Alves, R. L., Zortea, M., Serrano, P. V., Brugnera Tomedi, R., Pereira De Almeida, R., Torres, I. L. S., Fregni, F., & Caumo, W. (2023). High-beta oscillations at EEG resting state and hyperconnectivity of pain circuitry in fibromyalgia: An exploratory cross-sectional study. Frontiers in Neuroscience, 17, 1233979. 10.3389/fnins.2023.1233979

Amris, K., Luta, G., Christensen, R., Danneskiold-SamsÃe, B., Bliddal, H., & WÃ¦hrens, E. (2016). Predictors of improvement in observed functional ability in patients with fibromyalgia as an outcome of rehabilitation. Journal of Rehabilitation Medicine, 48(1), 65–71. 10.2340/16501977-2036

Apkarian, A. V., Sosa, Y., Sonty, S., Levy, R. M., Harden, R. N., Parrish, T. B., & Gitelman, D. R. (2004). Chronic Back Pain Is Associated with Decreased Prefrontal and Thalamic Gray Matter Density. The Journal of Neuroscience, 24(46), 10410–10415. 10.1523/JNEUROSCI.2541-04.2004

Baliki, M. N., Geha, P. Y., Apkarian, A. V., & Chialvo, D. R. (2008). Beyond Feeling: Chronic Pain Hurts the Brain, Disrupting the Default-Mode Network Dynamics. The Journal of Neuroscience, 28(6), 1398–1403. 10.1523/JNEUROSCI.4123-07.2008

Bekkelund, S. I., Pierre-Jerome, C., Husby, G., & Mellgren, S. I. (1995). Quantitative cerebral MR in rheumatoid arthritis. AJNR. American Journal of Neuroradiology, 16(4), 767–772.

Blankertz, B., Lemm, S., Treder, M., Haufe, S., & Müller, K.-R. (2011). Single-trial analysis and classification of ERP components—A tutorial. NeuroImage, 56(2), 814–825. 10.1016/j.neuroimage.2010.06.048

Bushnell, M. C., Čeko, M., & Low, L. A. (2013). Cognitive and emotional control of pain and its disruption in chronic pain. Nature Reviews Neuroscience, 14(7), 502–511. 10.1038/nrn3516

Cagnie, B., Coppieters, I., Denecker, S., Six, J., Danneels, L., & Meeus, M. (2014). Central sensitization in fibromyalgia? A systematic review on structural and functional brain MRI. Seminars in Arthritis and Rheumatism, 44(1), 68–75. 10.1016/j.semarthrit.2014.01.001

Carretié, L. (2014). Exogenous (automatic) attention to emotional stimuli: A review. Cognitive, Affective, & Behavioral Neuroscience, 14(4), 1228–1258. 10.3758/s13415-014-0270-2

Catalá, P., Gutiérrez, L., Écija, C., & Peñacoba, C. (2023). Pathological Cycle between Pain, Insomnia, and Anxiety in Women with Fibromyalgia and its Association with Disease Impact. Biomedicines, 11(1), 148. 10.3390/biomedicines11010148

Cetingok, S., Seker, O., & Cetingok, H. (2022). The relationship between fibromyalgia and depression, anxiety, anxiety sensitivity, fear avoidance beliefs, and quality of life in female patients. Medicine, 101(39), e30868. 10.1097/MD.0000000000030868

Choe, M. K., Lim, M., Kim, J. S., Lee, D. S., & Chung, C. K. (2018). Disrupted Resting State Network of Fibromyalgia in Theta frequency. Scientific Reports, 8(1), 2064. 10.1038/s41598-017-18999-z

Clauw, D. J. (2014). Fibromyalgia: A Clinical Review. JAMA, 311(15), 1547. 10.1001/jama.2014.3266

Clauw, D. J. (2024). From fibrositis to fibromyalgia to nociplastic pain: How rheumatology helped get us here and where do we go from here? Annals of the Rheumatic Diseases, 83(11), 1421–1427. 10.1136/ard-2023-225327

Cortes, C., & Vapnik, V. (1995). Support-vector networks. Machine Learning, 20(3), 273–297. 10.1007/BF00994018

D’Croz-Baron, D. F., Bréchet, L., Baker, M., & Karp, T. (2021). Auditory and Visual Tasks Influence the Temporal Dynamics of EEG Microstates During Post-encoding Rest. Brain Topography, 34(1), 19–28. 10.1007/s10548-020-00802-4

Delorme, A., & Makeig, S. (2004). EEGLAB: An open source toolbox for analysis of single-trial EEG dynamics including independent component analysis. Journal of Neuroscience Methods, 134(1), 9–21. 10.1016/j.jneumeth.2003.10.009

Engel, A. K., & Fries, P. (2010). Beta-band oscillations—Signalling the status quo? Current Opinion in Neurobiology, 20(2), 156–165. 10.1016/j.conb.2010.02.015

Fallon, N., Chiu, Y., Nurmikko, T., & Stancak, A. (2018). Altered theta oscillations in resting EEG of fibromyalgia syndrome patients. European Journal of Pain, 22(1), 49–57. 10.1002/ejp.1076

Fernandes-Magalhaes, R., Carpio, A., Ferrera, D., Peláez, I., De Lahoz, M. E., Van Ryckeghem, D., Van Damme, S., & Mercado, F. (2024). Neural mechanisms underlying attentional bias modification in fibromyalgia patients: A double-blind ERP study. European Archives of Psychiatry and Clinical Neuroscience, 274(5), 1197–1213. 10.1007/s00406-023-01709-4

Fernandes-Magalhaes, R., Ferrera, D., Peláez, I., Martín-Buro, M. C., Carpio, A., De Lahoz, M. E., Barjola, P., & Mercado, F. (2022). Neural correlates of the attentional bias towards pain-related faces in fibromyalgia patients: An ERP study using a dot-probe task. Neuropsychologia, 166, 108141. 10.1016/j.neuropsychologia.2021.108141

Ferrera, D., Gómez-Esquer, F., Peláez, I., Barjola, P., Fernandes-Magalhaes, R., Carpio, A., De Lahoz, M. E., Díaz-Gil, G., & Mercado, F. (2020). Effects of COMT Genotypes on Working Memory Performance in Fibromyalgia Patients. Journal of Clinical Medicine, 9(8), 2479. 10.3390/jcm9082479

Ferrera, D., Mercado, F., Peláez, I., Martínez-Iñigo, D., Fernandes-Magalhaes, R., Barjola, P., Écija, C., Díaz-Gil, G., & Gómez-Esquer, F. (2021). Fear of pain moderates the relationship between self-reported fatigue and methionine allele of catechol-O-methyltransferase gene in patients with fibromyalgia. PLOS ONE, 16(4), e0250547. 10.1371/journal.pone.0250547

Fitzcharles, M.-A., Cohen, S. P., Clauw, D. J., Littlejohn, G., Usui, C., & Häuser, W. (2021). Nociplastic pain: Towards an understanding of prevalent pain conditions. The Lancet, 397(10289), 2098–2110. 10.1016/s0140-6736(21)00392-5

Galvez-Sánchez, C. M., Montoro, C. I., Duschek, S., & Reyes Del Paso, G. A. (2020). Depression and trait-anxiety mediate the influence of clinical pain on health-related quality of life in fibromyalgia. Journal of Affective Disorders, 265, 486–495. 10.1016/j.jad.2020.01.129

Garcia-Larrea, L., & Bastuji, H. (2018). Pain and consciousness. Progress in Neuro-Psychopharmacology and Biological Psychiatry, 87, 193–199. 10.1016/j.pnpbp.2017.10.007

Garrett, D., Peterson, D. A., Anderson, C. W., & Thaut, M. H. (2003). Comparison of linear, nonlinear, and feature selection methods for EEG signal classification. IEEE Transactions on Neural Systems and Rehabilitation Engineering, 11(2), 141–144. 10.1109/TNSRE.2003.814441

Giorgi, V., Sirotti, S., Romano, M. E., Marotto, D., Ablin, J. N., Salaffi, F., & Sarzi-Puttini, P. (2022). Review Fibromyalgia: One year in review 2022. Clinical and Experimental Rheumatology.

González-Villar, A. J., Triñanes, Y., Gómez-Perretta, C., & Carrillo-de-la-Peña, M. T. (2020). Patients with fibromyalgia show increased beta connectivity across distant networks and microstates alterations in resting-state electroencephalogram. NeuroImage, 223, 117266. 10.1016/j.neuroimage.2020.117266

Hauck, M., Domnick, C., Lorenz, J., Gerloff, C., & Engel, A. K. (2015). Top-down and bottom-up modulation of pain-induced oscillations. Frontiers in Human Neuroscience, 9. 10.3389/fnhum.2015.00375

Hauck, M., Lorenz, J., & Engel, A. K. (2007). Attention to Painful Stimulation Enhances γ-Band Activity and Synchronization in Human Sensorimotor Cortex. The Journal of Neuroscience, 27(35), 9270–9277. 10.1523/JNEUROSCI.2283-07.2007

Haufe, S., Meinecke, F., Görgen, K., Dähne, S., Haynes, J.-D., Blankertz, B., & Bießmann, F. (2014). On the interpretation of weight vectors of linear models in multivariate neuroimaging. NeuroImage, 87, 96–110. 10.1016/j.neuroimage.2013.10.067

Henao-Pérez, M., López-Medina, D. C., Arboleda, A., Bedoya Monsalve, S., & Zea, J. A. (2022). Patients With Fibromyalgia, Depression, and/or Anxiety and Sex Differences. American Journal of Men’s Health, 16(4), 15579883221110351. 10.1177/15579883221110351

Henderson, L. A., Peck, C. C., Petersen, E. T., Rae, C. D., Youssef, A. M., Reeves, J. M., Wilcox, S. L., Akhter, R., Murray, G. M., & Gustin, S. M. (2013). Chronic Pain: Lost Inhibition? The Journal of Neuroscience, 33(17), 7574–7582. 10.1523/JNEUROSCI.0174-13.2013

Hsiao, F.-J., Chen, W.-T., Pan, L.-L. H., Liu, H.-Y., Wang, Y.-F., Chen, S.-P., Lai, K.-L., & Wang, S.-J. (2021). Machine learning–based prediction of heat pain sensitivity by using resting-state EEG. Frontiers in Bioscience-Landmark, 26(12), 1537–1547. 10.52586/5047

Hsiao, F.-J., Wang, S.-J., Lin, Y.-Y., Fuh, J.-L., Ko, Y.-C., Wang, P.-N., & Chen, W.-T. (2017). Altered insula–default mode network connectivity in fibromyalgia: A resting-state magnetoencephalographic study. The Journal of Headache and Pain, 18(1), 89. 10.1186/s10194-017-0799-x

Iemi, L., Busch, N. A., Laudini, A., Haegens, S., Samaha, J., Villringer, A., & Nikulin, V. V. (2019). Multiple mechanisms link prestimulus neural oscillations to sensory responses. eLife, 8, e43620. 10.7554/eLife.43620

Jin, C., Yuan, K., Zhao, L., Zhao, L., Yu, D., Von Deneen, K. M., Zhang, M., Qin, W., Sun, W., & Tian, J. (2013). Structural and functional abnormalities in migraine patients without aura. NMR in Biomedicine, 26(1), 58–64. 10.1002/nbm.2819

Khanam, F., Hossain, A. B. M. A., & Ahmad, M. (2023). Electroencephalogram-based cognitive load level classification using wavelet decomposition and support vector machine. Brain-Computer Interfaces, 10(1), 1–15. 10.1080/2326263X.2022.2109855

Kim, J., Loggia, M. L., Cahalan, C. M., Harris, R. E., Beissner, F., Garcia, R. G., Kim, H., Barbieri, R., Wasan, A. D., Edwards, R. R., & Napadow, V. (2015). The Somatosensory Link in Fibromyalgia: Functional Connectivity of the Primary Somatosensory Cortex Is Altered by Sustained Pain and Is Associated With Clinical/Autonomic Dysfunction. Arthritis & Rheumatology, 67(5), 1395–1405. 10.1002/art.39043

Kuchinad, A., Schweinhardt, P., Seminowicz, D. A., Wood, P. B., Chizh, B. A., & Bushnell, M. C. (2007). Accelerated Brain Gray Matter Loss in Fibromyalgia Patients: Premature Aging of the Brain? The Journal of Neuroscience, 27(15), 4004–4007. 10.1523/JNEUROSCI.0098-07.2007

Kumbhare, D., Ahmed, S., & Watter, S. (2018). A narrative review on the difficulties associated with fibromyalgia diagnosis. Therapeutic Advances in Musculoskeletal Disease, 10(1), 13–26. 10.1177/1759720X17740076

Llinás, R. R., Ribary, U., Jeanmonod, D., Kronberg, E., & Mitra, P. P. (1999). Thalamocortical dysrhythmia: A neurological and neuropsychiatric syndrome characterized by magnetoencephalography. Proceedings of the National Academy of Sciences, 96(26), 15222–15227. 10.1073/pnas.96.26.15222

López-García, D., Peñalver, J. M. G., Górriz, J. M., & Ruz, M. (2022). MVPAlab: A machine learning decoding toolbox for multidimensional electroencephalography data. Computer Methods and Programs in Biomedicine, 214, 106549. 10.1016/j.cmpb.2021.106549

López-Solà, M., Woo, C.-W., Pujol, J., Deus, J., Harrison, B. J., Monfort, J., & Wager, T. D. (2017). Towards a neurophysiological signature for fibromyalgia. Pain, 158(1), 34–47. 10.1097/j.pain.0000000000000707

MacCallum, R. C., Zhang, S., Preacher, K. J., & Rucker, D. D. (2002). On the practice of dichotomization of quantitative variables. Psychological Methods, 7(1), 19–40. 10.1037/1082-989X.7.1.19

Maris, E., & Oostenveld, R. (2007). Nonparametric statistical testing of EEG- and MEG-data. Journal of Neuroscience Methods, 164(1), 177–190. 10.1016/j.jneumeth.2007.03.024

Martín-Brufau, R., Gómez, M. N., Sanchez-Sanchez-Rojas, L., & Nombela, C. (2021). Fibromyalgia Detection Based on EEG Connectivity Patterns. Journal of Clinical Medicine, 10(15), 3277. 10.3390/jcm10153277

McCrae, C., O’Shea, A., Boissoneault, J., Vatthauer, K., Robinson, M., Staud, R., Perlstein, W., & Craggs, J. (2015). Fibromyalgia patients have reduced hippocampal volume compared with healthy controls. Journal of Pain Research, 47. 10.2147/JPR.S71959

Moons, K. G. M., Damen, J. A. A., Kaul, T., Hooft, L., Andaur Navarro, C., Dhiman, P., Beam, A. L., Van Calster, B., Celi, L. A., Denaxas, S., Denniston, A. K., Ghassemi, M., Heinze, G., Kengne, A. P., Maier-Hein, L., Liu, X., Logullo, P., McCradden, M. D., Liu, N., … Van Smeden, M. (2025). PROBAST+AI: An updated quality, risk of bias, and applicability assessment tool for prediction models using regression or artificial intelligence methods. BMJ, e082505. 10.1136/bmj-2024-082505

Murphy, M., Stickgold, R., Parr, M. E., Callahan, C., & Wamsley, E. J. (2018). Recurrence of task-related electroencephalographic activity during post-training quiet rest and sleep. Scientific Reports, 8(1), 5398. 10.1038/s41598-018-23590-1

Nichols, T. E., & Holmes, A. P. (2002). Nonparametric permutation tests for functional neuroimaging: A primer with examples. Human Brain Mapping, 15(1), 1–25. 10.1002/hbm.1058

Oostenveld, R., Fries, P., Maris, E., & Schoffelen, J.-M. (2011). FieldTrip: Open Source Software for Advanced Analysis of MEG, EEG, and Invasive Electrophysiological Data. Computational Intelligence and Neuroscience, 2011, 1–9. 10.1155/2011/156869

Ploghaus, A., Narain, C., Beckmann, C. F., Clare, S., Bantick, S., Wise, R., Matthews, P. M., Rawlins, J. N. P., & Tracey, I. (2001). Exacerbation of Pain by Anxiety Is Associated with Activity in a Hippocampal Network. The Journal of Neuroscience, 21(24), 9896–9903. 10.1523/JNEUROSCI.21-24-09896.2001

Pujol, J., Blanco-Hinojo, L., Doreste, A., Ojeda, F., Martínez-Vilavella, G., Pérez-Sola, V., Deus, J., & Monfort, J. (2022). Distinctive alterations in the functional anatomy of the cerebral cortex in pain-sensitized osteoarthritis and fibromyalgia patients. Arthritis Research & Therapy, 24(1), 252. 10.1186/s13075-022-02942-3

Pujol, J., Macià, D., Garcia-Fontanals, A., Blanco-Hinojo, L., López-Solà, M., Garcia-Blanco, S., Poca-Dias, V., Harrison, B. J., Contreras-Rodríguez, O., Monfort, J., Garcia-Fructuoso, F., & Deus, J. (2014). The contribution of sensory system functional connectivity reduction to clinical pain in fibromyalgia. Pain, 155(8), 1492–1503. 10.1016/j.pain.2014.04.028

Raichle, M. E., & Snyder, A. Z. (2007). A default mode of brain function: A brief history of an evolving idea. NeuroImage, 37(4), 1083–1090. 10.1016/j.neuroimage.2007.02.041

Rodriguez-Raecke, R., Niemeier, A., Ihle, K., Ruether, W., & May, A. (2013). Structural Brain Changes in Chronic Pain Reflect Probably Neither Damage Nor Atrophy. PLoS ONE, 8(2), e54475. 10.1371/journal.pone.0054475

Ruiz De Miras, J., Ibáñez-Molina, A. J., Soriano, M. F., & Iglesias-Parro, S. (2023). Schizophrenia classification using machine learning on resting state EEG signal. Biomedical Signal Processing and Control, 79, 104233. 10.1016/j.bspc.2022.104233

Saviola, F., Pappaianni, E., Monti, A., Grecucci, A., Jovicich, J., & De Pisapia, N. (2020). Trait and state anxiety are mapped differently in the human brain. Scientific Reports, 10(1), 11112. 10.1038/s41598-020-68008-z

Sha’abani, M. N. A. H., Fuad, N., Jamal, N., & Ismail, M. F. (2020). kNN and SVM Classification for EEG: A Review. In A. N. Kasruddin Nasir, M. A. Ahmad, M. S. Najib, Y. Abdul Wahab, N. A. Othman, N. M. Abd Ghani, A. Irawan, S. Khatun, R. M. T. Raja Ismail, M. M. Saari, M. R. Daud, & A. A. Mohd Faudzi (Eds.), InECCE2019 (Vol. 632, pp. 555–565). Springer Singapore. 10.1007/978-981-15-2317-5_47

Soldic, D., Martín-Buro, M. C., López-García, D., Del Pino, A. B., Fernandes-Magalhaes, R., Ferrera, D., Peláez, I., Carretié, L., & Mercado, F. (2025). Multivariate pattern analysis reveals resting-state EEG biomarkers in fibromyalgia. bioRxiv [Preprint]. 10.1101/2025.05.30.657056

Spielberger, C. D., Gorsuch, R. L., & Lushene, E. E. (1982). Manual del Cuestionario de Ansiedad Estado/Rasgo (STAI). TEA Ediciones, Madrid.

Stefani, L. C., Leite, F. M., Da Graça L. Tarragó, M., Zanette, S. A., De Souza, A., Castro, S. M., & Caumo, W. (2019). BDNF and serum S100B levels according the spectrum of structural pathology in chronic pain patients. Neuroscience Letters, 706, 105–109. 10.1016/j.neulet.2019.05.021

Thanh Nhu, N., Chen, D. Y.-T., & Kang, J.-H. (2022). Identification of Resting-State Network Functional Connectivity and Brain Structural Signatures in Fibromyalgia Using a Machine Learning Approach. Biomedicines, 10(12), 3002. 10.3390/biomedicines10123002

The MathWorks, Inc. (2023). *MATLAB* (Version 2023a) [Computer software]. https://www.mathworks.com/

Van Calster, B., McLernon, D. J., Van Smeden, M., Wynants, L., & Steyerberg, E. W. (2019). Calibration: The Achilles heel of predictive analytics. BMC Medicine, 17(1), 230. 10.1186/s12916-019-1466-7

Vanneste, S., Ost, J., Van Havenbergh, T., & De Ridder, D. (2017). Resting state electrical brain activity and connectivity in fibromyalgia. PLOS ONE, 12(6), e0178516. 10.1371/journal.pone.0178516

Vanneste, S., Song, J.-J., & De Ridder, D. (2018). Thalamocortical dysrhythmia detected by machine learning. Nature Communications, 9(1), 1103. 10.1038/s41467-018-02820-0

Villafaina, S., Collado-Mateo, D., & Fuentes-García, J. P. (2019). Impact of Fibromyalgia on Alpha-2 EEG Power Spectrum in the Resting Condition: A Descriptive Correlational Study. BioMed Research International, 2019, 1–6. 10.1155/2019/7851047

Villafaina, S., Collado-Mateo, D., Fuentes-García, J. P., Domínguez-Muñoz, F. J., & Gusi, N. (2019). Duration of the Symptoms and Brain Aging in Women with Fibromyalgia: A Cross-Sectional Study. Applied Sciences, 9(10), 2106. 10.3390/app9102106

Walz, J. M., Goldman, R. I., Carapezza, M., Muraskin, J., Brown, T. R., & Sajda, P. (2014). Simultaneous EEG–fMRI reveals a temporal cascade of task-related and default-mode activations during a simple target detection task. NeuroImage, 102, 229–239. 10.1016/j.neuroimage.2013.08.014

Wolfe, F., Clauw, D. J., Fitzcharles, M., Goldenberg, D. L., Katz, R. S., Mease, P., Russell, A. S., Russell, I. J., Winfield, J. B., & Yunus, M. B. (2010). The American College of Rheumatology Preliminary Diagnostic Criteria for Fibromyalgia and Measurement of Symptom Severity. Arthritis Care & Research, 62(5), 600–610. 10.1002/acr.20140

Wolfe, F., Clauw, D. J., Fitzcharles, M.-A., Goldenberg, D. L., Häuser, W., Katz, R. L., Mease, P. J., Russell, A. S., Russell, I. J., & Walitt, B. (2016). 2016 Revisions to the 2010/2011 fibromyalgia diagnostic criteria. Seminars in Arthritis and Rheumatism, 46(3), 319–329. 10.1016/j.semarthrit.2016.08.012

Wolfe, F., Smythe, H. A., Yunus, M. B., Bennett, R. M., Bombardier, C., Goldenberg, D. L., Tugwell, P., Campbell, S. M., Abeles, M., Clark, P., Fam, A. G., Farber, S. J., Fiechtner, J. J., Michael Franklin, C., Gatter, R. A., Hamaty, D., Lessard, J., Lichtbroun, A. S., Masi, A. T., … Sheon, R. P. (1990). The american college of rheumatology 1990 criteria for the classification of fibromyalgia. Arthritis & Rheumatism, 33(2), 160–172. 10.1002/art.1780330203

Yu, C., & Wang, M. (2022). Survey of emotion recognition methods using EEG information. Cognitive Robotics, 2, 132–146. 10.1016/j.cogr.2022.06.001

Zafar, R., Kamel, N., Naufal, M., Malik, A. S., Dass, S. C., Ahmad, R. F., Abdullah, J. M., & Reza, F. (2018). A study of decoding human brain activities from simultaneous data of EEG and fMRI using MVPA. Australasian Physical & Engineering Sciences in Medicine, 41(3), 633–645. 10.1007/s13246-018-0656-5

