## Supplementary Table 1 for "Multivariate pattern analysis identifies potential inter-trial resting-state EEG biomarkers in fibromyalgia"

**Supplementary Table 1 PROBAST+AI evaluation of risk of bias and applicability assessment**

| | |
|---|---|
| **PICOTS** | |
| Population | Adult women aged 35-65, including 25 FM patients diagnosed using ACR criteria and 26 matched HC. Exclusion criteria included neurological, psychiatric, metabolic, or substance-related conditions. |
| Index model(s) | LSVM and LDA models trained on PSD features from RS-EEG and PV. |
| Comparator model(s) | No external models. Internal comparison was made between LSVM and LDA using identical datasets. |
| Outcome(s) | Binary classification of FM and HC, and STFM and LTFM. |
| Timing | Prediction based on RS-EEG segments with no follow-up. Classification reflects the group status at the time of recording. |
| Setting and intended use of the prediction model | Controlled research setting. Models are exploratory and aim to identify EEG-based features for potential biomarker development in FM. |
| **Classify the assessment based on its aim** | |
| **Type of prediction study** | **Explanation** |
| Combination (Model development and evaluation) | Prediction model development combined in the same study (publication) with the evaluation of its apparent performance, internal validation performance, or external validation performance. |
| **Quality and applicability or risk of bias and applicability** | |
| Publication reference | Current study |
| Models of interest | LSVM and LDA |
| Outcome of interest | (1) FM vs HC classification<br>(2) STFM vs LTFM classification |
| **PROBAST+AI: MODEL DEVELOPMENT** | |
| **DOMAIN 1: Participants and data sources** | |
| A. Quality | Describe the sources of data and criteria for participant selection: Data were collected from 51 women, including 25 FM patients and 26 HC. FM patients were recruited via regional associations in Madrid and diagnosed by rheumatologists following ACR criteria. HC were recruited from the university community. Exclusion criteria included neurological disorders, metabolic alterations, cognitive impairments, substance abuse, and psychiatric disorders. Inclusion criteria included age between 35 and 65 years. The FM and HC groups were matched by age and educational level. Medication use was recorded. EEG data were recorded in a controlled laboratory environment using a 60-electrode system. |
| 1.1 Were appropriate data sources used? | Y |
| 1.2 Was an appropriate study design used? | Y |
| 1.3 Did the in- and exclusions of study participants result in a representative dataset? | PY |
| Concern regarding quality of selection of participants and data sources | QUALITY CONCERN: low |
| Rationale of quality rating | Data sources and participant selection were appropriate and clearly described, with a well-defined case-control design suited for model development. Therefore, the overall concern is low. |
| B. Applicability | Describe included data sources, participants, setting, and dates: The study included 51 adult women aged 35-65 from Madrid, comprising 25 FM patients and 26 HC. Data were collected in a controlled laboratory setting. |
| Concern that the (data of the) included participants do not match the review question or the assessor's intended use of the prediction model | APPLICABILITY CONCERN: low |
| Rationale of applicability rating | Participant selection reflects the typical demographic distribution of FM, which predominantly affects adult women. The sample and setting align well with the intended clinical use of the model, supporting low concern regarding applicability. |
| **DOMAIN 2: Predictors** | |
| A. Quality | List and describe predictors included in the final prediction model, how they were defined and assessed, and their timing of assessment: The predictors included in the model were PSD features extracted from RS-EEG recordings across multiple frequency bands (delta, theta, alpha, beta, gamma) from 60 scalp electrodes, along with psychometric measures of state and |

| | |
|---|---|
| | trait anxiety. EEG data were recorded under standardized conditions using the same equipment and preprocessing pipeline for all participants. PSD values were calculated using the multitaper method and normalized spectrally. Psychometric anxiety scores were self-reported using a standardized questionnaire. |
| 2.1 Were predictors defined and assessed in a similar way for all participants? | Y |
| 2.2 Was any pre-processing of predictors similar for all participants? | Y |
| 2.3 Were predictor assessments made without knowledge of outcome data? | Y |
| 2.4 Were the predictors included in the model available at the time the model was intended to be used? | Y |
| Concern regarding the quality of the predictors or their assessment | QUALITY CONCERN: low |
| Rationale of quality rating | Predictors were consistently defined and assessed using the same EEG protocol and questionnaires for all participants. Preprocessing was uniform, and predictor extraction was independent of outcome data. All predictors were available at the time of intended use, indicating low quality concern. |
| Concern that the definition, pre-processing, assessment, or timing of assessment of the predictors in the model do not match the review question or the assessor's intended use | APPLICABILITY CONCERN: low |
| Rationale of applicability rating | Predictors were defined, pre-processed, and assessed consistently using standard EEG acquisition and psychometric tools applicable to clinical or research settings investigating FM biomarkers. The timing of predictor assessment (RS-EEG and anxiety questionnaires prior to classification) aligns with the intended use of the model, supporting low applicability concern. |
| **DOMAIN 3: Outcome** | |
| A. Quality | Describe the outcome, how it was defined and determined, and the time interval between predictor assessment and outcome determination: The primary outcome was the diagnostic group classification (FM versus HC). FM diagnoses were determined by experienced rheumatologists from hospitals in Madrid, following standardized criteria. HC were screened to ensure the absence of chronic pain conditions or psychiatric and neurological disorders. A secondary outcome involved subdividing FM participants based on symptom duration. Diagnostic duration, reported in months, was used to split the FM group into STFM and LTFM subgroups, with a cut-off at the group's median value of 92 months. The duration data were collected prior to EEG acquisition. EEG recordings (predictors) were obtained during the same session in which clinical and diagnostic information was already known, ensuring that the outcome data preceded the predictor assessment. Therefore, the time interval between outcome assessment and predictor acquisition was immediate and consistent across participants. |
| 3.1 Were outcomes defined and assessed appropriately? | Y |
| 3.2 Were outcomes defined and assessed in a similar way for all participants? | Y |
| 3.3 Were outcome assessments made without use or knowledge of predictor data? | Y |
| 3.4 Was the time interval between predictor assessment and outcome assessment appropriate? | Y |
| Concern regarding quality of the outcome or its determination | QUALITY CONCERN: low |
| Rationale of quality rating | The outcomes were clearly defined using established clinical criteria and uniformly assessed across all participants. The sequence of data collection ensured that outcome determination was made independently of EEG predictor data. The temporal alignment between outcome and predictor assessments was appropriate, resulting in a low concern for quality. |
| B. Applicability | At what time point was the outcome determined: Group classification was determined prior to EEG recording, based on clinical diagnosis and self-reported duration of symptoms.<br><br>If a composite outcome was used, describe the relative frequency/distribution of each contributing outcome: Not applicable; no composite outcome was used. |

| | |
|---|---|
| Concern that the outcome, its definition, assessment, or timing of assessment do not match the review question or the assessor’s intended use | APPLICABILITY CONCERN: low |
| Rationale of applicability rating | The outcome definition (clinical diagnosis of FM and duration of symptoms) aligns with the intended use of the model. Assessments were performed using standard procedures applicable to similar clinical and research settings, supporting low applicability concern. |
| **DOMAIN 4: Analysis** | |
| A. Quality | Describe the numbers of participants, number of candidate predictors, number of outcome events: The study included 51 participants (25 FM, 26 HC). Candidate predictors included PSD values across 60 EEG channels for 10 frequency bands and PV from STAI subscales. The outcome was group membership (FM or HC; STFM or LTFM).<br>Describe how the prediction model was developed (e.g., with respect to modelling technique, predictor selection, and classification or risk group definition):<br>Prediction models were developed using LDA and LSVM, trained on normalized PSD and/or STAI values. Classification focused on FM vs. HC and STFM vs. LTFM. No data-driven predictor selection was applied.<br>Describe the performance measures of the prediction model, e.g., (re)calibration, discrimination, (re)classification, net benefit, and whether they were adjusted for optimism:<br>Model performance was evaluated using accuracy, AUC, recall, precision, F1-score, and calibration. Ten-fold cross-validation was repeated 1000 times to reduce variability and risk of overfitting. Permutation testing was used as a baseline.<br>Describe missing data on predictors and outcomes as well as methods used for handling these missing data:<br>There were no missing data. |
| 4.1 Was there evidence that the sample size was reasonable? | PY |
| 4.2 Were continuous and categorical predictors handled appropriately? | Y |
| 4.3 Were participants with missing or censored data handled appropriately in the analysis? | NA |
| 4.4 If methods to address class imbalance were used, was the model or the model predictions recalibrated?* | NA |
| 4.5 Were methods used to address potential model overfitting? | Y |
| Concern regarding quality of the analysis | QUALITY CONCERN: low |
| Rationale of quality rating | The small sample size relative to the high number of features raises some concern about model stability and overfitting, despite the use of repeated cross-validation. No major class imbalance was present, and all methodological choices were transparently reported. Overall, the concern quality is low. |
| **PROBAST+AI: MODEL EVALUATION** | |
| **DOMAIN 1: Participants and data sources** | |
| A. Risk of bias | Describe the sources of data and criteria for participant selection: Participants were recruited from well-defined clinical associations for FM patients and university community for HC. Inclusion and exclusion criteria were clearly stated, ensuring participants were free from confounding neurological, metabolic, psychiatric, or substance abuse conditions. |
| 1.1 Were appropriate data sources used? | Y |
| 1.2 Was an appropriate study design used? | PY |
| 1.3 Did the in- and exclusions of study participants result in a representative dataset? | Y |
| Risk of bias introduced by the selection of participants and data sources | RISK OF BIAS: low |
| Rationale of risk of bias rating | Data sources were well-defined and traceable, with clear inclusion and exclusion criteria ensuring a representative sample. The prospective observational design was appropriate, and participant selection minimized bias. Therefore, the risk of bias is low. |
| B. Applicability | Describe included data sources, participants, setting, and dates: The study included well-defined participant groups recruited from relevant clinical and community sources within a controlled laboratory setting. Data were collected with clear in/exclusion criteria reflecting the intended clinical population. |

| | |
|---|---|
| Concern that the (data of the) included participants do not match the review question or the assessor’s intended use of the prediction model | APPLICABILITY CONCERN: low |
| Rationale of applicability rating | The participant characteristics, recruitment methods, and study setting closely align with the intended use of the prediction model for distinguishing FM patients from HC. Thus, applicability concern is low. |
| **DOMAIN 2: Predictors** | |
| A. Risk of bias | List and describe predictors included in the evaluated model, e.g., definition and timing of assessment: Predictors included PSD values extracted from RS-EEG across frequency bands, and PV derived from the STAI questionnaire. All EEG data were recorded and preprocessed using standardized protocols for all participants. Psychometric scores were obtained via validated questionnaires administered uniformly. |
| 2.1 Were predictors defined and assessed in a similar way for all participants? | Y |
| 2.2 Was any pre-processing of predictors similar for all participants? | Y |
| 2.3 Were predictor assessments made without knowledge of outcome data? | Y |
| 2.4 Were the predictors included in the model available at the time the model was intended to be used? | Y |
| Risk of bias introduced by the predictors or their assessment | RISK OF BIAS: low |
| Rationale of risk of bias rating | Predictors were clearly defined, consistently assessed, and preprocessed uniformly across all participants. All predictors are available at the time the model would be used, supporting a low risk of bias. |
| Concern that the definition, pre-processing, assessment, or timing of assessment of the predictors in the model do not match the review question or the assessor’s intended use | APPLICABILITY CONCERN: low |
| Rationale of applicability rating | Predictors were consistently defined and measured using standard EEG and anxiety assessments appropriate for the intended use. Preprocessing and timing were uniform, supporting applicability to similar clinical and research settings, yielding low applicability concern. |
| **DOMAIN 3: Outcome** | |
| A. Risk of bias | Describe the outcome, how it was defined and determined, and the time interval between predictor assessment and outcome determination: Outcomes included FM diagnosis based on ACR criteria and anxiety levels assessed via the STAI questionnaire. EEG (predictor) and outcome assessments were conducted during the same session. Standardized methods were applied consistently across participants, and outcome assessors were blinded to EEG data. |
| 3.1 Were outcomes defined and assessed appropriately? | Y |
| 3.2 Were outcomes defined and assessed in a similar way for all participants? | Y |
| 3.3 Were outcome assessments made without use or knowledge of predictor data? | Y |
| 3.4 Was the time interval between predictor assessment and outcome assessment appropriate? | Y |
| Risk of bias introduced by the outcome or its determination | RISK OF BIAS: low |
| Rationale of risk of bias rating | Uniform, validated outcome definitions and assessments, blinded outcome measurement, and appropriate timing minimize risk of bias. Thus, it is assessed as low. |
| B. Applicability | At what time point was the outcome determined: Clinical diagnosis and symptom-duration subgrouping were established before EEG acquisition, during the same visit.<br><br>If a composite outcome was used, describe the relative frequency/distribution of each contributing outcome:<br>Not applicable. Only single binary outcomes were used. |
| Concern that the outcome, its definition, assessment, or timing of assessment do not match the review question or the assessor’s intended use | APPLICABILITY CONCERN: low |
| Rationale of applicability rating | Outcome definitions (ACR criteria and diagnostic duration) and their timing align with the intended use of the model. They can be obtained in routine clinical assessment prior to EEG-based prediction, so no applicability mismatch is present. Thus, the applicability concern is low. |
| **DOMAIN 4: Analysis** | |

| | |
|---|---|
| A. Risk of bias | Describe numbers of participants, number of predictors, outcome events and events per predictor:<br>51 participants (25 FM, 26 HC). Predictors included PSD values per frequency band and STAI scores as PV. Binary outcomes were FM vs HC and LTFM vs STFM.<br>Describe the performance measures of the evaluated model, e.g., (re)calibration, discrimination, (re)classification, net benefit, and whether they were adjusted for optimism:<br>Performance assessed with accuracy, AUC, recall, precision, F1-score, and calibration metrics. Overfitting was controlled via ten-fold cross-validation over 1000 iterations. Permutation testing was used as a data sanity check.<br>Describe any participants who were excluded from the analysis:<br>No participants were excluded.<br>Describe missing data on predictors and outcomes as well as methods used for handling these missing data:<br>No missing data. |
| 4.1 Was model evaluation based on only apparent performance avoided? | Y |
| 4.2 Was there evidence that the sample size was reasonable? | A: PY; I: PY; E: NA |
| 4.3 Were participants with missing or censored data handled appropriately in the analysis? | A: NA; I: NA; E: NA |
| 4.4 If methods to address class imbalance were used, was the evaluation done in a dataset without imbalance correction?* | A: NA; I: NA; E: NA |
| 4.5 If data splitting was done to create training and test datasets, was there evidence that data leakage was avoided?* | A: NA; I: Y; E: NA |
| 4.6 If resampling methods were used to evaluate model performance, were all model development steps replicated in the resampling process?* | A: NA; I: Y; E: NA |
| 4.7 Was the predictive performance of the model evaluated appropriately, e.g., calibration, discrimination, and net benefit? | A: Y; I: Y; E: NA |
| Risk of bias introduced by the analysis | RISK OF BIAS: low |
| Rationale of risk of bias rating | The analysis avoided apparent performance by employing 10-fold cross-validation repeated over 1000 iterations, ensuring performance estimates were not based on the same data used for model development. All model development steps including class balancing, feature selection, and model training were repeated within each cross-validation fold, reducing the risk of optimistic bias. Model performance was evaluated using appropriate discrimination metrics (AUC, F1-score, accuracy, precision, recall) and calibration (intercept and slope). There was no missing data in predictors or outcomes, and no participants were excluded. Although external validation was not performed, the use of rigorous internal validation procedures supports a rating of low risk of bias in the analysis domain. |
| **OVERALL JUDGEMENT OF PREDICTION MODEL** | |
| Overall judgement of quality (development) | QUALITY CONCERN: low |
| Summary of quality concern | All four domains, Participants and Data Sources, Predictors, Outcome, and Analysis, were rated as low concern regarding quality. The study design was appropriate, predictors and outcomes were consistently and clearly defined, and analysis was conducted using robust internal validation techniques. While the sample size was relatively small, methodological safeguards (e.g., repeated cross-validation) mitigated concerns about overfitting. Therefore, overall concern is low. |
| Overall judgement of risk of bias (evaluation) | RISK OF BIAS: low |
| Summary of sources of potential bias | All domains were rated low concern for applicability. The population (middle-aged women with FM) matches the clinical target population, predictors and outcomes align with potential clinical use, and timing and setting of assessments are appropriate. Therefore, the concern that the model development context would misalign with its intended application is low. |
| Overall judgement of applicability (development) | APPLICABILITY CONCERN: low |
| Summary of applicability concern | All four domains were rated low risk of bias. Apparent performance was avoided, internal validation was properly conducted with all model development steps nested within cross-validation folds, and performance was evaluated using multiple appropriate metrics (e.g., AUC, F1-score, calibration slope and intercept). No participants or data were excluded, and there was no missing data. Despite the absence of external validation, internal procedures were sufficiently robust to support a low risk of bias. |

| | |
|---|---|
| Overall judgement of applicability (evaluation) | APPLICABILITY CONCERN: low |
| Summary of applicability concern | All three domains were rated low concern for applicability. The population, predictors, and outcome assessments during evaluation aligned with their counterparts in development and with the intended clinical/research use. The setting was consistent and relevant for future clinical translation, supporting a low concern. |

Abbreviations: FM = Fibromyalgia; ACR = American College of Rheumatology; HC = Healthy control; LSVM = Linear support vector machine; LDA = Linear discriminant analysis; RS = Resting-state; EEG = Electroencephalography; PV: Psychometric values; STFM = Short-term fibromyalgia; LTFM = Long-term fibromyalgia; Y = Yes; PY = Probably yes; NA = Not applicable; PSD = Power spectral density; STAI = State-trait anxiety inventory; AUC = Area under the curve; A = Apparent performance; I = Internal validation or evaluation; E = External validation.
