## Supplementary Table 2 for "Multivariate pattern analysis identifies potential inter-trial resting-state EEG biomarkers in fibromyalgia"

**Supplementary Table 2 Differences in normalized PSD values across frequency bands between FM and HC groups**

| | Delta | Theta | Alpha | Alpha-1 | Alpha-2 | Beta | Beta-1 | Beta-2 | Beta-3 | Gamma |
|---|---|---|---|---|---|---|---|---|---|---|
| Fp1 | 0.0092 | 0.0078 | 0.0041 | 0.0052 | 0.0031 | 0.0007 | 0.0028 | -0.0005 | -0.0005 | 0.0021 |
| Fpz | 0.0099 | 0.0087 | 0.0073 | 0.0075 | 0.0072 | 0.0103 | 0.0084 | 0.0093 | 0.0138 | 0.0181 |
| Fp2 | 0.0112 | 0.0101 | 0.0081 | 0.0088 | 0.0075 | 0.0049 | 0.0055 | 0.0063 | 0.0058 | 0.0107 |
| AF7 | 0.0019 | 0.0002 | -0.0004 | 0.0002 | -0.0008 | -0.0078 | -0.0035 | -0.0078 | -0.0127 | -0.0069 |
| AF3 | -0.0002 | -0.0009 | -0.0017 | -0.0016 | -0.0016 | -0.0098 | -0.0061 | -0.0127 | -0.0115 | -0.0163 |
| AF4 | 0.0068 | 0.0038 | 0.0004 | 0.0008 | -0.0002 | -0.0065 | -0.0039 | -0.0052 | -0.0084 | -0.0106 |
| AF8 | 0.0096 | 0.0094 | 0.0062 | 0.0071 | 0.0054 | 0.0039 | 0.0033 | 0.0024 | 0.0049 | 0.0031 |
| F7 | -0.0047 | -0.0022 | -0.0016 | -0.0015 | -0.0019 | -0.0086 | -0.0048 | -0.0084 | -0.0108 | -0.0102 |
| F5 | -0.0003 | 0.0002 | -0.0003 | 0.0001 | -0.0009 | -0.0109 | -0.0065 | -0.0116 | -0.0135 | -0.0128 |
| F3 | -0.0013 | -0.0018 | -0.0023 | -0.0014 | -0.0029 | -0.0078 | -0.0056 | -0.0078 | -0.0100 | -0.0111 |
| F1 | -0.0022 | -0.0016 | -0.0016 | -0.0012 | -0.0018 | -0.0019 | -0.0020 | -0.0016 | -0.0021 | -0.0025 |
| Fz | -0.0018 | -0.0013 | -0.0016 | -0.0011 | -0.0018 | -0.0014 | -0.0017 | -0.0011 | -0.0012 | -0.0013 |
| F2 | -0.0007 | -0.0009 | -0.0008 | -0.0004 | -0.0011 | -0.0009 | -0.0009 | -0.0001 | -0.0010 | -0.0005 |
| F4 | 0.0005 | -0.0010 | -0.0029 | -0.0018 | -0.0035 | -0.0075 | -0.0068 | -0.0040 | -0.0076 | -0.0100 |
| F6 | 0.0020 | 0.0009 | -0.0004 | 0.0000 | -0.0008 | -0.0049 | -0.0022 | -0.0042 | -0.0070 | -0.0083 |
| F8 | 0.0004 | 0.0002 | 0.0004 | 0.0007 | 0.0002 | -0.0005 | 0.0005 | 0.0001 | -0.0013 | -0.0009 |
| FC5 | -0.0008 | 0.0001 | -0.0006 | -0.0004 | -0.0010 | -0.0056 | -0.0046 | -0.0056 | -0.0069 | -0.0108 |
| FC3 | -0.0015 | -0.0010 | -0.0016 | -0.0010 | -0.0022 | -0.0021 | -0.0025 | -0.0023 | -0.0020 | -0.0041 |
| FC1 | -0.0022 | -0.0016 | -0.0019 | -0.0014 | -0.0024 | -0.0018 | -0.0024 | -0.0017 | -0.0014 | -0.0022 |
| FCz | -0.0021 | -0.0010 | -0.0018 | -0.0012 | -0.0023 | -0.0022 | -0.0029 | -0.0021 | -0.0016 | -0.0018 |
| FC2 | -0.0026 | -0.0020 | -0.0018 | -0.0013 | -0.0022 | -0.0017 | -0.0025 | -0.0014 | -0.0011 | -0.0011 |
| FC4 | -0.0007 | -0.0008 | -0.0012 | -0.0006 | -0.0019 | -0.0019 | -0.0020 | -0.0016 | -0.0018 | -0.0018 |
| FC6 | -0.0002 | -0.0004 | -0.0004 | -0.0004 | -0.0005 | 0.0001 | -0.0003 | -0.0003 | 0.0005 | -0.0006 |
| T7 | -0.0011 | 0.0001 | 0.0030 | 0.0033 | 0.0024 | -0.0080 | -0.0060 | -0.0118 | -0.0082 | -0.0087 |
| C5 | 0.0003 | 0.0001 | 0.0003 | 0.0006 | -0.0001 | -0.0005 | -0.0020 | -0.0005 | 0.0005 | -0.0030 |
| C3 | -0.0013 | -0.0018 | -0.0016 | -0.0009 | -0.0020 | -0.0008 | -0.0025 | 0.0001 | 0.0002 | -0.0015 |
| C1 | -0.0024 | -0.0023 | -0.0022 | -0.0014 | -0.0025 | -0.0011 | -0.0026 | -0.0007 | 0.0000 | -0.0009 |
| Cz | -0.0022 | -0.0013 | -0.0010 | -0.0007 | -0.0012 | -0.0008 | -0.0024 | -0.0007 | 0.0006 | -0.0009 |
| C2 | -0.0016 | -0.0015 | -0.0021 | -0.0019 | -0.0024 | -0.0010 | -0.0022 | -0.0012 | -0.0002 | -0.0003 |
| C4 | -0.0006 | -0.0020 | -0.0040 | -0.0046 | -0.0039 | 0.0000 | -0.0021 | -0.0004 | 0.0018 | 0.0024 |
| C6 | -0.0003 | -0.0006 | -0.0003 | -0.0003 | -0.0005 | 0.0066 | 0.0032 | 0.0069 | 0.0098 | 0.0136 |
| T8 | -0.0012 | -0.0005 | -0.0012 | -0.0003 | -0.0014 | 0.0041 | 0.0015 | 0.0055 | 0.0068 | 0.0095 |
| TP7 | 0.0002 | 0.0008 | 0.0018 | 0.0024 | 0.0013 | -0.0013 | 0.0012 | -0.0007 | -0.0037 | -0.0017 |
| CP5 | -0.0005 | -0.0002 | 0.0000 | -0.0001 | 0.0001 | 0.0005 | -0.0004 | 0.0014 | 0.0008 | -0.0017 |
| CP3 | -0.0003 | -0.0008 | -0.0004 | -0.0004 | -0.0001 | 0.0001 | -0.0010 | 0.0014 | 0.0006 | -0.0003 |
| CP1 | 0.0006 | -0.0006 | -0.0003 | 0.0001 | -0.0002 | -0.0003 | -0.0012 | 0.0002 | 0.0003 | 0.0001 |
| CPz | -0.0016 | -0.0012 | 0.0005 | 0.0006 | 0.0006 | 0.0000 | -0.0009 | -0.0001 | 0.0007 | 0.0002 |
| CP2 | -0.0009 | -0.0008 | 0.0001 | -0.0004 | 0.0004 | 0.0003 | -0.0003 | 0.0002 | 0.0009 | 0.0005 |
| CP4 | -0.0010 | -0.0010 | -0.0008 | -0.0018 | -0.0003 | 0.0010 | 0.0002 | 0.0007 | 0.0016 | 0.0016 |
| CP6 | -0.0002 | -0.0004 | 0.0002 | -0.0011 | 0.0014 | 0.0060 | 0.0054 | 0.0051 | 0.0068 | 0.0110 |
| TP8 | -0.0016 | 0.0002 | 0.0037 | 0.0013 | 0.0062 | 0.0141 | 0.0124 | 0.0134 | 0.0168 | 0.0268 |
| P7 | -0.0005 | -0.0006 | -0.0008 | -0.0011 | -0.0003 | 0.0014 | 0.0015 | 0.0016 | 0.0009 | 0.0018 |
| P5 | 0.0000 | -0.0004 | -0.0006 | -0.0007 | -0.0003 | 0.0019 | 0.0010 | 0.0024 | 0.0021 | 0.0015 |
| P3 | -0.0006 | -0.0006 | -0.0014 | -0.0018 | -0.0012 | 0.0009 | 0.0004 | 0.0015 | 0.0010 | 0.0005 |
| P1 | -0.0015 | -0.0014 | -0.0016 | -0.0023 | -0.0015 | 0.0002 | -0.0002 | 0.0004 | 0.0004 | 0.0002 |
| Pz | -0.0015 | -0.0013 | -0.0008 | -0.0015 | -0.0007 | 0.0004 | 0.0000 | 0.0002 | 0.0006 | 0.0003 |
| P2 | -0.0019 | -0.0013 | -0.0016 | -0.0026 | -0.0016 | 0.0005 | 0.0000 | 0.0002 | 0.0008 | 0.0005 |
| P4 | -0.0012 | -0.0008 | 0.0008 | -0.0003 | 0.0016 | 0.0022 | 0.0022 | 0.0018 | 0.0018 | 0.0011 |
| P6 | -0.0020 | -0.0034 | -0.0014 | -0.0028 | 0.0000 | 0.0035 | 0.0032 | 0.0027 | 0.0035 | 0.0047 |
| P8 | -0.0018 | -0.0006 | -0.0002 | -0.0006 | 0.0010 | 0.0052 | 0.0040 | 0.0047 | 0.0063 | 0.0075 |
| PO7 | 0.0012 | 0.0008 | 0.0015 | 0.0016 | 0.0018 | 0.0031 | 0.0037 | 0.0027 | 0.0018 | 0.0011 |
| PO5 | -0.0013 | -0.0016 | -0.0006 | -0.0010 | -0.0003 | 0.0022 | 0.0025 | 0.0020 | 0.0011 | 0.0006 |
| PO3 | -0.0005 | -0.0005 | -0.0003 | -0.0013 | -0.0003 | 0.0014 | 0.0020 | 0.0014 | 0.0006 | -0.0001 |
| POz | -0.0016 | -0.0009 | -0.0007 | -0.0014 | -0.0010 | 0.0009 | 0.0011 | 0.0010 | 0.0006 | -0.0002 |
| PO4 | -0.0021 | -0.0010 | 0.0000 | -0.0011 | 0.0002 | 0.0009 | 0.0009 | 0.0012 | 0.0006 | -0.0001 |

| | | | | | | | | | | |
|---|---|---|---|---|---|---|---|---|---|---|
| PO6 | -0.0014 | -0.0009 | 0.0007 | 0.0001 | 0.0015 | 0.0033 | 0.0025 | 0.0034 | 0.0034 | 0.0030 |
| PO8 | -0.0002 | 0.0003 | 0.0021 | 0.0016 | 0.0032 | 0.0111 | 0.0070 | 0.0104 | 0.0139 | 0.0157 |
| O1 | 0.0001 | 0.0006 | 0.0027 | 0.0024 | 0.0031 | 0.0024 | 0.0040 | 0.0018 | 0.0004 | -0.0017 |
| Oz | 0.0024 | 0.0019 | 0.0018 | 0.0019 | 0.0018 | 0.0018 | 0.0025 | 0.0019 | 0.0009 | -0.0007 |
| O2 | -0.0003 | 0.0005 | 0.0013 | 0.0015 | 0.0013 | 0.0017 | 0.0019 | 0.0022 | 0.0008 | -0.0026 |
