## Supplementary Table 3 for "Multivariate pattern analysis identifies potential inter-trial resting-state EEG biomarkers in fibromyalgia"

**Supplementary Table 3 Differences in normalized PSD values across frequency bands between STFM and LTFM groups**

| | Delta | Theta | Alpha | Alpha-1 | Alpha-2 | Beta | Beta-1 | Beta-2 | Beta-3 | Gamma |
|---|---|---|---|---|---|---|---|---|---|---|
| Fp1 | 0.0045 | 0.0036 | 0.0067 | 0.0054 | 0.0082 | 0.0025 | 0.0037 | 0.0027 | -0.0001 | 0.0084 |
| Fpz | 0.0052 | 0.0060 | 0.0030 | 0.0029 | 0.0034 | 0.0034 | 0.0036 | 0.0045 | 0.0026 | 0.0015 |
| Fp2 | 0.0031 | 0.0043 | 0.0020 | 0.0014 | 0.0030 | 0.0116 | 0.0086 | 0.0130 | 0.0109 | 0.0094 |
| AF7 | 0.0027 | 0.0014 | 0.0030 | 0.0018 | 0.0040 | 0.0071 | 0.0060 | 0.0051 | 0.0066 | 0.0062 |
| AF3 | 0.0011 | 0.0006 | 0.0008 | 0.0010 | 0.0006 | -0.0046 | 0.0061 | 0.0049 | -0.0161 | 0.0032 |
| AF4 | 0.0013 | 0.0030 | 0.0028 | 0.0022 | 0.0035 | 0.0041 | 0.0063 | 0.0002 | 0.0033 | 0.0136 |
| AF8 | 0.0027 | 0.0025 | 0.0017 | 0.0010 | 0.0027 | 0.0098 | 0.0093 | 0.0094 | 0.0140 | 0.0224 |
| F7 | 0.0048 | 0.0014 | 0.0004 | 0.0014 | -0.0012 | -0.0156 | -0.0071 | -0.0130 | -0.0195 | -0.0173 |
| F5 | 0.0021 | 0.0004 | -0.0009 | -0.0002 | -0.0021 | -0.0126 | -0.0044 | -0.0073 | -0.0165 | -0.0121 |
| F3 | 0.0018 | 0.0025 | 0.0007 | 0.0018 | -0.0008 | -0.0061 | -0.0020 | -0.0041 | -0.0093 | -0.0087 |
| F1 | 0.0030 | 0.0035 | 0.0031 | 0.0035 | 0.0021 | 0.0018 | 0.0015 | 0.0019 | 0.0020 | 0.0011 |
| Fz | 0.0023 | 0.0035 | 0.0031 | 0.0033 | 0.0023 | 0.0034 | 0.0024 | 0.0034 | 0.0038 | 0.0029 |
| F2 | 0.0020 | 0.0032 | 0.0028 | 0.0028 | 0.0022 | 0.0034 | 0.0023 | 0.0035 | 0.0039 | 0.0033 |
| F4 | 0.0014 | 0.0030 | 0.0046 | 0.0031 | 0.0052 | 0.0060 | 0.0087 | 0.0041 | 0.0063 | 0.0160 |
| F6 | 0.0011 | 0.0006 | 0.0012 | 0.0007 | 0.0016 | 0.0035 | 0.0038 | 0.0027 | 0.0066 | 0.0071 |
| F8 | -0.0012 | -0.0013 | 0.0000 | 0.0000 | -0.0001 | 0.0018 | 0.0018 | 0.0000 | 0.0035 | 0.0037 |
| FC5 | 0.0055 | 0.0014 | -0.0005 | -0.0004 | -0.0011 | -0.0100 | -0.0055 | -0.0124 | -0.0129 | -0.0192 |
| FC3 | -0.0019 | 0.0006 | 0.0007 | 0.0007 | 0.0003 | 0.0010 | 0.0014 | 0.0006 | 0.0011 | 0.0001 |
| FC1 | 0.0003 | 0.0012 | 0.0010 | 0.0012 | 0.0003 | 0.0022 | 0.0017 | 0.0022 | 0.0027 | 0.0016 |
| FCz | 0.0007 | 0.0026 | 0.0025 | 0.0027 | 0.0019 | 0.0052 | 0.0039 | 0.0061 | 0.0058 | 0.0043 |
| FC2 | 0.0025 | 0.0033 | 0.0024 | 0.0025 | 0.0018 | 0.0042 | 0.0033 | 0.0043 | 0.0047 | 0.0032 |
| FC4 | -0.0003 | 0.0001 | 0.0000 | 0.0000 | -0.0005 | 0.0017 | 0.0011 | 0.0024 | 0.0023 | 0.0013 |
| FC6 | -0.0013 | -0.0017 | -0.0031 | -0.0032 | -0.0031 | -0.0046 | -0.0030 | -0.0021 | -0.0053 | -0.0053 |
| T7 | 0.0019 | -0.0014 | -0.0011 | -0.0014 | -0.0011 | -0.0152 | -0.0159 | -0.0225 | -0.0125 | -0.0276 |
| C5 | 0.0006 | -0.0004 | 0.0017 | 0.0008 | 0.0023 | -0.0034 | -0.0020 | -0.0030 | -0.0060 | -0.0122 |
| C3 | 0.0017 | 0.0006 | 0.0030 | 0.0015 | 0.0040 | 0.0015 | 0.0026 | 0.0022 | -0.0004 | -0.0027 |
| C1 | -0.0027 | -0.0020 | 0.0003 | -0.0008 | 0.0011 | 0.0018 | 0.0017 | 0.0017 | 0.0020 | 0.0016 |
| Cz | -0.0008 | 0.0005 | 0.0013 | 0.0011 | 0.0014 | 0.0037 | 0.0027 | 0.0049 | 0.0040 | 0.0040 |
| C2 | -0.0005 | 0.0001 | 0.0021 | 0.0016 | 0.0026 | 0.0034 | 0.0035 | 0.0036 | 0.0030 | 0.0023 |
| C4 | -0.0006 | -0.0017 | 0.0003 | -0.0006 | 0.0011 | 0.0025 | 0.0031 | 0.0026 | 0.0014 | 0.0000 |
| C6 | 0.0003 | -0.0009 | -0.0002 | -0.0005 | 0.0000 | -0.0006 | -0.0003 | -0.0011 | -0.0019 | -0.0049 |
| T8 | 0.0021 | 0.0001 | -0.0011 | 0.0016 | -0.0028 | -0.0108 | -0.0081 | -0.0076 | -0.0125 | -0.0121 |
| TP7 | -0.0007 | -0.0015 | -0.0025 | -0.0020 | -0.0032 | -0.0115 | -0.0089 | -0.0133 | -0.0149 | -0.0242 |
| CP5 | 0.0029 | 0.0015 | 0.0023 | 0.0021 | 0.0019 | 0.0004 | 0.0011 | 0.0018 | -0.0016 | -0.0071 |
| CP3 | -0.0016 | -0.0012 | 0.0009 | 0.0004 | 0.0010 | 0.0021 | 0.0017 | 0.0035 | 0.0012 | -0.0005 |
| CP1 | -0.0016 | -0.0019 | -0.0009 | -0.0012 | -0.0004 | 0.0008 | -0.0004 | 0.0016 | 0.0016 | 0.0009 |
| CPz | -0.0018 | -0.0017 | -0.0008 | -0.0007 | -0.0002 | 0.0010 | -0.0004 | 0.0015 | 0.0020 | 0.0015 |
| CP2 | -0.0018 | -0.0014 | 0.0000 | 0.0002 | 0.0006 | 0.0013 | 0.0006 | 0.0011 | 0.0016 | 0.0011 |
| CP4 | -0.0019 | -0.0017 | 0.0003 | 0.0005 | 0.0005 | 0.0018 | 0.0013 | 0.0015 | 0.0016 | 0.0007 |
| CP6 | -0.0001 | -0.0006 | 0.0010 | 0.0008 | 0.0013 | 0.0022 | 0.0014 | 0.0020 | 0.0025 | 0.0007 |
| TP8 | 0.0034 | 0.0005 | 0.0008 | 0.0010 | 0.0011 | -0.0019 | -0.0019 | -0.0022 | -0.0016 | -0.0006 |
| P7 | 0.0004 | -0.0016 | -0.0027 | -0.0036 | -0.0020 | 0.0020 | 0.0011 | 0.0000 | 0.0025 | 0.0026 |
| P5 | 0.0006 | 0.0000 | -0.0002 | 0.0000 | -0.0003 | 0.0020 | 0.0009 | 0.0017 | 0.0023 | 0.0008 |
| P3 | -0.0005 | -0.0001 | 0.0015 | 0.0020 | 0.0012 | 0.0024 | 0.0013 | 0.0028 | 0.0022 | 0.0011 |
| P1 | -0.0019 | -0.0011 | 0.0003 | 0.0012 | 0.0002 | 0.0007 | -0.0010 | 0.0011 | 0.0017 | 0.0012 |
| Pz | -0.0032 | -0.0017 | -0.0013 | -0.0002 | -0.0013 | 0.0005 | -0.0013 | 0.0006 | 0.0016 | 0.0013 |
| P2 | -0.0031 | -0.0017 | -0.0002 | 0.0012 | -0.0004 | 0.0006 | -0.0012 | 0.0007 | 0.0017 | 0.0015 |
| P4 | -0.0015 | 0.0001 | -0.0005 | -0.0001 | -0.0011 | 0.0015 | -0.0001 | 0.0011 | 0.0027 | 0.0026 |
| P6 | -0.0104 | -0.0091 | -0.0048 | -0.0053 | -0.0044 | -0.0027 | -0.0030 | -0.0033 | -0.0028 | -0.0030 |
| P8 | 0.0035 | 0.0003 | -0.0017 | -0.0011 | -0.0016 | -0.0001 | -0.0027 | -0.0006 | 0.0024 | 0.0032 |
| PO7 | -0.0016 | -0.0022 | -0.0058 | -0.0061 | -0.0053 | -0.0016 | -0.0034 | -0.0028 | 0.0002 | -0.0021 |

| | | | | | | | | | | |
|---|---|---|---|---|---|---|---|---|---|---|
| PO5 | -0.0008 | -0.0006 | -0.0044 | -0.0038 | -0.0045 | -0.0012 | -0.0033 | -0.0020 | 0.0005 | 0.0000 |
| PO3 | -0.0038 | -0.0020 | -0.0025 | -0.0018 | -0.0029 | -0.0008 | -0.0033 | -0.0011 | 0.0013 | 0.0020 |
| POz | -0.0051 | -0.0030 | -0.0035 | -0.0031 | -0.0038 | -0.0007 | -0.0033 | -0.0010 | 0.0014 | 0.0022 |
| PO4 | -0.0048 | -0.0024 | -0.0038 | -0.0033 | -0.0044 | 0.0000 | -0.0029 | -0.0006 | 0.0023 | 0.0028 |
| PO6 | -0.0038 | -0.0027 | -0.0035 | -0.0034 | -0.0032 | -0.0008 | -0.0039 | -0.0018 | 0.0012 | 0.0007 |
| PO8 | -0.0038 | -0.0032 | -0.0037 | -0.0035 | -0.0034 | -0.0007 | -0.0036 | -0.0017 | 0.0013 | 0.0013 |
| O1 | 0.0002 | -0.0002 | -0.0041 | -0.0046 | -0.0037 | -0.0016 | -0.0026 | -0.0035 | 0.0000 | 0.0010 |
| Oz | -0.0012 | -0.0007 | -0.0022 | -0.0022 | -0.0023 | 0.0002 | -0.0017 | -0.0007 | 0.0022 | 0.0040 |
| O2 | -0.0017 | -0.0007 | -0.0022 | -0.0024 | -0.0020 | 0.0018 | -0.0014 | 0.0007 | 0.0052 | 0.0091 |
