## Supplementary Table 4 for "Multivariate pattern analysis identifies potential inter-trial resting-state EEG biomarkers in fibromyalgia"

**Supplementary Table 4 Comparison of LSVM and LDA model performance across frequency bands**

| Frequency | Model | Permuted | ACC | ACC SD | AUC | AUC SD | Precision | Precision SD | Recall | Recall SD | F1-Score | F1-Score SD | Intercept | Intercept SD | Slope | Slope SD |
|---|---|---|---|---|---|---|---|---|---|---|---|---|---|---|---|---|
| Delta | LSVM | Yes | 0.511 | 0.004 | 0.512 | 0.004 | 0.676 | 0.007 | 0.516 | 0.003 | 0.585 | 0.004 | 0.030 | 0.005 | 0.063 | 0.022 |
| | | No | 0.715 | 0.002 | 0.779 | 0.001 | 0.696 | 0.003 | 0.733 | 0.002 | 0.714 | 0.002 | 0.036 | 0.002 | 0.988 | 0.008 |
| | LDA | Yes | 0.511 | 0.004 | 0.512 | 0.004 | 0.601 | 0.006 | 0.518 | 0.003 | 0.556 | 0.004 | 0.036 | 0.002 | 0.168 | 0.044 |
| | | No | 0.703 | 0.002 | 0.772 | 0.001 | 0.684 | 0.003 | 0.720 | 0.002 | 0.701 | 0.002 | 0.031 | 0.002 | 1.017 | 0.011 |
| Theta | LSVM | Yes | 0.517 | 0.004 | 0.521 | 0.003 | 0.652 | 0.006 | 0.522 | 0.003 | 0.580 | 0.004 | 0.020 | 0.004 | 0.131 | 0.020 |
| | | No | 0.738 | 0.002 | 0.809 | 0.001 | 0.721 | 0.002 | 0.755 | 0.002 | 0.738 | 0.002 | 0.042 | 0.002 | 1.144 | 0.006 |
| | LDA | Yes | 0.513 | 0.004 | 0.515 | 0.003 | 0.587 | 0.005 | 0.521 | 0.003 | 0.552 | 0.004 | 0.034 | 0.002 | 0.219 | 0.043 |
| | | No | 0.735 | 0.002 | 0.806 | 0.001 | 0.735 | 0.003 | 0.743 | 0.002 | 0.739 | 0.002 | 0.012 | 0.002 | 1.008 | 0.006 |
| Alpha | LSVM | Yes | 0.509 | 0.004 | 0.507 | 0.004 | 0.658 | 0.006 | 0.515 | 0.003 | 0.578 | 0.004 | 0.036 | 0.004 | 0.037 | 0.023 |
| | | No | 0.773 | 0.001 | 0.853 | 0.001 | 0.811 | 0.002 | 0.761 | 0.002 | 0.785 | 0.001 | -0.067 | 0.003 | 1.391 | 0.006 |
| | LDA | Yes | 0.503 | 0.004 | 0.499 | 0.004 | 0.593 | 0.006 | 0.511 | 0.004 | 0.549 | 0.004 | 0.045 | 0.002 | -0.031 | 0.054 |
| | | No | 0.773 | 0.001 | 0.856 | 0.001 | 0.810 | 0.002 | 0.761 | 0.001 | 0.785 | 0.001 | -0.059 | 0.002 | 1.010 | 0.004 |
| Alpha-1 | LSVM | Yes | 0.505 | 0.004 | 0.498 | 0.004 | 0.630 | 0.007 | 0.513 | 0.003 | 0.565 | 0.004 | 0.045 | 0.004 | -0.012 | 0.024 |
| | | No | 0.747 | 0.002 | 0.831 | 0.001 | 0.788 | 0.002 | 0.736 | 0.002 | 0.761 | 0.002 | -0.047 | 0.003 | 1.286 | 0.006 |
| | LDA | Yes | 0.502 | 0.004 | 0.499 | 0.004 | 0.578 | 0.006 | 0.511 | 0.004 | 0.542 | 0.004 | 0.045 | 0.002 | -0.035 | 0.054 |
| | | No | 0.746 | 0.002 | 0.833 | 0.001 | 0.786 | 0.002 | 0.735 | 0.002 | 0.760 | 0.002 | -0.043 | 0.002 | 1.014 | 0.005 |
| Alpha-2 | LSVM | Yes | 0.495 | 0.005 | 0.485 | 0.004 | 0.651 | 0.008 | 0.504 | 0.003 | 0.568 | 0.005 | 0.065 | 0.005 | -0.110 | 0.027 |
| | | No | 0.759 | 0.002 | 0.832 | 0.001 | 0.808 | 0.002 | 0.743 | 0.002 | 0.774 | 0.002 | -0.095 | 0.002 | 1.302 | 0.005 |
| | LDA | Yes | 0.492 | 0.004 | 0.485 | 0.004 | 0.583 | 0.006 | 0.502 | 0.004 | 0.540 | 0.004 | 0.055 | 0.003 | -0.260 | 0.066 |
| | | No | 0.754 | 0.002 | 0.834 | 0.001 | 0.789 | 0.002 | 0.745 | 0.002 | 0.767 | 0.002 | -0.048 | 0.002 | 1.015 | 0.005 |
| Beta | LSVM | Yes | 0.497 | 0.005 | 0.493 | 0.004 | 0.628 | 0.009 | 0.506 | 0.004 | 0.561 | 0.005 | 0.052 | 0.005 | -0.054 | 0.027 |
| | | No | 0.871 | 0.001 | 0.932 | 0.000 | 0.865 | 0.002 | 0.880 | 0.002 | 0.872 | 0.001 | 0.020 | 0.003 | 1.742 | 0.007 |
| | LDA | Yes | 0.500 | 0.005 | 0.496 | 0.004 | 0.572 | 0.006 | 0.509 | 0.004 | 0.539 | 0.005 | 0.048 | 0.003 | -0.112 | 0.059 |
| | | No | 0.862 | 0.001 | 0.931 | 0.000 | 0.860 | 0.002 | 0.869 | 0.002 | 0.864 | 0.001 | 0.041 | 0.003 | 1.015 | 0.004 |
| Beta-1 | LSVM | Yes | 0.495 | 0.004 | 0.481 | 0.005 | 0.703 | 0.009 | 0.504 | 0.003 | 0.587 | 0.005 | 0.081 | 0.007 | -0.149 | 0.029 |
| | | No | 0.789 | 0.002 | 0.871 | 0.001 | 0.794 | 0.002 | 0.794 | 0.002 | 0.794 | 0.002 | -0.005 | 0.003 | 1.410 | 0.006 |
| | LDA | Yes | 0.490 | 0.005 | 0.479 | 0.004 | 0.590 | 0.007 | 0.500 | 0.004 | 0.541 | 0.005 | 0.062 | 0.003 | -0.425 | 0.072 |
| | | No | 0.789 | 0.002 | 0.873 | 0.001 | 0.800 | 0.002 | 0.790 | 0.002 | 0.795 | 0.002 | 0.011 | 0.002 | 1.029 | 0.006 |
| Beta-2 | LSVM | Yes | 0.512 | 0.004 | 0.513 | 0.004 | 0.601 | 0.008 | 0.520 | 0.004 | 0.557 | 0.005 | 0.032 | 0.003 | 0.085 | 0.023 |
| | | No | 0.791 | 0.001 | 0.867 | 0.001 | 0.805 | 0.002 | 0.789 | 0.002 | 0.797 | 0.001 | -0.045 | 0.002 | 1.348 | 0.006 |

| | | | | | | | | | | | | | | | |
|---|---|---|---|---|---|---|---|---|---|---|---|---|---|---|---|
| | LDA | Yes | 0.516 | 0.004 | 0.520 | 0.003 | 0.574 | 0.006 | 0.524 | 0.003 | 0.548 | 0.004 | 0.032 | 0.002 | 0.256 | 0.040 |
| | | No | 0.783 | 0.002 | 0.862 | 0.001 | 0.791 | 0.002 | 0.785 | 0.002 | 0.788 | 0.002 | 0.020 | 0.002 | 1.008 | 0.005 |
| Beta-3 | LSVM | Yes | 0.501 | 0.005 | 0.494 | 0.004 | 0.674 | 0.008 | 0.508 | 0.003 | 0.580 | 0.005 | 0.054 | 0.006 | -0.047 | 0.025 |
| | | No | 0.876 | 0.001 | 0.930 | 0.000 | 0.864 | 0.002 | 0.891 | 0.001 | 0.877 | 0.001 | 0.012 | 0.003 | 1.635 | 0.006 |
| | LDA | Yes | 0.499 | 0.004 | 0.493 | 0.004 | 0.596 | 0.006 | 0.509 | 0.004 | 0.549 | 0.004 | 0.050 | 0.003 | -0.144 | 0.059 |
| | | No | 0.861 | 0.001 | 0.930 | 0.000 | 0.853 | 0.002 | 0.873 | 0.002 | 0.863 | 0.001 | 0.098 | 0.003 | 0.951 | 0.003 |
| Gamma | LSVM | Yes | 0.509 | 0.004 | 0.507 | 0.004 | 0.662 | 0.007 | 0.515 | 0.003 | 0.579 | 0.004 | 0.034 | 0.005 | 0.046 | 0.022 |
| | | No | 0.905 | 0.001 | 0.963 | 0.000 | 0.881 | 0.002 | 0.929 | 0.001 | 0.904 | 0.001 | 0.190 | 0.005 | 2.121 | 0.011 |
| | LDA | Yes | 0.509 | 0.004 | 0.506 | 0.004 | 0.599 | 0.006 | 0.517 | 0.004 | 0.555 | 0.004 | 0.040 | 0.002 | 0.077 | 0.051 |
| | | No | 0.888 | 0.001 | 0.956 | 0.000 | 0.867 | 0.002 | 0.910 | 0.001 | 0.888 | 0.001 | 0.233 | 0.004 | 0.976 | 0.004 |

Abbreviations: LSVM = Linear support vector machine; LDA = Linear discriminant analysis; ACC = Accuracy; SD = Standard deviation; AUC = Area under the curve.
