## Supplementary Table 5 for "Multivariate pattern analysis identifies potential inter-trial resting-state EEG biomarkers in fibromyalgia"

**Supplementary Table 5 Performance metrics of the LDA model across PV and frequency bands for the comparison between FM and HC groups.**

| Frequency | Feature | Permuted | ACC | ACC SD | AUC | AUC SD | Precision | Precision SD | Recall | Recall STD | F1-Score | F1-Score SD | Intercept | Intercept SD | Slope | Slope SD |
|---|---|---|---|---|---|---|---|---|---|---|---|---|---|---|---|---|
| | STAI-S | Yes | 0.506 | 0.004 | 0.509 | 0.002 | 0.358 | 0.016 | 0.493 | 0.005 | 0.414 | 0.012 | -0.011 | 0.005 | 0.741 | 0.114 |
| | | No | 0.822 | 0.000 | 0.911 | 0.000 | 0.759 | 0.000 | 0.862 | 0.000 | 0.807 | 0.000 | 0.205 | 0.000 | 0.769 | 0.000 |
| | STAI-T | Yes | 0.511 | 0.001 | 0.478 | 0.006 | 0.001 | 0.003 | - | - | - | - | -0.389 | 0.021 | -8.077 | 0.496 |
| | | No | 0.802 | 0.000 | 0.923 | 0.000 | 0.879 | 0.000 | 0.757 | 0.000 | 0.813 | 0.000 | -0.296 | 0.000 | 0.814 | 0.000 |
| | STAI-S + STAI-T | Yes | 0.508 | 0.003 | 0.481 | 0.006 | 0.014 | 0.011 | - | - | - | - | -0.197 | 0.035 | -3.600 | 0.811 |
| | | No | 0.862 | 0.000 | 0.943 | 0.000 | 0.920 | 0.000 | 0.820 | 0.000 | 0.867 | 0.000 | -0.149 | 0.000 | 0.773 | 0.000 |
| Delta | PSD | Yes | 0.511 | 0.004 | 0.512 | 0.004 | 0.601 | 0.006 | 0.518 | 0.003 | 0.556 | 0.004 | 0.036 | 0.002 | 0.168 | 0.044 |
| | | No | 0.703 | 0.002 | 0.772 | 0.001 | 0.684 | 0.003 | 0.720 | 0.002 | 0.701 | 0.002 | 0.031 | 0.002 | 1.017 | 0.011 |
| | PSD + STAI-S | Yes | 0.510 | 0.004 | 0.511 | 0.004 | 0.600 | 0.006 | 0.518 | 0.004 | 0.556 | 0.004 | 0.036 | 0.002 | 0.156 | 0.044 |
| | | No | 0.866 | 0.001 | 0.952 | 0.000 | 0.914 | 0.001 | 0.838 | 0.001 | 0.875 | 0.001 | -0.260 | 0.004 | 0.737 | 0.004 |
| | PSD + STAI-T | Yes | 0.492 | 0.005 | 0.483 | 0.004 | 0.594 | 0.007 | 0.502 | 0.004 | 0.544 | 0.005 | 0.052 | 0.003 | -0.211 | 0.061 |
| | | No | 0.881 | 0.001 | 0.958 | 0.000 | 0.871 | 0.001 | 0.893 | 0.001 | 0.882 | 0.001 | 0.209 | 0.003 | 0.687 | 0.003 |
| | PSD + STAI-S + STAI-T | Yes | 0.501 | 0.004 | 0.501 | 0.004 | 0.582 | 0.006 | 0.510 | 0.004 | 0.544 | 0.004 | 0.043 | 0.002 | 0.016 | 0.038 |
| | | No | 0.918 | 0.001 | 0.977 | 0.000 | 0.911 | 0.001 | 0.927 | 0.001 | 0.919 | 0.001 | 0.047 | 0.005 | 0.688 | 0.004 |
| Theta | PSD | Yes | 0.513 | 0.004 | 0.515 | 0.003 | 0.587 | 0.005 | 0.521 | 0.003 | 0.552 | 0.004 | 0.034 | 0.002 | 0.219 | 0.043 |
| | | No | 0.735 | 0.002 | 0.806 | 0.001 | 0.735 | 0.003 | 0.743 | 0.002 | 0.739 | 0.002 | 0.012 | 0.002 | 1.008 | 0.006 |
| | PSD + STAI-S | Yes | 0.512 | 0.004 | 0.514 | 0.003 | 0.586 | 0.005 | 0.520 | 0.003 | 0.551 | 0.004 | 0.034 | 0.002 | 0.216 | 0.044 |
| | | No | 0.881 | 0.001 | 0.952 | 0.000 | 0.913 | 0.001 | 0.863 | 0.001 | 0.887 | 0.001 | -0.255 | 0.003 | 0.732 | 0.003 |
| | PSD + STAI-T | Yes | 0.488 | 0.005 | 0.478 | 0.004 | 0.573 | 0.006 | 0.499 | 0.004 | 0.533 | 0.005 | 0.060 | 0.003 | -0.403 | 0.075 |
| | | No | 0.917 | 0.001 | 0.962 | 0.000 | 0.902 | 0.001 | 0.934 | 0.001 | 0.917 | 0.001 | 0.202 | 0.003 | 0.698 | 0.002 |
| | PSD + STAI-S + STAI-T | Yes | 0.501 | 0.004 | 0.498 | 0.004 | 0.576 | 0.006 | 0.510 | 0.004 | 0.541 | 0.004 | 0.047 | 0.002 | -0.088 | 0.054 |
| | | No | 0.931 | 0.001 | 0.981 | 0.000 | 0.915 | 0.001 | 0.948 | 0.001 | 0.931 | 0.001 | 0.100 | 0.005 | 0.695 | 0.003 |
| Alpha | PSD | Yes | 0.503 | 0.004 | 0.499 | 0.004 | 0.593 | 0.006 | 0.511 | 0.004 | 0.549 | 0.004 | 0.045 | 0.002 | -0.031 | 0.054 |
| | | No | 0.773 | 0.001 | 0.856 | 0.001 | 0.810 | 0.002 | 0.761 | 0.001 | 0.785 | 0.001 | -0.059 | 0.002 | 1.010 | 0.004 |
| | PSD + | Yes | 0.502 | 0.004 | 0.498 | 0.004 | 0.592 | 0.006 | 0.511 | 0.004 | 0.549 | 0.004 | 0.045 | 0.002 | -0.043 | 0.055 |

| | | | | | | | | | | | | | | | | |
|---|---|---|---|---|---|---|---|---|---|---|---|---|---|---|---|---|
| | STAI-S | | | | | | | | | | | | | | | |
| | | No | 0.897 | 0.001 | 0.968 | 0.000 | 0.920 | 0.001 | 0.883 | 0.001 | 0.901 | 0.001 | -0.287 | 0.003 | 0.742 | 0.002 |
| | PSD + STAI-T | Yes | 0.503 | 0.004 | 0.497 | 0.004 | 0.581 | 0.006 | 0.512 | 0.004 | 0.544 | 0.004 | 0.044 | 0.002 | -0.013 | 0.055 |
| | | No | 0.909 | 0.001 | 0.973 | 0.000 | 0.909 | 0.001 | 0.913 | 0.001 | 0.911 | 0.001 | 0.166 | 0.003 | 0.759 | 0.002 |
| | PSD + STAI-S + STAI-T | Yes | 0.500 | 0.004 | 0.494 | 0.004 | 0.606 | 0.006 | 0.509 | 0.004 | 0.553 | 0.004 | 0.049 | 0.003 | -0.142 | 0.059 |
| | | No | 0.935 | 0.001 | 0.986 | 0.000 | 0.924 | 0.001 | 0.947 | 0.001 | 0.935 | 0.001 | 0.102 | 0.004 | 0.729 | 0.002 |
| Alpha-1 | PSD | Yes | 0.502 | 0.004 | 0.499 | 0.004 | 0.578 | 0.006 | 0.511 | 0.004 | 0.542 | 0.004 | 0.045 | 0.002 | -0.035 | 0.054 |
| | | No | 0.746 | 0.002 | 0.833 | 0.001 | 0.786 | 0.002 | 0.735 | 0.002 | 0.760 | 0.002 | -0.043 | 0.002 | 1.014 | 0.005 |
| | PSD + STAI-S | Yes | 0.502 | 0.004 | 0.499 | 0.004 | 0.577 | 0.006 | 0.511 | 0.004 | 0.542 | 0.004 | 0.045 | 0.002 | -0.041 | 0.053 |
| | | No | 0.893 | 0.001 | 0.959 | 0.000 | 0.920 | 0.001 | 0.876 | 0.001 | 0.898 | 0.001 | -0.261 | 0.003 | 0.730 | 0.003 |
| | PSD + STAI-T | Yes | 0.492 | 0.005 | 0.486 | 0.004 | 0.571 | 0.006 | 0.502 | 0.004 | 0.535 | 0.005 | 0.052 | 0.003 | -0.208 | 0.064 |
| | | No | 0.906 | 0.001 | 0.964 | 0.000 | 0.897 | 0.001 | 0.918 | 0.001 | 0.907 | 0.001 | 0.208 | 0.003 | 0.743 | 0.003 |
| | PSD + STAI-S + STAI-T | Yes | 0.514 | 0.004 | 0.515 | 0.004 | 0.583 | 0.005 | 0.522 | 0.004 | 0.551 | 0.004 | 0.037 | 0.002 | 0.134 | 0.045 |
| | | No | 0.929 | 0.001 | 0.980 | 0.000 | 0.913 | 0.001 | 0.947 | 0.001 | 0.929 | 0.001 | 0.118 | 0.004 | 0.703 | 0.003 |
| Alpha-2 | PSD | Yes | 0.492 | 0.004 | 0.485 | 0.004 | 0.583 | 0.006 | 0.502 | 0.004 | 0.540 | 0.004 | 0.055 | 0.003 | -0.260 | 0.066 |
| | | No | 0.754 | 0.002 | 0.834 | 0.001 | 0.789 | 0.002 | 0.745 | 0.002 | 0.767 | 0.002 | -0.048 | 0.002 | 1.015 | 0.005 |
| | PSD + STAI-S | Yes | 0.491 | 0.005 | 0.484 | 0.004 | 0.582 | 0.006 | 0.501 | 0.004 | 0.539 | 0.005 | 0.055 | 0.003 | -0.275 | 0.066 |
| | | No | 0.883 | 0.001 | 0.962 | 0.000 | 0.919 | 0.001 | 0.861 | 0.001 | 0.889 | 0.001 | -0.309 | 0.003 | 0.759 | 0.002 |
| | PSD + STAI-T | Yes | 0.505 | 0.004 | 0.509 | 0.003 | 0.572 | 0.006 | 0.514 | 0.004 | 0.541 | 0.004 | 0.037 | 0.002 | 0.132 | 0.046 |
| | | No | 0.898 | 0.001 | 0.968 | 0.000 | 0.891 | 0.001 | 0.907 | 0.001 | 0.899 | 0.001 | 0.204 | 0.003 | 0.756 | 0.002 |
| | PSD + STAI-S + STAI-T | Yes | 0.499 | 0.005 | 0.494 | 0.004 | 0.588 | 0.006 | 0.508 | 0.004 | 0.545 | 0.005 | 0.050 | 0.003 | -0.163 | 0.062 |
| | | No | 0.920 | 0.001 | 0.982 | 0.000 | 0.910 | 0.001 | 0.931 | 0.001 | 0.921 | 0.001 | 0.082 | 0.004 | 0.746 | 0.003 |
| Beta | PSD | Yes | 0.500 | 0.005 | 0.496 | 0.004 | 0.572 | 0.006 | 0.509 | 0.004 | 0.539 | 0.005 | 0.048 | 0.003 | -0.112 | 0.059 |
| | | No | 0.862 | 0.001 | 0.931 | 0.000 | 0.860 | 0.002 | 0.869 | 0.002 | 0.864 | 0.001 | 0.041 | 0.003 | 1.015 | 0.004 |
| | PSD + STAI-S | Yes | 0.499 | 0.004 | 0.496 | 0.004 | 0.571 | 0.006 | 0.509 | 0.004 | 0.538 | 0.005 | 0.049 | 0.003 | -0.124 | 0.059 |
| | | No | 0.928 | 0.001 | 0.981 | 0.000 | 0.934 | 0.001 | 0.926 | 0.001 | 0.930 | 0.001 | -0.070 | 0.004 | 0.671 | 0.002 |

| | | | | | | | | | | | | | | | | |
|---|---|---|---|---|---|---|---|---|---|---|---|---|---|---|---|---|
| | PSD + STAI-T | Yes | 0.519 | 0.004 | 0.522 | 0.003 | 0.577 | 0.005 | 0.527 | 0.004 | 0.551 | 0.004 | 0.033 | 0.002 | 0.227 | 0.041 |
| | | No | 0.948 | 0.001 | 0.994 | 0.000 | 0.927 | 0.001 | 0.970 | 0.001 | 0.948 | 0.001 | 0.887 | 0.007 | 0.806 | 0.004 |
| | PSD + STAI-S + STAI-T | Yes | 0.476 | 0.005 | 0.466 | 0.005 | 0.559 | 0.007 | 0.489 | 0.004 | 0.522 | 0.005 | 0.071 | 0.004 | -0.614 | 0.078 |
| | | No | 0.971 | 0.001 | 0.996 | 0.000 | 0.958 | 0.001 | 0.985 | 0.001 | 0.971 | 0.001 | 0.819 | 0.010 | 0.770 | 0.004 |
| Beta-1 | PSD | Yes | 0.490 | 0.005 | 0.479 | 0.004 | 0.590 | 0.007 | 0.500 | 0.004 | 0.541 | 0.005 | 0.062 | 0.003 | -0.425 | 0.072 |
| | | No | 0.789 | 0.002 | 0.873 | 0.001 | 0.800 | 0.002 | 0.790 | 0.002 | 0.795 | 0.002 | 0.011 | 0.002 | 1.029 | 0.006 |
| | PSD + STAI-S | Yes | 0.492 | 0.005 | 0.482 | 0.004 | 0.584 | 0.006 | 0.502 | 0.004 | 0.540 | 0.005 | 0.059 | 0.003 | -0.359 | 0.066 |
| | | No | 0.905 | 0.001 | 0.969 | 0.000 | 0.919 | 0.001 | 0.896 | 0.001 | 0.908 | 0.001 | -0.149 | 0.003 | 0.719 | 0.002 |
| | PSD + STAI-T | Yes | 0.502 | 0.004 | 0.499 | 0.004 | 0.568 | 0.006 | 0.511 | 0.004 | 0.538 | 0.004 | 0.044 | 0.002 | -0.018 | 0.055 |
| | | No | 0.933 | 0.001 | 0.982 | 0.000 | 0.904 | 0.001 | 0.962 | 0.001 | 0.932 | 0.001 | 0.579 | 0.005 | 0.834 | 0.003 |
| | PSD + STAI-S + STAI-T | Yes | 0.488 | 0.004 | 0.488 | 0.004 | 0.555 | 0.006 | 0.499 | 0.004 | 0.525 | 0.005 | 0.050 | 0.003 | -0.148 | 0.060 |
| | | No | 0.948 | 0.001 | 0.990 | 0.000 | 0.932 | 0.001 | 0.965 | 0.001 | 0.948 | 0.001 | 0.464 | 0.006 | 0.823 | 0.003 |
| Beta-2 | PSD | Yes | 0.516 | 0.004 | 0.520 | 0.003 | 0.574 | 0.006 | 0.524 | 0.003 | 0.548 | 0.004 | 0.032 | 0.002 | 0.256 | 0.040 |
| | | No | 0.783 | 0.002 | 0.862 | 0.001 | 0.791 | 0.002 | 0.785 | 0.002 | 0.788 | 0.002 | 0.020 | 0.002 | 1.008 | 0.005 |
| | PSD + STAI-S | Yes | 0.515 | 0.004 | 0.522 | 0.003 | 0.572 | 0.006 | 0.523 | 0.003 | 0.547 | 0.004 | 0.031 | 0.002 | 0.286 | 0.041 |
| | | No | 0.893 | 0.001 | 0.959 | 0.000 | 0.920 | 0.001 | 0.877 | 0.001 | 0.898 | 0.001 | -0.191 | 0.003 | 0.697 | 0.002 |
| | PSD + STAI-T | Yes | 0.507 | 0.004 | 0.502 | 0.004 | 0.597 | 0.006 | 0.515 | 0.004 | 0.553 | 0.004 | 0.042 | 0.002 | 0.038 | 0.053 |
| | | No | 0.925 | 0.001 | 0.982 | 0.000 | 0.904 | 0.001 | 0.946 | 0.001 | 0.925 | 0.001 | 0.478 | 0.004 | 0.768 | 0.002 |
| | PSD + STAI-S + STAI-T | Yes | 0.500 | 0.004 | 0.501 | 0.004 | 0.556 | 0.006 | 0.509 | 0.004 | 0.531 | 0.004 | 0.040 | 0.002 | 0.063 | 0.050 |
| | | No | 0.940 | 0.001 | 0.989 | 0.000 | 0.923 | 0.001 | 0.958 | 0.001 | 0.940 | 0.001 | 0.392 | 0.005 | 0.799 | 0.003 |
| Beta-3 | PSD | Yes | 0.499 | 0.004 | 0.493 | 0.004 | 0.596 | 0.006 | 0.509 | 0.004 | 0.549 | 0.004 | 0.050 | 0.003 | -0.144 | 0.059 |
| | | No | 0.861 | 0.001 | 0.930 | 0.000 | 0.853 | 0.002 | 0.873 | 0.002 | 0.863 | 0.001 | 0.098 | 0.003 | 0.951 | 0.003 |
| | PSD + STAI-S | Yes | 0.498 | 0.004 | 0.492 | 0.004 | 0.594 | 0.006 | 0.507 | 0.004 | 0.547 | 0.004 | 0.050 | 0.003 | -0.159 | 0.060 |
| | | No | 0.929 | 0.001 | 0.984 | 0.000 | 0.953 | 0.001 | 0.912 | 0.001 | 0.932 | 0.001 | -0.309 | 0.005 | 0.714 | 0.002 |
| | PSD + STAI-T | Yes | 0.503 | 0.004 | 0.506 | 0.003 | 0.583 | 0.006 | 0.512 | 0.003 | 0.545 | 0.004 | 0.037 | 0.002 | 0.132 | 0.045 |
| | | No | 0.942 | 0.001 | 0.992 | 0.000 | 0.923 | 0.001 | 0.962 | 0.001 | 0.942 | 0.001 | 0.700 | 0.006 | 0.822 | 0.004 |

| | | | | | | | | | | | | | | | | |
|---|---|---|---|---|---|---|---|---|---|---|---|---|---|---|---|---|
| | PSD + STAI-S + STAI-T | Yes | 0.506 | 0.004 | 0.504 | 0.004 | 0.586 | 0.006 | 0.515 | 0.004 | 0.548 | 0.004 | 0.041 | 0.002 | 0.062 | 0.051 |
| | | No | 0.971 | 0.001 | 0.997 | 0.000 | 0.967 | 0.001 | 0.975 | 0.001 | 0.971 | 0.001 | 0.312 | 0.009 | 0.889 | 0.005 |
| Gamma | PSD | Yes | 0.509 | 0.004 | 0.506 | 0.004 | 0.599 | 0.006 | 0.517 | 0.004 | 0.555 | 0.004 | 0.040 | 0.002 | 0.077 | 0.051 |
| | | No | 0.888 | 0.001 | 0.956 | 0.000 | 0.867 | 0.002 | 0.910 | 0.001 | 0.888 | 0.001 | 0.233 | 0.004 | 0.976 | 0.004 |
| | PSD + STAI-S | Yes | 0.508 | 0.004 | 0.505 | 0.004 | 0.596 | 0.006 | 0.516 | 0.004 | 0.553 | 0.004 | 0.041 | 0.002 | 0.061 | 0.050 |
| | | No | 0.962 | 0.001 | 0.994 | 0.000 | 0.978 | 0.001 | 0.948 | 0.001 | 0.963 | 0.001 | -0.468 | 0.008 | 0.828 | 0.004 |
| | PSD + STAI-T | Yes | 0.496 | 0.005 | 0.492 | 0.004 | 0.577 | 0.006 | 0.506 | 0.004 | 0.539 | 0.005 | 0.048 | 0.003 | -0.108 | 0.060 |
| | | No | 0.958 | 0.001 | 0.994 | 0.000 | 0.939 | 0.001 | 0.977 | 0.001 | 0.958 | 0.001 | 0.743 | 0.010 | 0.818 | 0.004 |
| | PSD + STAI-S + STAI-T | Yes | 0.508 | 0.004 | 0.508 | 0.004 | 0.589 | 0.006 | 0.517 | 0.003 | 0.550 | 0.004 | 0.040 | 0.002 | 0.081 | 0.049 |
| | | No | 0.984 | 0.001 | 0.999 | 0.000 | 0.989 | 0.001 | 0.980 | 0.001 | 0.985 | 0.000 | -0.340 | 0.017 | 1.030 | 0.009 |

Abbreviations: PSD = Power spectral density; STAI-S = STAI-State; STAI-T = STAI-Trait; ACC = Accuracy; SD = Standard deviation; AUC = Area under the curve.
