## Supplementary Tables 6 for "Multivariate pattern analysis identifies potential inter-trial resting-state EEG biomarkers in fibromyalgia"

**Supplementary Table 6 LDA performance metrics for STFM and LTFM group comparison**

| Frequency | Permuted | ACC | ACC SD | AUC | AUC SD | Precision | Precision SD | Recall | Recall SD | F1-Score | F1-Score SD | Intercept | Intercept SD | Slope | Slope SD |
|---|---|---|---|---|---|---|---|---|---|---|---|---|---|---|---|
| Delta | Yes | 0.492 | 0.006 | 0.482 | 0.006 | 0.378 | 0.008 | 0.469 | 0.008 | 0.419 | 0.008 | -0.084 | 0.004 | -0.241 | 0.053 |
| | No | 0.813 | 0.003 | 0.890 | 0.001 | 0.804 | 0.004 | 0.809 | 0.003 | 0.806 | 0.003 | -0.019 | 0.005 | 0.979 | 0.009 |
| Theta | Yes | 0.512 | 0.006 | 0.513 | 0.005 | 0.430 | 0.008 | 0.495 | 0.007 | 0.460 | 0.007 | -0.055 | 0.003 | 0.164 | 0.043 |
| | No | 0.836 | 0.003 | 0.916 | 0.001 | 0.831 | 0.004 | 0.830 | 0.003 | 0.830 | 0.003 | -0.029 | 0.005 | 1.040 | 0.009 |
| Alpha | Yes | 0.515 | 0.006 | 0.515 | 0.005 | 0.429 | 0.008 | 0.498 | 0.007 | 0.461 | 0.007 | -0.058 | 0.003 | 0.127 | 0.047 |
| | No | 0.829 | 0.002 | 0.906 | 0.001 | 0.807 | 0.003 | 0.834 | 0.003 | 0.821 | 0.002 | 0.040 | 0.005 | 0.950 | 0.008 |
| Alpha-1 | Yes | 0.497 | 0.006 | 0.493 | 0.006 | 0.403 | 0.009 | 0.476 | 0.008 | 0.436 | 0.008 | -0.073 | 0.004 | -0.098 | 0.058 |
| | No | 0.798 | 0.003 | 0.874 | 0.001 | 0.780 | 0.004 | 0.797 | 0.003 | 0.789 | 0.003 | 0.000 | 0.004 | 0.970 | 0.008 |
| Alpha-2 | Yes | 0.515 | 0.006 | 0.515 | 0.005 | 0.429 | 0.008 | 0.498 | 0.007 | 0.461 | 0.007 | -0.058 | 0.003 | 0.120 | 0.046 |
| | No | 0.791 | 0.002 | 0.878 | 0.001 | 0.761 | 0.004 | 0.797 | 0.003 | 0.779 | 0.003 | 0.049 | 0.005 | 0.961 | 0.008 |
| Beta | Yes | 0.480 | 0.006 | 0.470 | 0.006 | 0.379 | 0.008 | 0.455 | 0.008 | 0.413 | 0.008 | -0.085 | 0.005 | -0.275 | 0.066 |
| | No | 0.934 | 0.001 | 0.983 | 0.000 | 0.953 | 0.002 | 0.914 | 0.002 | 0.933 | 0.001 | -0.363 | 0.010 | 0.847 | 0.008 |
| Beta-1 | Yes | 0.480 | 0.007 | 0.461 | 0.006 | 0.381 | 0.009 | 0.455 | 0.008 | 0.415 | 0.008 | -0.099 | 0.005 | -0.478 | 0.075 |
| | No | 0.866 | 0.002 | 0.940 | 0.001 | 0.866 | 0.003 | 0.859 | 0.003 | 0.862 | 0.002 | -0.068 | 0.005 | 0.870 | 0.006 |
| Beta-2 | Yes | 0.505 | 0.006 | 0.502 | 0.005 | 0.422 | 0.008 | 0.486 | 0.007 | 0.452 | 0.008 | -0.067 | 0.004 | -0.008 | 0.053 |
| | No | 0.868 | 0.002 | 0.943 | 0.001 | 0.892 | 0.003 | 0.844 | 0.003 | 0.867 | 0.002 | -0.194 | 0.006 | 0.951 | 0.006 |
| Beta-3 | Yes | 0.492 | 0.006 | 0.486 | 0.006 | 0.397 | 0.009 | 0.470 | 0.007 | 0.430 | 0.008 | -0.077 | 0.004 | -0.164 | 0.058 |
| | No | 0.935 | 0.001 | 0.981 | 0.000 | 0.947 | 0.002 | 0.922 | 0.002 | 0.934 | 0.001 | -0.264 | 0.008 | 0.827 | 0.006 |
| Gamma | Yes | 0.492 | 0.006 | 0.487 | 0.006 | 0.384 | 0.008 | 0.469 | 0.008 | 0.422 | 0.008 | -0.078 | 0.004 | -0.169 | 0.063 |
| | No | 0.958 | 0.001 | 0.994 | 0.000 | 0.967 | 0.001 | 0.948 | 0.002 | 0.957 | 0.001 | -0.453 | 0.014 | 0.912 | 0.008 |

Abbreviations: ACC = Accuracy; SD = Standard deviation; AUC = Area under the curve.
