## Supplementary Table 7 for "Multivariate pattern analysis identifies potential inter-trial resting-state EEG biomarkers in fibromyalgia"

**Supplementary Table 7 Haufe-corrected weights for LDA comparison between FM and HC groups**

| Channel | Delta | Theta | Alpha | Alpha-1 | Alpha-2 | Beta | Beta-1 | Beta-2 | Beta-3 | Gamma |
|---|---|---|---|---|---|---|---|---|---|---|
| Fp1 | 0.0130 | 0.0089 | 0.0061 | 0.0066 | 0.0057 | 0.0011 | 0.0049 | -0.0013 | -0.0011 | 0.0053 |
| Fpz | 0.0137 | 0.0103 | 0.0113 | 0.0102 | 0.0115 | 0.0207 | 0.0140 | 0.0144 | 0.0296 | 0.0452 |
| Fp2 | 0.0155 | 0.0122 | 0.0125 | 0.0120 | 0.0120 | 0.0094 | 0.0094 | 0.0095 | 0.0124 | 0.0263 |
| AF7 | 0.0031 | -0.0005 | -0.0008 | -0.0004 | -0.0008 | -0.0158 | -0.0056 | -0.0124 | -0.0276 | -0.0168 |
| AF3 | 0.0000 | -0.0017 | -0.0028 | -0.0027 | -0.0020 | -0.0199 | -0.0098 | -0.0198 | -0.0246 | -0.0406 |
| AF4 | 0.0091 | 0.0045 | 0.0005 | 0.0008 | 0.0002 | -0.0137 | -0.0061 | -0.0083 | -0.0180 | -0.0264 |
| AF8 | 0.0130 | 0.0117 | 0.0096 | 0.0098 | 0.0087 | 0.0075 | 0.0058 | 0.0032 | 0.0102 | 0.0076 |
| F7 | -0.0060 | -0.0030 | -0.0026 | -0.0023 | -0.0029 | -0.0174 | -0.0077 | -0.0133 | -0.0232 | -0.0253 |
| F5 | -0.0004 | 0.0003 | -0.0006 | 0.0000 | -0.0014 | -0.0220 | -0.0106 | -0.0183 | -0.0289 | -0.0317 |
| F3 | -0.0017 | -0.0025 | -0.0036 | -0.0021 | -0.0043 | -0.0157 | -0.0091 | -0.0122 | -0.0215 | -0.0277 |
| F1 | -0.0030 | -0.0020 | -0.0025 | -0.0018 | -0.0027 | -0.0039 | -0.0033 | -0.0024 | -0.0046 | -0.0063 |
| Fz | -0.0026 | -0.0015 | -0.0024 | -0.0015 | -0.0027 | -0.0029 | -0.0029 | -0.0017 | -0.0026 | -0.0032 |
| F2 | -0.0010 | -0.0010 | -0.0013 | -0.0004 | -0.0015 | -0.0018 | -0.0015 | 0.0000 | -0.0022 | -0.0014 |
| F4 | 0.0007 | -0.0013 | -0.0044 | -0.0027 | -0.0051 | -0.0153 | -0.0110 | -0.0061 | -0.0163 | -0.0251 |
| F6 | 0.0025 | 0.0013 | -0.0006 | 0.0001 | -0.0010 | -0.0102 | -0.0035 | -0.0069 | -0.0149 | -0.0207 |
| F8 | 0.0005 | 0.0003 | 0.0006 | 0.0011 | 0.0003 | -0.0010 | 0.0009 | -0.0002 | -0.0028 | -0.0026 |
| FC5 | -0.0011 | 0.0004 | -0.0009 | -0.0005 | -0.0017 | -0.0112 | -0.0076 | -0.0087 | -0.0147 | -0.0272 |
| FC3 | -0.0021 | -0.0010 | -0.0025 | -0.0013 | -0.0033 | -0.0042 | -0.0041 | -0.0034 | -0.0045 | -0.0105 |
| FC1 | -0.0031 | -0.0018 | -0.0030 | -0.0018 | -0.0037 | -0.0036 | -0.0040 | -0.0024 | -0.0031 | -0.0056 |
| FCz | -0.0030 | -0.0010 | -0.0028 | -0.0015 | -0.0035 | -0.0046 | -0.0048 | -0.0031 | -0.0036 | -0.0046 |
| FC2 | -0.0036 | -0.0025 | -0.0028 | -0.0016 | -0.0034 | -0.0034 | -0.0041 | -0.0021 | -0.0024 | -0.0029 |
| FC4 | -0.0011 | -0.0008 | -0.0019 | -0.0006 | -0.0028 | -0.0037 | -0.0033 | -0.0025 | -0.0040 | -0.0046 |
| FC6 | -0.0004 | -0.0003 | -0.0006 | -0.0004 | -0.0008 | 0.0007 | -0.0003 | -0.0004 | 0.0010 | -0.0016 |
| T7 | -0.0015 | 0.0003 | 0.0048 | 0.0051 | 0.0032 | -0.0155 | -0.0095 | -0.0185 | -0.0174 | -0.0210 |
| C5 | 0.0003 | 0.0002 | 0.0005 | 0.0011 | -0.0004 | -0.0009 | -0.0032 | -0.0006 | 0.0011 | -0.0075 |
| C3 | -0.0017 | -0.0023 | -0.0024 | -0.0010 | -0.0032 | -0.0017 | -0.0040 | 0.0005 | 0.0004 | -0.0039 |
| C1 | -0.0032 | -0.0029 | -0.0033 | -0.0019 | -0.0039 | -0.0023 | -0.0043 | -0.0010 | -0.0002 | -0.0023 |
| Cz | -0.0031 | -0.0015 | -0.0015 | -0.0009 | -0.0020 | -0.0016 | -0.0040 | -0.0009 | 0.0012 | -0.0025 |
| C2 | -0.0022 | -0.0018 | -0.0033 | -0.0027 | -0.0038 | -0.0021 | -0.0036 | -0.0018 | -0.0004 | -0.0008 |
| C4 | -0.0009 | -0.0025 | -0.0061 | -0.0067 | -0.0060 | 0.0000 | -0.0034 | -0.0005 | 0.0037 | 0.0058 |
| C6 | -0.0005 | -0.0006 | -0.0003 | -0.0002 | -0.0009 | 0.0142 | 0.0054 | 0.0113 | 0.0209 | 0.0341 |
| T8 | -0.0018 | -0.0004 | -0.0018 | -0.0001 | -0.0024 | 0.0089 | 0.0027 | 0.0088 | 0.0147 | 0.0240 |
| TP7 | 0.0001 | 0.0013 | 0.0030 | 0.0039 | 0.0015 | -0.0024 | 0.0021 | -0.0011 | -0.0080 | -0.0042 |
| CP5 | -0.0007 | -0.0001 | 0.0001 | 0.0001 | 0.0000 | 0.0010 | -0.0005 | 0.0024 | 0.0017 | -0.0044 |
| CP3 | -0.0004 | -0.0010 | -0.0006 | -0.0005 | -0.0003 | 0.0002 | -0.0016 | 0.0023 | 0.0013 | -0.0007 |
| CP1 | 0.0008 | -0.0008 | -0.0005 | -0.0001 | -0.0004 | -0.0006 | -0.0021 | 0.0003 | 0.0006 | 0.0002 |
| CPz | -0.0020 | -0.0016 | 0.0008 | 0.0006 | 0.0008 | -0.0001 | -0.0015 | -0.0001 | 0.0015 | 0.0003 |
| CP2 | -0.0012 | -0.0011 | 0.0002 | -0.0008 | 0.0005 | 0.0006 | -0.0006 | 0.0004 | 0.0018 | 0.0011 |
| CP4 | -0.0013 | -0.0012 | -0.0012 | -0.0028 | -0.0006 | 0.0019 | 0.0003 | 0.0012 | 0.0034 | 0.0039 |
| CP6 | -0.0004 | -0.0005 | 0.0004 | -0.0015 | 0.0019 | 0.0125 | 0.0091 | 0.0082 | 0.0148 | 0.0276 |
| TP8 | -0.0023 | 0.0006 | 0.0061 | 0.0022 | 0.0090 | 0.0294 | 0.0205 | 0.0214 | 0.0361 | 0.0670 |
| P7 | -0.0008 | -0.0006 | -0.0012 | -0.0011 | -0.0007 | 0.0030 | 0.0025 | 0.0026 | 0.0020 | 0.0045 |
| P5 | -0.0001 | -0.0004 | -0.0009 | -0.0007 | -0.0005 | 0.0038 | 0.0016 | 0.0037 | 0.0044 | 0.0036 |
| P3 | -0.0009 | -0.0008 | -0.0023 | -0.0026 | -0.0017 | 0.0018 | 0.0006 | 0.0023 | 0.0020 | 0.0012 |
| P1 | -0.0019 | -0.0019 | -0.0026 | -0.0037 | -0.0021 | 0.0003 | -0.0006 | 0.0005 | 0.0008 | 0.0004 |
| Pz | -0.0019 | -0.0017 | -0.0013 | -0.0027 | -0.0009 | 0.0006 | -0.0002 | 0.0003 | 0.0012 | 0.0006 |
| P2 | -0.0025 | -0.0017 | -0.0026 | -0.0042 | -0.0022 | 0.0008 | -0.0002 | 0.0003 | 0.0016 | 0.0011 |
| P4 | -0.0015 | -0.0010 | 0.0012 | -0.0005 | 0.0023 | 0.0044 | 0.0034 | 0.0028 | 0.0038 | 0.0026 |
| P6 | -0.0025 | -0.0046 | -0.0022 | -0.0041 | -0.0001 | 0.0069 | 0.0050 | 0.0041 | 0.0077 | 0.0118 |
| P8 | -0.0024 | -0.0007 | -0.0002 | -0.0006 | 0.0013 | 0.0107 | 0.0064 | 0.0074 | 0.0137 | 0.0189 |
| PO7 | 0.0014 | 0.0013 | 0.0024 | 0.0029 | 0.0026 | 0.0063 | 0.0059 | 0.0040 | 0.0040 | 0.0030 |
| PO5 | -0.0018 | -0.0020 | -0.0010 | -0.0011 | -0.0005 | 0.0043 | 0.0039 | 0.0030 | 0.0023 | 0.0017 |
| PO3 | -0.0006 | -0.0006 | -0.0007 | -0.0020 | -0.0003 | 0.0027 | 0.0031 | 0.0020 | 0.0013 | -0.0003 |
| POz | -0.0021 | -0.0012 | -0.0013 | -0.0023 | -0.0012 | 0.0015 | 0.0015 | 0.0015 | 0.0012 | -0.0005 |
| PO4 | -0.0026 | -0.0013 | -0.0001 | -0.0019 | 0.0005 | 0.0016 | 0.0012 | 0.0017 | 0.0013 | -0.0004 |

| | | | | | | | | | | |
|---|---|---|---|---|---|---|---|---|---|---|
| PO6 | -0.0019 | -0.0012 | 0.0011 | 0.0003 | 0.0022 | 0.0066 | 0.0038 | 0.0052 | 0.0074 | 0.0074 |
| PO8 | -0.0004 | 0.0005 | 0.0033 | 0.0025 | 0.0046 | 0.0229 | 0.0112 | 0.0161 | 0.0300 | 0.0395 |
| O1 | 0.0000 | 0.0011 | 0.0042 | 0.0039 | 0.0046 | 0.0046 | 0.0065 | 0.0024 | 0.0011 | -0.0038 |
| Oz | 0.0029 | 0.0029 | 0.0029 | 0.0030 | 0.0026 | 0.0035 | 0.0039 | 0.0028 | 0.0020 | -0.0015 |
| O2 | -0.0004 | 0.0009 | 0.0020 | 0.0024 | 0.0018 | 0.0033 | 0.0028 | 0.0032 | 0.0019 | -0.0062 |
