## Supplementary Table 8 for "Multivariate pattern analysis identifies potential inter-trial resting-state EEG biomarkers in fibromyalgia"

**Supplementary Table 8 Haufe-corrected weights for LDA comparison between STFM and LTFM groups**

| | Delta | Theta | Alpha | Alpha-1 | Alpha-2 | Beta | Beta-1 | Beta-2 | Beta-3 | Gamma |
|---|---|---|---|---|---|---|---|---|---|---|
| Fp1 | 0.0072 | 0.0074 | 0.0136 | 0.0094 | 0.0157 | 0.0092 | 0.0089 | 0.0073 | -0.0006 | 0.0366 |
| Fpz | 0.0083 | 0.0120 | 0.0058 | 0.0053 | 0.0062 | 0.0123 | 0.0083 | 0.0109 | 0.0091 | 0.0065 |
| Fp2 | 0.0046 | 0.0088 | 0.0041 | 0.0028 | 0.0057 | 0.0415 | 0.0200 | 0.0315 | 0.0378 | 0.0411 |
| AF7 | 0.0045 | 0.0028 | 0.0063 | 0.0030 | 0.0077 | 0.0254 | 0.0144 | 0.0128 | 0.0228 | 0.0270 |
| AF3 | 0.0017 | 0.0012 | 0.0019 | 0.0018 | 0.0015 | -0.0166 | 0.0147 | 0.0119 | -0.0561 | 0.0144 |
| AF4 | 0.0020 | 0.0061 | 0.0057 | 0.0041 | 0.0065 | 0.0144 | 0.0150 | 0.0005 | 0.0113 | 0.0590 |
| AF8 | 0.0045 | 0.0050 | 0.0036 | 0.0020 | 0.0048 | 0.0350 | 0.0216 | 0.0226 | 0.0485 | 0.0974 |
| F7 | 0.0089 | 0.0026 | 0.0014 | 0.0019 | -0.0019 | -0.0558 | -0.0160 | -0.0303 | -0.0680 | -0.0747 |
| F5 | 0.0040 | 0.0006 | -0.0015 | -0.0007 | -0.0035 | -0.0452 | -0.0096 | -0.0169 | -0.0574 | -0.0524 |
| F3 | 0.0032 | 0.0050 | 0.0015 | 0.0032 | -0.0012 | -0.0219 | -0.0042 | -0.0095 | -0.0322 | -0.0375 |
| F1 | 0.0052 | 0.0070 | 0.0060 | 0.0063 | 0.0040 | 0.0065 | 0.0035 | 0.0044 | 0.0070 | 0.0046 |
| Fz | 0.0040 | 0.0070 | 0.0059 | 0.0060 | 0.0042 | 0.0119 | 0.0055 | 0.0079 | 0.0133 | 0.0126 |
| F2 | 0.0033 | 0.0064 | 0.0054 | 0.0052 | 0.0039 | 0.0120 | 0.0053 | 0.0082 | 0.0136 | 0.0141 |
| F4 | 0.0023 | 0.0059 | 0.0092 | 0.0056 | 0.0094 | 0.0211 | 0.0206 | 0.0094 | 0.0220 | 0.0690 |
| F6 | 0.0018 | 0.0013 | 0.0029 | 0.0011 | 0.0027 | 0.0126 | 0.0089 | 0.0064 | 0.0229 | 0.0308 |
| F8 | -0.0020 | -0.0025 | 0.0005 | -0.0001 | -0.0004 | 0.0065 | 0.0041 | 0.0001 | 0.0120 | 0.0160 |
| FC5 | 0.0097 | 0.0028 | -0.0005 | -0.0010 | -0.0017 | -0.0359 | -0.0128 | -0.0289 | -0.0451 | -0.0826 |
| FC3 | -0.0032 | 0.0011 | 0.0015 | 0.0011 | 0.0007 | 0.0037 | 0.0034 | 0.0014 | 0.0040 | 0.0006 |
| FC1 | 0.0007 | 0.0023 | 0.0020 | 0.0022 | 0.0007 | 0.0077 | 0.0038 | 0.0050 | 0.0094 | 0.0068 |
| FCz | 0.0013 | 0.0052 | 0.0048 | 0.0049 | 0.0036 | 0.0186 | 0.0091 | 0.0140 | 0.0202 | 0.0183 |
| FC2 | 0.0045 | 0.0067 | 0.0047 | 0.0045 | 0.0034 | 0.0149 | 0.0076 | 0.0099 | 0.0162 | 0.0138 |
| FC4 | -0.0003 | 0.0004 | 0.0000 | 0.0000 | -0.0008 | 0.0061 | 0.0026 | 0.0053 | 0.0079 | 0.0054 |
| FC6 | -0.0023 | -0.0032 | -0.0055 | -0.0058 | -0.0057 | -0.0166 | -0.0070 | -0.0050 | -0.0184 | -0.0228 |
| T7 | 0.0036 | -0.0025 | -0.0017 | -0.0029 | -0.0019 | -0.0541 | -0.0365 | -0.0526 | -0.0432 | -0.1191 |
| C5 | 0.0012 | -0.0007 | 0.0038 | 0.0011 | 0.0040 | -0.0123 | -0.0046 | -0.0072 | -0.0208 | -0.0527 |
| C3 | 0.0029 | 0.0011 | 0.0062 | 0.0023 | 0.0073 | 0.0054 | 0.0061 | 0.0050 | -0.0012 | -0.0117 |
| C1 | -0.0046 | -0.0040 | 0.0007 | -0.0014 | 0.0023 | 0.0064 | 0.0039 | 0.0039 | 0.0070 | 0.0069 |
| Cz | -0.0012 | 0.0010 | 0.0024 | 0.0019 | 0.0027 | 0.0132 | 0.0063 | 0.0112 | 0.0138 | 0.0170 |
| C2 | -0.0006 | 0.0003 | 0.0041 | 0.0027 | 0.0050 | 0.0121 | 0.0081 | 0.0082 | 0.0104 | 0.0100 |
| C4 | -0.0008 | -0.0032 | 0.0009 | -0.0014 | 0.0025 | 0.0090 | 0.0074 | 0.0057 | 0.0050 | 0.0001 |
| C6 | 0.0006 | -0.0018 | 0.0001 | -0.0012 | -0.0001 | -0.0022 | -0.0006 | -0.0027 | -0.0067 | -0.0211 |
| T8 | 0.0036 | 0.0003 | -0.0012 | 0.0020 | -0.0054 | -0.0384 | -0.0184 | -0.0180 | -0.0432 | -0.0519 |
| TP7 | -0.0011 | -0.0028 | -0.0047 | -0.0038 | -0.0058 | -0.0409 | -0.0208 | -0.0311 | -0.0519 | -0.1045 |
| CP5 | 0.0048 | 0.0031 | 0.0048 | 0.0035 | 0.0034 | 0.0015 | 0.0026 | 0.0041 | -0.0057 | -0.0306 |
| CP3 | -0.0030 | -0.0024 | 0.0020 | 0.0007 | 0.0019 | 0.0074 | 0.0037 | 0.0080 | 0.0042 | -0.0020 |
| CP1 | -0.0028 | -0.0040 | -0.0018 | -0.0020 | -0.0006 | 0.0030 | -0.0012 | 0.0037 | 0.0054 | 0.0040 |
| CPz | -0.0029 | -0.0036 | -0.0017 | -0.0013 | -0.0003 | 0.0037 | -0.0010 | 0.0035 | 0.0070 | 0.0063 |
| CP2 | -0.0029 | -0.0030 | 0.0000 | 0.0003 | 0.0012 | 0.0046 | 0.0013 | 0.0025 | 0.0054 | 0.0046 |
| CP4 | -0.0031 | -0.0035 | 0.0007 | 0.0006 | 0.0011 | 0.0063 | 0.0029 | 0.0032 | 0.0057 | 0.0029 |
| CP6 | -0.0001 | -0.0013 | 0.0021 | 0.0010 | 0.0023 | 0.0077 | 0.0032 | 0.0044 | 0.0087 | 0.0031 |
| TP8 | 0.0057 | 0.0009 | 0.0020 | 0.0010 | 0.0014 | -0.0065 | -0.0043 | -0.0054 | -0.0055 | -0.0026 |
| P7 | 0.0006 | -0.0028 | -0.0053 | -0.0062 | -0.0039 | 0.0074 | 0.0025 | -0.0002 | 0.0089 | 0.0109 |
| P5 | 0.0008 | 0.0002 | -0.0006 | 0.0001 | -0.0008 | 0.0071 | 0.0019 | 0.0039 | 0.0081 | 0.0033 |
| P3 | -0.0010 | -0.0003 | 0.0026 | 0.0037 | 0.0018 | 0.0085 | 0.0028 | 0.0064 | 0.0078 | 0.0049 |
| P1 | -0.0033 | -0.0023 | 0.0001 | 0.0024 | 0.0002 | 0.0025 | -0.0025 | 0.0027 | 0.0058 | 0.0052 |
| Pz | -0.0054 | -0.0037 | -0.0029 | -0.0001 | -0.0024 | 0.0016 | -0.0032 | 0.0015 | 0.0055 | 0.0057 |
| P2 | -0.0052 | -0.0036 | -0.0008 | 0.0022 | -0.0008 | 0.0022 | -0.0031 | 0.0015 | 0.0058 | 0.0064 |
| P4 | -0.0025 | 0.0000 | -0.0016 | 0.0000 | -0.0022 | 0.0055 | -0.0004 | 0.0023 | 0.0094 | 0.0111 |
| P6 | -0.0183 | -0.0184 | -0.0100 | -0.0090 | -0.0081 | -0.0094 | -0.0072 | -0.0078 | -0.0098 | -0.0131 |
| P8 | 0.0061 | 0.0004 | -0.0036 | -0.0021 | -0.0032 | -0.0003 | -0.0063 | -0.0017 | 0.0086 | 0.0136 |
| PO7 | -0.0028 | -0.0041 | -0.0116 | -0.0105 | -0.0098 | -0.0057 | -0.0081 | -0.0066 | 0.0007 | -0.0090 |
| PO5 | -0.0014 | -0.0010 | -0.0090 | -0.0062 | -0.0084 | -0.0043 | -0.0078 | -0.0047 | 0.0017 | 0.0000 |
| PO3 | -0.0067 | -0.0041 | -0.0056 | -0.0027 | -0.0056 | -0.0027 | -0.0080 | -0.0024 | 0.0047 | 0.0087 |
| POz | -0.0087 | -0.0062 | -0.0075 | -0.0050 | -0.0071 | -0.0026 | -0.0080 | -0.0024 | 0.0050 | 0.0094 |
| PO4 | -0.0081 | -0.0050 | -0.0084 | -0.0054 | -0.0082 | 0.0001 | -0.0071 | -0.0015 | 0.0081 | 0.0119 |

| | | | | | | | | | |
|---|---|---|---|---|---|---|---|---|---|
| PO6 | -0.0064 | -0.0055 | -0.0077 | -0.0057 | -0.0063 | -0.0027 | -0.0092 | -0.0043 | 0.0043 | 0.0029 |
| PO8 | -0.0066 | -0.0065 | -0.0079 | -0.0059 | -0.0065 | -0.0025 | -0.0087 | -0.0040 | 0.0045 | 0.0056 |
| O1 | 0.0004 | -0.0001 | -0.0085 | -0.0077 | -0.0069 | -0.0054 | -0.0063 | -0.0081 | 0.0002 | 0.0039 |
| Oz | -0.0019 | -0.0013 | -0.0046 | -0.0037 | -0.0044 | 0.0009 | -0.0040 | -0.0017 | 0.0077 | 0.0172 |
| O2 | -0.0027 | -0.0014 | -0.0048 | -0.0040 | -0.0039 | 0.0065 | -0.0035 | 0.0016 | 0.0183 | 0.0387 |
