## Supplementary Table 9 for "Multivariate pattern analysis identifies potential inter-trial resting-state EEG biomarkers in fibromyalgia"

**Supplementary Table 9 Medication effects between STFM and LTFM groups across frequency bands**

| | Delta | | Theta | | Alpha | | Alpha-1 | | Alpha-2 | | Beta | | Beta-1 | | Beta-2 | | Beta-3 | | Gamma | |
|---|---|---|---|---|---|---|---|---|---|---|---|---|---|---|---|---|---|---|---|---|
| **Channel** | **AD** | **AG** | **AD** | **AG** | **AD** | **AG** | **AD** | **AG** | **AD** | **AG** | **AD** | **AG** | **AD** | **AG** | **AD** | **AG** | **AD** | **AG** | **AD** | **AG** |
| Fp1 | 0.53 | 1.00 | 0.99 | 1.00 | 0.95 | 1.00 | 1.00 | 1.00 | 0.92 | 1.00 | 1.00 | 1.00 | 1.00 | 1.00 | 1.00 | 1.00 | 1.00 | 1.00 | 1.00 | 1.00 |
| Fpz | 0.69 | 1.00 | 1.00 | 1.00 | 0.93 | 1.00 | 1.00 | 1.00 | 0.86 | 1.00 | 1.00 | 1.00 | 1.00 | 1.00 | 1.00 | 1.00 | 1.00 | 1.00 | 1.00 | 1.00 |
| Fp2 | 0.27 | 1.00 | 0.96 | 1.00 | 0.87 | 1.00 | 0.94 | 1.00 | 0.75 | 1.00 | 1.00 | 1.00 | 1.00 | 1.00 | 1.00 | 1.00 | 1.00 | 1.00 | 1.00 | 1.00 |
| AF7 | 0.92 | 1.00 | 1.00 | 1.00 | 0.99 | 1.00 | 1.00 | 1.00 | 0.93 | 1.00 | 1.00 | 0.99 | 0.99 | 0.28 | 0.57 | **0.03** | 0.60 | 0.07 | 1.00 | 0.08 |
| AF3 | 0.84 | 1.00 | 1.00 | 1.00 | 1.00 | 1.00 | 1.00 | 1.00 | 0.99 | 1.00 | 1.00 | 1.00 | 0.99 | 1.00 | 1.00 | 1.00 | 1.00 | 1.00 | 1.00 | 1.00 |
| AF4 | 1.00 | 1.00 | 1.00 | 1.00 | 1.00 | 1.00 | 1.00 | 1.00 | 1.00 | 1.00 | 1.00 | 1.00 | 1.00 | 0.92 | 1.00 | 0.95 | 1.00 | 0.92 | 1.00 | 0.90 |
| AF8 | 1.00 | 1.00 | 1.00 | 1.00 | 0.99 | 1.00 | 1.00 | 1.00 | 0.99 | 1.00 | 1.00 | 1.00 | 0.98 | 1.00 | 1.00 | 0.99 | 1.00 | 1.00 | 1.00 | 1.00 |
| F7 | 1.00 | 0.54 | 1.00 | 1.00 | 1.00 | 0.96 | 1.00 | 1.00 | 1.00 | 0.98 | 1.00 | 1.00 | 1.00 | 0.71 | 0.58 | 0.10 | 0.87 | 0.18 | 1.00 | 0.62 |
| F5 | 1.00 | 0.96 | 1.00 | 1.00 | 1.00 | 1.00 | 1.00 | 1.00 | 1.00 | 1.00 | 1.00 | 0.96 | 1.00 | 0.40 | 0.55 | 0.07 | 0.78 | 0.10 | 1.00 | 0.40 |
| F3 | 1.00 | 1.00 | 1.00 | 1.00 | 1.00 | 1.00 | 1.00 | 1.00 | 1.00 | 1.00 | 1.00 | 1.00 | 1.00 | 0.98 | 1.00 | 1.00 | 1.00 | 0.98 | 1.00 | 0.86 |
| F1 | 1.00 | 1.00 | 1.00 | 1.00 | 1.00 | 1.00 | 1.00 | 1.00 | 1.00 | 1.00 | 1.00 | 1.00 | 1.00 | 1.00 | 1.00 | 1.00 | 1.00 | 1.00 | 1.00 | 1.00 |
| Fz | 1.00 | 1.00 | 1.00 | 1.00 | 1.00 | 1.00 | 1.00 | 1.00 | 1.00 | 1.00 | 1.00 | 1.00 | 1.00 | 1.00 | 1.00 | 1.00 | 1.00 | 1.00 | 1.00 | 1.00 |
| F2 | 1.00 | 1.00 | 1.00 | 1.00 | 1.00 | 1.00 | 1.00 | 1.00 | 1.00 | 1.00 | 1.00 | 1.00 | 1.00 | 1.00 | 1.00 | 1.00 | 1.00 | 1.00 | 1.00 | 1.00 |
| F4 | 1.00 | 1.00 | 1.00 | 1.00 | 1.00 | 1.00 | 1.00 | 1.00 | 1.00 | 1.00 | 1.00 | 0.99 | 1.00 | 0.79 | 0.92 | 0.54 | 0.88 | 0.51 | 0.98 | 0.47 |
| F6 | 1.00 | 1.00 | 1.00 | 1.00 | 1.00 | 1.00 | 1.00 | 1.00 | 1.00 | 0.99 | 1.00 | 1.00 | 1.00 | 1.00 | 1.00 | 1.00 | 1.00 | 1.00 | 1.00 | 1.00 |
| F8 | 1.00 | 1.00 | 1.00 | 1.00 | 1.00 | 0.99 | 1.00 | 1.00 | 1.00 | 0.98 | 1.00 | 1.00 | 1.00 | 1.00 | 1.00 | 1.00 | 1.00 | 1.00 | 1.00 | 1.00 |
| FC5 | 0.97 | 0.25 | 1.00 | 0.61 | 1.00 | 0.65 | 0.99 | 1.00 | 1.00 | 0.69 | 1.00 | 1.00 | 1.00 | 1.00 | 1.00 | 1.00 | 1.00 | 1.00 | 1.00 | 1.00 |
| FC3 | 1.00 | 1.00 | 1.00 | 1.00 | 1.00 | 1.00 | 1.00 | 1.00 | 1.00 | 1.00 | 1.00 | 1.00 | 1.00 | 1.00 | 1.00 | 1.00 | 1.00 | 1.00 | 1.00 | 1.00 |
| FC1 | 1.00 | 1.00 | 1.00 | 1.00 | 1.00 | 1.00 | 1.00 | 1.00 | 1.00 | 1.00 | 1.00 | 1.00 | 1.00 | 1.00 | 1.00 | 1.00 | 1.00 | 1.00 | 1.00 | 1.00 |
| FCz | 1.00 | 1.00 | 1.00 | 1.00 | 1.00 | 1.00 | 1.00 | 0.99 | 1.00 | 1.00 | 1.00 | 1.00 | 1.00 | 1.00 | 1.00 | 1.00 | 1.00 | 1.00 | 1.00 | 1.00 |
| FC2 | 1.00 | 1.00 | 1.00 | 1.00 | 1.00 | 1.00 | 1.00 | 1.00 | 1.00 | 1.00 | 1.00 | 1.00 | 1.00 | 1.00 | 1.00 | 1.00 | 1.00 | 1.00 | 1.00 | 1.00 |
| FC4 | 1.00 | 1.00 | 1.00 | 1.00 | 1.00 | 1.00 | 1.00 | 1.00 | 1.00 | 1.00 | 1.00 | 1.00 | 1.00 | 1.00 | 1.00 | 1.00 | 1.00 | 1.00 | 1.00 | 1.00 |
| FC6 | 1.00 | 1.00 | 1.00 | 1.00 | 1.00 | 1.00 | 1.00 | 1.00 | 1.00 | 1.00 | 1.00 | 1.00 | 1.00 | 1.00 | 1.00 | 1.00 | 1.00 | 1.00 | 1.00 | 1.00 |
| T7 | 1.00 | 0.99 | 1.00 | 1.00 | 1.00 | 0.98 | 1.00 | 1.00 | 1.00 | 0.95 | 0.66 | 0.81 | 0.85 | 0.92 | 1.00 | 1.00 | 1.00 | 1.00 | 1.00 | 0.81 |
| C5 | 1.00 | 1.00 | 1.00 | 1.00 | 1.00 | 1.00 | 1.00 | 1.00 | 1.00 | 1.00 | 1.00 | 1.00 | 1.00 | 1.00 | 1.00 | 1.00 | 1.00 | 1.00 | 1.00 | 1.00 |
| C3 | 1.00 | 1.00 | 1.00 | 1.00 | 1.00 | 1.00 | 1.00 | 1.00 | 1.00 | 0.98 | 1.00 | 1.00 | 1.00 | 1.00 | 1.00 | 1.00 | 1.00 | 1.00 | 1.00 | 1.00 |
| C1 | 1.00 | 1.00 | 1.00 | 1.00 | 1.00 | 1.00 | 1.00 | 1.00 | 1.00 | 0.99 | 1.00 | 1.00 | 1.00 | 1.00 | 1.00 | 1.00 | 1.00 | 1.00 | 1.00 | 1.00 |
| Cz | 1.00 | 1.00 | 1.00 | 1.00 | 1.00 | 1.00 | 1.00 | 1.00 | 1.00 | 1.00 | 1.00 | 1.00 | 1.00 | 1.00 | 1.00 | 1.00 | 1.00 | 1.00 | 1.00 | 1.00 |
| C2 | 1.00 | 1.00 | 1.00 | 1.00 | 1.00 | 1.00 | 1.00 | 1.00 | 1.00 | 0.98 | 1.00 | 1.00 | 1.00 | 1.00 | 1.00 | 1.00 | 1.00 | 1.00 | 1.00 | 1.00 |

| | | | | | | | | | | | | | | | | | | | | |
|---|---|---|---|---|---|---|---|---|---|---|---|---|---|---|---|---|---|---|---|---|
| C4 | 1.00 | 1.00 | 1.00 | 1.00 | 1.00 | 0.99 | 1.00 | 0.97 | 1.00 | 0.95 | 1.00 | 1.00 | 1.00 | 1.00 | 1.00 | 1.00 | 1.00 | 1.00 | 1.00 | 1.00 |
| C6 | 1.00 | 1.00 | 1.00 | 1.00 | 1.00 | 1.00 | 1.00 | 1.00 | 1.00 | 1.00 | 1.00 | 1.00 | 1.00 | 1.00 | 1.00 | 1.00 | 1.00 | 1.00 | 1.00 | 1.00 |
| T8 | 1.00 | 1.00 | 1.00 | 1.00 | 1.00 | 0.98 | 0.99 | 1.00 | 1.00 | 0.99 | 0.42 | 1.00 | 0.79 | 1.00 | 1.00 | 1.00 | 1.00 | 1.00 | 0.90 | 1.00 |
| TP7 | 1.00 | 1.00 | 1.00 | 1.00 | 1.00 | 1.00 | 1.00 | 1.00 | 1.00 | 0.99 | 1.00 | 1.00 | 1.00 | 1.00 | 1.00 | 1.00 | 1.00 | 1.00 | 1.00 | 1.00 |
| CP5 | 1.00 | 1.00 | 1.00 | 1.00 | 1.00 | 1.00 | 1.00 | 1.00 | 1.00 | 1.00 | 1.00 | 0.92 | 1.00 | 1.00 | 1.00 | 1.00 | 1.00 | 1.00 | 1.00 | 1.00 |
| CP3 | 1.00 | 1.00 | 1.00 | 1.00 | 0.89 | 0.82 | 1.00 | 1.00 | 0.91 | 0.77 | 1.00 | 0.96 | 1.00 | 1.00 | 1.00 | 1.00 | 1.00 | 1.00 | 1.00 | 1.00 |
| CP1 | 1.00 | 1.00 | 1.00 | 1.00 | 0.85 | 0.82 | 1.00 | 1.00 | 0.90 | 0.82 | 1.00 | 1.00 | 1.00 | 1.00 | 1.00 | 1.00 | 1.00 | 1.00 | 1.00 | 1.00 |
| CPz | 1.00 | 1.00 | 1.00 | 1.00 | 0.98 | 0.98 | 1.00 | 1.00 | 0.99 | 0.98 | 1.00 | 1.00 | 1.00 | 1.00 | 1.00 | 1.00 | 1.00 | 1.00 | 1.00 | 1.00 |
| CP2 | 1.00 | 1.00 | 1.00 | 1.00 | 1.00 | 0.95 | 1.00 | 1.00 | 1.00 | 0.94 | 1.00 | 1.00 | 1.00 | 1.00 | 1.00 | 1.00 | 1.00 | 1.00 | 1.00 | 1.00 |
| CP4 | 1.00 | 1.00 | 1.00 | 1.00 | 1.00 | 0.94 | 1.00 | 0.99 | 1.00 | 0.89 | 1.00 | 0.97 | 1.00 | 1.00 | 1.00 | 1.00 | 1.00 | 1.00 | 1.00 | 1.00 |
| CP6 | 1.00 | 1.00 | 1.00 | 1.00 | 1.00 | 1.00 | 1.00 | 1.00 | 1.00 | 1.00 | 1.00 | 1.00 | 1.00 | 1.00 | 1.00 | 1.00 | 1.00 | 1.00 | 1.00 | 1.00 |
| TP8 | 1.00 | 1.00 | 1.00 | 1.00 | 1.00 | 1.00 | 0.92 | 1.00 | 1.00 | 0.98 | 1.00 | 1.00 | 1.00 | 1.00 | 1.00 | 1.00 | 1.00 | 1.00 | 1.00 | 1.00 |
| P7 | 1.00 | 1.00 | 1.00 | 1.00 | 1.00 | 0.99 | 1.00 | 1.00 | 1.00 | 0.95 | 1.00 | 1.00 | 1.00 | 1.00 | 1.00 | 1.00 | 1.00 | 1.00 | 1.00 | 1.00 |
| P5 | 1.00 | 1.00 | 1.00 | 1.00 | 1.00 | 0.96 | 1.00 | 1.00 | 1.00 | 0.92 | 1.00 | 0.96 | 1.00 | 1.00 | 1.00 | 1.00 | 1.00 | 1.00 | 1.00 | 1.00 |
| P3 | 1.00 | 1.00 | 1.00 | 1.00 | 0.87 | 0.70 | 1.00 | 0.95 | 0.99 | 0.61 | 1.00 | 0.97 | 1.00 | 1.00 | 1.00 | 1.00 | 1.00 | 1.00 | 1.00 | 1.00 |
| P1 | 1.00 | 1.00 | 1.00 | 1.00 | 0.93 | 0.71 | 0.99 | 0.99 | 1.00 | 0.66 | 1.00 | 1.00 | 1.00 | 1.00 | 1.00 | 1.00 | 1.00 | 1.00 | 1.00 | 1.00 |
| Pz | 1.00 | 1.00 | 1.00 | 1.00 | 1.00 | 0.95 | 0.99 | 1.00 | 1.00 | 0.90 | 1.00 | 1.00 | 1.00 | 1.00 | 1.00 | 1.00 | 1.00 | 1.00 | 1.00 | 1.00 |
| P2 | 1.00 | 1.00 | 1.00 | 1.00 | 1.00 | 0.86 | 0.97 | 0.82 | 1.00 | 0.71 | 1.00 | 1.00 | 1.00 | 1.00 | 1.00 | 1.00 | 1.00 | 1.00 | 1.00 | 1.00 |
| P4 | 1.00 | 1.00 | 1.00 | 1.00 | 1.00 | 0.97 | 0.91 | 0.99 | 1.00 | 0.87 | 1.00 | 1.00 | 1.00 | 1.00 | 1.00 | 1.00 | 1.00 | 1.00 | 1.00 | 1.00 |
| P6 | 1.00 | 0.99 | 1.00 | 1.00 | 1.00 | 0.97 | 0.99 | 0.93 | 1.00 | 0.85 | 1.00 | 0.99 | 1.00 | 1.00 | 1.00 | 1.00 | 1.00 | 1.00 | 1.00 | 1.00 |
| P8 | 1.00 | 1.00 | 1.00 | 1.00 | 1.00 | 0.98 | 0.97 | 0.93 | 1.00 | 0.91 | 1.00 | 1.00 | 1.00 | 1.00 | 1.00 | 1.00 | 1.00 | 1.00 | 1.00 | 1.00 |
| PO7 | 0.86 | 0.99 | 1.00 | 1.00 | 0.78 | 0.79 | 1.00 | 0.98 | 0.90 | 0.74 | 1.00 | 1.00 | 1.00 | 1.00 | 1.00 | 1.00 | 1.00 | 1.00 | 1.00 | 1.00 |
| PO5 | 0.87 | 0.95 | 1.00 | 1.00 | 0.71 | 0.76 | 1.00 | 1.00 | 0.82 | 0.74 | 1.00 | 1.00 | 1.00 | 1.00 | 1.00 | 1.00 | 1.00 | 1.00 | 1.00 | 1.00 |
| PO3 | 0.96 | 0.96 | 1.00 | 1.00 | 0.88 | 0.56 | 1.00 | 0.72 | 0.98 | 0.49 | 1.00 | 1.00 | 1.00 | 1.00 | 1.00 | 1.00 | 1.00 | 1.00 | 1.00 | 1.00 |
| POz | 1.00 | 0.99 | 1.00 | 1.00 | 1.00 | 0.75 | 1.00 | 0.50 | 1.00 | 0.51 | 1.00 | 1.00 | 1.00 | 1.00 | 1.00 | 1.00 | 1.00 | 1.00 | 1.00 | 1.00 |
| PO4 | 1.00 | 0.99 | 1.00 | 1.00 | 1.00 | 0.71 | 0.71 | 0.39 | 1.00 | 0.42 | 1.00 | 0.99 | 1.00 | 1.00 | 1.00 | 1.00 | 1.00 | 1.00 | 1.00 | 1.00 |
| PO6 | 1.00 | 1.00 | 1.00 | 1.00 | 1.00 | 0.76 | 0.81 | 0.69 | 1.00 | 0.56 | 1.00 | 0.99 | 1.00 | 1.00 | 1.00 | 1.00 | 1.00 | 1.00 | 1.00 | 1.00 |
| PO8 | 1.00 | 1.00 | 1.00 | 1.00 | 1.00 | 0.77 | 1.00 | 0.87 | 1.00 | 0.66 | 1.00 | 1.00 | 1.00 | 1.00 | 1.00 | 1.00 | 1.00 | 1.00 | 1.00 | 1.00 |
| O1 | 0.90 | 1.00 | 1.00 | 1.00 | 0.69 | 0.72 | 1.00 | 0.95 | 0.75 | 0.68 | 1.00 | 1.00 | 1.00 | 1.00 | 1.00 | 1.00 | 1.00 | 1.00 | 1.00 | 1.00 |
| Oz | 1.00 | 0.25 | 0.99 | 0.29 | 1.00 | 0.82 | 1.00 | 0.74 | 1.00 | 0.65 | 1.00 | 0.95 | 1.00 | 1.00 | 1.00 | 1.00 | 1.00 | 1.00 | 1.00 | 1.00 |
| O2 | 1.00 | 1.00 | 1.00 | 1.00 | 0.98 | 0.79 | 1.00 | 0.75 | 1.00 | 0.65 | 1.00 | 1.00 | 1.00 | 1.00 | 1.00 | 1.00 | 1.00 | 1.00 | 1.00 | 1.00 |

Significant values ($p < 0.05$) are shown in bold. Abbreviations: AD = Antidepressants; AG = Analgesics.
